# A spatial atlas of the seaweed CO_2_-fixation machinery reveals a unique Rubisco condensation mechanism

**DOI:** 10.64898/2026.07.16.738900

**Authors:** James Barrett, Clément Dégut, Alejandro Sebiani Calvo, Michael R. Hodgkinson, Hannah Derry, Aidan Salter, Emma Coulembier Vandelannoote, Rebecca Beveridge, John H. Bothwell, Thomas B. Jacobs, Olivier De Clerck, James N. Blaza, Michael J. Plevin, Luke C.M. Mackinder, Jonas Blomme

## Abstract

Seaweeds (macroalgae) are important primary producers that sustain food webs in coastal ecosystems. Most algae accelerate inorganic carbon assimilation by actively concentrating CO_2_ in a Rubisco-rich specialized organelle called the pyrenoid. However, the molecular composition of this pathway is unknown in seaweeds. Here, we investigated the intracellular localization of 160 proteins associated with CO_2_ acquisition in the green seaweed *Ulva* (Sea lettuce). We assign 68 proteins to different pyrenoid subdomains and identify a consensus *Ulva* Rubisco binding motif revealing the molecular logic of the *Ulva* pyrenoid. We reveal Seaweed *Ulva* Pyrenoid Assembly 1 (SUPA1) as the core pyrenoid assembly factor. We show that Rubisco condensation is driven by a unique mechanism: the helical folding of SUPA1 motifs upon Rubisco binding, combined with steric hindrance that halves the available Rubisco binding sites from eight to four. Our data gives an unprecedented sub-cellular spatial understanding on seaweed carbon fixation and provides insights into the evolution of this important pathway in the global carbon cycle.

## Introduction

Seaweeds are a colorful, biodiverse and polyphyletic group of organisms. There are over 12,000 currently described seaweed species, most of which are spread across three evolutionarily distinct groups: the brown, green and red macroalgae. These groups are named after their dominant pigmentations and each group has independently evolved multicellularity^1,2^ so that all three groups contain species across a remarkable diversity of sizes and shapes.^3^ The seaweeds as a whole are critical primary producers and ecosystem engineers in coastal areas worldwide, and both the area and productivity of their biomes are comparable to those of the Amazon rainforest.^4^ Their ability to rapidly transform captured inorganic carbon (Ci) into growth potential is exemplified by frequent and extensive blooms that can lead to hugely damaging economic and environmental impacts.^5^ Conversely, harvested or cultivated seaweed biomass is often processed as food, feed or fertilizer, and holds promise as a source of pharmaceuticals and cosmetics.^6^

Despite this economic, evolutionary and environmental importance, the assimilation of Ci by seaweeds remains relatively poorly understood at the molecular level. Although the fixation of CO_2_ by Rubisco is conserved in all photosynthetic organisms, this carboxylation reaction is catalytically slow^7^ and Rubisco has a poor selectivity for CO_2_ over O_2_, which can reduce net carbon fixation by up to 50%.^8^ Mechanisms have therefore evolved to concentrate CO_2_ in the vicinity of Rubisco to favor the carboxylation reaction^9,10^ and accelerate biomass accumulation.

In the majority of marine algae, this Ci concentration takes place in a specialized chloroplast organelle called the pyrenoid, and pyrenoid-based CO_2_-concentrating mechanisms (pCCM)^9,11^ have been estimated to contribute ∼30% to global photosynthetic carbon capture.^12^ Pyrenoids are biomolecular condensates consisting of Rubisco and other proteins that mediate organelle formation, structure and maintenance.^13,14^ Under limiting CO_2_ conditions, the pCCM functions by actively assimilating bicarbonate (HCO_3_^-^) from (sea)water and ultimately dehydrating it to CO_2_ by a carbonic anhydrase (CA) enzyme close to Rubisco.^15^ The molecular pCCM components have been characterized to varying extents in a small number of microalgae and hornworts.^16–20^ In the freshwater microalga *Chlamydomonas reinhardtii* (*Chlamydomonas* hereafter), for example, high-throughput protein-tagging experiments have provided extensive insights into the pCCM. By mapping molecular components to subcellular regions, these studies revealed a spatially organized pathway in which HCO_3_^-^ is transported across the plasma membrane, chloroplast envelope and thylakoid membranes to the pyrenoid composed of multiple functionally-specialized subdomains.^16,21,22^ These studies, and localization of components in other algal systems, underscore that the precise subcellular localization of core pCCM transporters, enzymes and structural components is critical for pathway function.^16,18,20,21^

However, data indicate that pyrenoids have convergently evolved across different algal clades and may therefore display structural and molecular diversity.^11,17–19^ For example, in the core chlorophyte lineages of the Chlorophyceae and Trebouxiophyceae, multivalent disordered linker proteins have independently evolved to bind and condense Rubisco to form the pyrenoid matrix.^12,19^ The interaction of linkers and other structural proteins with Rubisco is mediated by small Rubisco-Binding Motifs (RBMs), which bind to the small or large Rubisco subunit (RbcS or RbcL) in *Chlamydomonas* and *Chlorella*, respectively.^19,22^

Consequently, determining the spatial organization of the pCCM is a crucial first step to understanding how the principles of Ci assimilation work in larger, multicellular, marine algae. The scale of such experiments necessitates *bona fide* genetic model systems that, to date, have only been available in the unicellular freshwater chlorophyte *Chlamydomonas*^16,21^ and, to a lesser extent, the unicellular diatom *Thalassiosira pseudonana*.^23^ Although recent advances in macroalgal functional biology have yielded genetic models for *Ectocarpus*, *Neopyropia* and *Ulva* that support stable transformation and targeted mutagenesis^24–29^, high-throughput gene manipulation remains to be demonstrated in these models.^30,31^

In this study, we rectify this omission to provide a spatial protein blueprint of the carbon capture pathway in a species from the green seaweed genus *Ulva* (Sea lettuce), a much-studied exemplar for marine macroalgal photosynthesis.^32–34^ *Ulva* is a globally distributed genus that occurs primarily in marine, but also in brackish and even freshwater habitats.^35–37^ *Ulva* species are morphologically plastic but typically comprise of a single layer of photosynthesizing blade cells and a small holdfast for attachment (**Fig. 1A**).^38–40^ *Ulva* species feature the world’s highest photosynthetic rates per area^4,33^ and opportunistic *Ulva* blooms, known as green tides, are increasingly perturbing coastal ecosystems worldwide, with extremely detrimental environmental and economic impacts^5^. More positively, *Ulva* biomass is also exploited in aquaculture applications.^41,42^ Either way, *Ulva*’s prolific growth is associated with an efficient conversion of Ci and nutrients into biomass^43,44^, driven by a highly efficient pCCM that has been confirmed by physiological studies^34,45–47^, but whose molecular components remain unexplored and undescribed.

**Figure 1.**
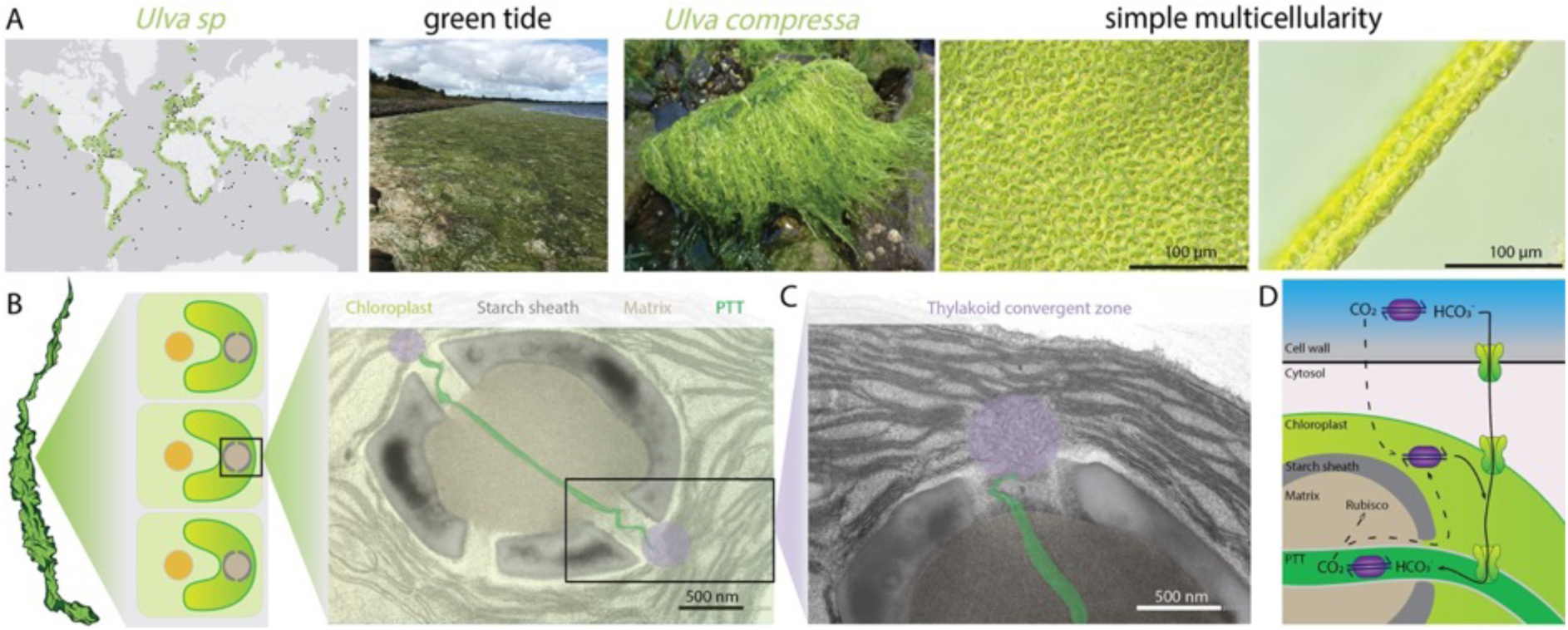
*Ulva* is a green seaweed with a pyrenoid-based CO_2_ concentrating mechanism. A. *Ulva* is globally distributed and ecologically relevant. First panel: Ocean Biodiversity Information System (OBIS) distribution data. Each black dot on the map represents the recorded presence of *Ulva* or its synonym *Enteromorpha* at genus level; main coastlines are highlighted in green. Second panel: *Ulva* green tide in Frederikshavn, Denmark (Picture courtesy of Sophie Steinhagen). Third panel: *Ulva compressa* is a common species in coastal ecosystems. Picture from Neeltje Jans, The Netherlands (Picture courtesy of Luna van der Loos). Fourth and fifth panel: Light microscope pictures illustrate that *Ulva* is a simple multicellular organism. Each cell contains a single cup-shaped chloroplast. Scale: 100 µm. B. TEM image of *Ulva* chloroplast containing a single pyrenoid, surrounded by a starch sheath and a pyrenoid traversing thylakoid (PTT). Also see **Fig. S1**. Scale: 500 nm. C. Stromal thylakoid membranes undergo structural changes and fuse to form the PTT in a thylakoid convergent zone. Also see **Fig. S1**. Scale: 500 nm. D. The putative inorganic carbon uptake pathway components based on the *Chlamydomonas reinhardtii* pCCM that shuttle CO_2_/HCO_3_^-^ from the surrounding environment to Rubisco in the pyrenoid. Carbonic anhydrases (purple) mediate the reversible interconversion of CO_2_ into HCO_3_^-^. Inorganic carbon transporters (green) at plasma membrane, chloroplast envelope and thylakoid membrane move HCO_3_^-^ from the environment to the PTT.

In this study we leverage our recently developed genetic tools in *Ulva compressa/mutabilis* (*Ulva* hereafter) to fluorescently tag 201 proteins associated with the pCCM, providing an unprecedented first spatial understanding of a seaweed pCCM. Our microscopy data reveal core components of the carbon fixation pathway, identifying 68 proteins that localize to subdomains of the pyrenoid, of which 15 are exclusively localized to this condensate. We identify the core pyrenoid assembly protein, which we name Seaweed *Ulva* Pyrenoid Assembly protein 1 (SUPA1). Using a combination of *in vitro*, *in vivo* and structural studies we show that SUPA1 is required for Rubisco condensation to form the pyrenoid by binding to both the small and large Rubisco subunits at four locations per holoenzyme. Our study is a first demonstration that high-throughput screens are possible in macroalgae and provides a strong foundation for the molecular understanding of carbon capture in an ecologically and economically important organism.

## Results + discussion

### *Ulva* has a distinct pyrenoid ultrastructure

In agreement with previous studies^48–50^ TEM microscopy showed that *Ulva* cells contain a large chloroplast that occupies ∼35% of the cell area (**Fig. S1**). Within each chloroplast is a single pyrenoid encapsulated by a starch sheath that was typically formed from two, but sometimes three, starch plates. In all TEMs with a central pyrenoid mid-section we observed a single traversing membrane continuous with the thylakoid network. This suggests that the pyrenoid traversing thylakoids (PTTs) in *Ulva* consist of a single thylakoid membrane sheet (**Fig. 1B-C; Fig. S1**), in contrast to the chlorophyte *Chlamydomonas*, which contains multiple tubular PTTs fusing at the center of the pyrenoid^51^, and the trebouxiophyte *Chlorella*, which has a double-layered sheet-like PTT.^52^ Additionally, *Ulva* displays a large thylakoid structural change adjacent to the PTT entry sites on both sides of the pyrenoid. In these regions, multiple (20-45, n=11) stromal thylakoids converge, transitioning from typical flat ordered stacks into a disordered network where they appear to fuse into a single swollen thylakoid that then enters the pyrenoid matrix (**Fig. 1B-C; Fig. S1**). We term these regions thylakoid convergent zones and propose that they enable the connection of a large volume of the thylakoid lumen to the PTT.

### Identification of *Ulva* pCCM-related genes

We reasoned that an orthology-based approach would be the quickest way to identify an initial set of proteins putatively involved in the *Ulva* pCCM. We therefore turned to the well-characterized pCCM of the single-celled model green alga *Chlamydomonas*^16,21,53^, which belongs to the Chlorophyceae, a class that diverged from the Ulvophyceae in the Mesoproterozoic or early Neoproterozoic (1300-800 Mya).^54,55^ Components of the *Chlamydomonas* pCCM can be broadly split into three categories: 1) Ci transport uptake components that shuttle CO_2_/HCO_3_^-^ from the surrounding environment to the pyrenoid (**Fig. 1D**); 2) pyrenoid structural components; and 3) regulatory components that control pCCM induction and pyrenoid assembly. In addition to these core components, many further *Chlamydomonas* genes have been associated with its pCCM in large-scale gene-tagging, transcriptomic and/or proteomic studies, and through detailed biochemical characterization. However, the exact function of many of these genes remains to be demonstrated.^13,14^ Recent advances in the understanding of pCCM components in the Trebouxiophyceae (split from *Ulva* >850 Mya^19,55^) and hornworts (split from *Ulva* >1,1 Gya^20,56,57^) suggested that some *Ulva* proteins responsible for Ci uptake and/or pyrenoid assembly would be conserved with *Chlamydomonas* while others would not.

We therefore narrowed down a list of 610 *Chlamydomonas* pCCM-related genes that includes known *Chlamydomonas* pCCM components, pyrenoid-related genes and genes that are differentially expressed under low CO_2_ conditions^16^ by excluding genes with a less well-established link to the pCCM pathway (“other” category in Mackinder *et al.*^16^). We supplemented this list with organelle markers (chloroplast, ER, Golgi, mitochondria, nucleus, peroxisomes, plasma membrane and vacuole)^16^ and more recently identified pyrenoid-localized proteins in *Chlamydomonas*.^21,53^ From the resulting 343 pCCM-associated *Chlamydomonas* genes we determined orthologous gene relationships using the PLAZA platform that integrates support from different methods.^58^ We did not identify an *Ulva* ortholog for 130 genes, but we were able to identify a single ortholog for 156 genes (151 unique genes) and two or more orthologs for the remaining 57 genes (156 unique genes; **Fig. 2A**; **Table S1; Table S2; Table S3**). In addition, we identified five putative *Ulva* pyrenoid proteins based on physicochemical properties^19^, but not sequence conservation, to known pyrenoid proteins (see below; **Fig. 2A**). Taking all approaches together, we identified 309 unique genes including 33 organelle markers and 276 putative pCCM-associated genes in *Ulva*.

**Figure 2.**
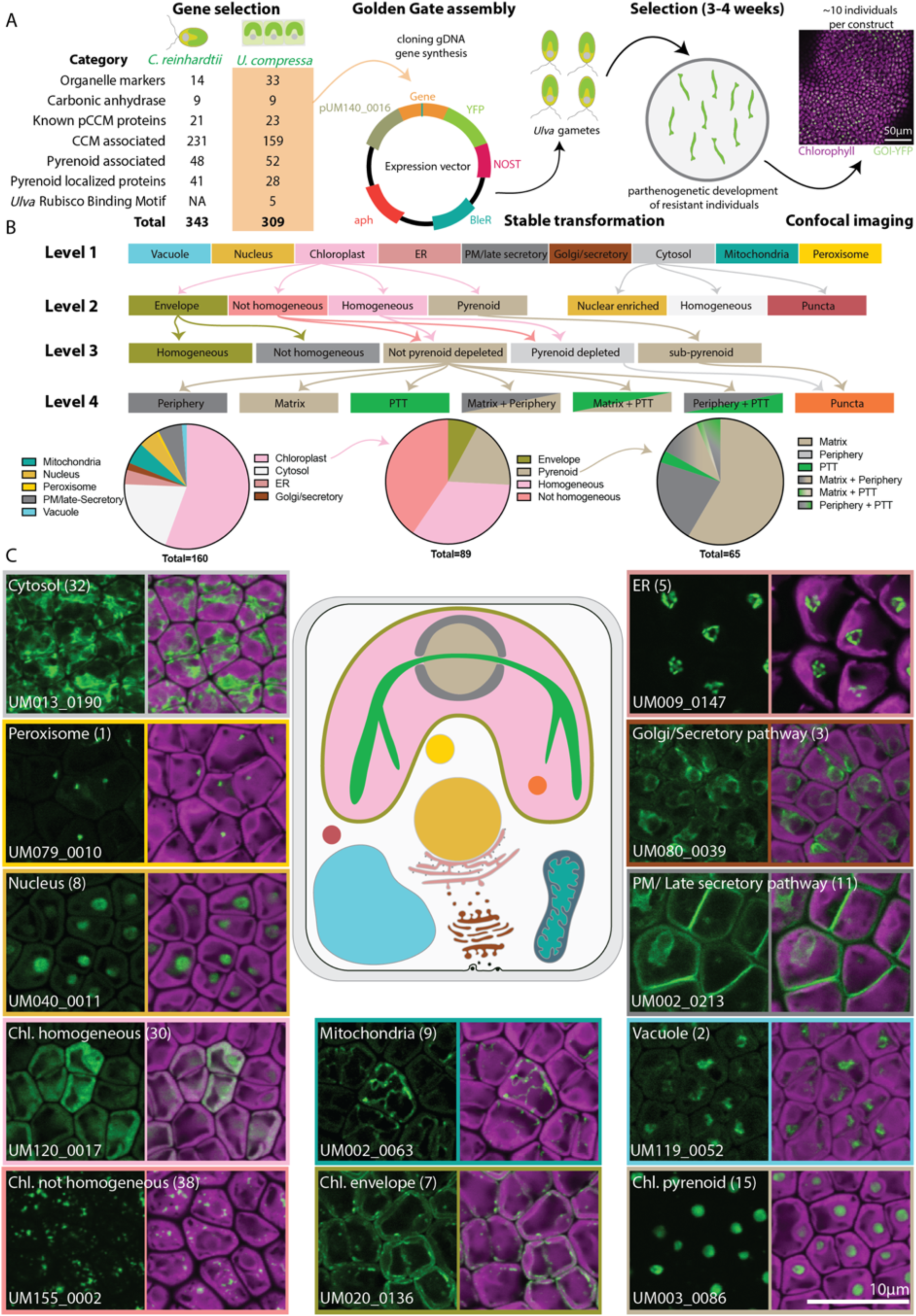
pCCM localization overview. A. Illustration of cloning and imaging pipeline. Sequence orthology reveals pCCM- related genes in *Ulva*. *Chlamydomonas* genes were identified in earlier studies.^15,23,57^ Total number of orthologous *Ulva* genes per category are indicated. See also **Table S2**; **Table S3**. All genes are assembled in an expression vector under control of a strong endogenous promoter (pUM140_0016), coupled to a YFP sequence and a terminator (NOST). The expression vector also contains an antibiotic resistance cassette for cloning (aph) and *Ulva* selection (BleR). See also **Table S7**). Gametes are stably transformed with the generated vectors, after 3-4 weeks the transformants are imaged using confocal microscopy. Magenta: Chlorophyll autofluorescence; Green: YFP signal of tagged protein. Scale: 50 µm. B. Classification of observed localizations in different levels of detail. Protein localization decision tree to assign proteins to specific subcellular locations. Pie charts visualize the number of proteins per level, with a focus on sub-chloroplast and sub-pyrenoid localization for sublevel 2, 3 and 4. C. Illustration of observed intracellular localizations. For all localizations, the number of proteins associated with the cellular sub compartment is indicated. Magenta: Chlorophyll autofluorescence; Green: YFP signal of tagged protein. Scale: 10 µm. Also see **Fig. S3** and **Table S2**.

### Large-scale fluorescent tagging assigns 160 proteins to 34 subcellular locations

We next aimed to leverage stable transformation^25,59^ to get a comprehensive and spatially-resolved view of the molecular components of the *Ulva* pCCM that determine its architecture and function (**Fig. 1B-D**). Accordingly, we fluorescently tagged the gDNA sequence from 201 of 309 genes (65%; **Table S2**), with cloning success dependent on a combination of factors including gene size, GC content and FPKM (**Fig. S2A-C**). All genes were fused to a C-terminal Yellow Fluorescent Protein (YFP) sequence via Golden Gate assembly in an expression vector, under control of a constitutive promoter and the nopaline synthase terminator (NOST) (**Fig. 2A**). Expression vectors were stably transformed into *Ulva* gametes. After three weeks of selection, 3-29 (average 10) independent stably transformed individuals were imaged per construct (**Fig. 2A**, **Table S2**).

This allowed unambiguous intracellular localization of 160 tagged proteins (**Fig. S3**). Except for two proteins, we obtained localization data in at least two, on average six, independent transformants (**Table S2**). Our success rate of 80% of constructs yielding protein localization data is high compared to 33-49% in *Chlamydomonas* in similar screens.^16,21^ This indicates that transgene silencing^60^ may be less of a challenge in *Ulva*, further supported by 57% of transformants per construct expressing the fusion protein compared to <15% in *Chlamydomonas* (**Table S2; Fig. S2A-H**).^61^ To better understand endogenous abundances of pCCM-related proteins and to determine if they impacted localization success rates, we integrated available RNA expression data^62^ and generated whole cell proteomic datasets of *Ulva* (**Table S2; Table S4**). Overall, localization success is not correlated with RNA expression level, unlike *Chlamydomonas*^16^, but is with higher protein abundance (**Fig. S2A-L**).

Since this is the first protein-tagging study of this scale in a seaweed, we used the localization of marker proteins to guide the classification of unknown proteins. Marker proteins were selected based on established organelle and sub-organelle localization from orthology or previously generated localization data.^25,39,59,63^ Guided by a decision tree that integrated marker protein data and image features, we classified the 160 localized proteins into 34 subcellular regions (**Fig. 2B; Table S2**). Despite our reference gene set and success in transgene expression, artefacts or incomplete observations cannot be excluded since we used a constitutive promoter and we imaged at a single developmental time point. For example, flagella are absent during vegetative development in *Ulva* so localization to this cellular structure was not evaluated. For some proteins a specific sub-cytosol or secretory pathway localization was challenging to classify. Additional localization details or observations from alternative independent lines were recorded where relevant (**Table S2**).

At the whole-cell level, we were able to distinguish nine major intracellular locations (**Fig. 2B-C; Fig. S3**). Most proteins were localized to the chloroplast (89), cytosol (32) or secretory pathway (ER, Golgi and/or PM; 19). We also identified proteins with mitochondrial (9) or nuclear (8) localization and proteins targeted to the vacuole (2) or peroxisome (1; **Fig. 2B-C**). 32 proteins were localized simultaneously to two cellular compartments. The majority of these proteins appeared to be concurrently localized to different organelles of the secretory pathway (ER, Golgi and/or PM; 12; *e.g.* UM080_0043), cytosol and PM (10, *e.g.* UM022_0185) or chloroplast and cytosol (6; *e.g.* UM002_0253; **Table S2; Fig. S3**). We compared the observed localizations with the predictions generated by three models: PB-Chlamy^21^, PredAlgo^64^ and TargetP^65,66^. These use different algorithms and have been trained on different datasets: TargetP is broadly trained on eukaryotic protein sequences with experimentally verified localizations, while PredAlgo and PB-Chlamy are trained on *Chlamydomonas* protein localization datasets. The algorithms differed in their false positive and false negative rates, but PB-Chlamy provided the most accurate prediction for *Ulva* protein localization. Predicting 99% of chloroplast localized proteins correctly, 70% for mitochondrial localized proteins and 67% for secreted proteins (**Fig. S4**).

We further subgrouped cytosolic and chloroplast proteins (**Fig. 2B-C**). Most cytosolic proteins were nuclear enriched (17) or homogeneously distributed (13), but we also observed cytosolic puncta for the putative peroxiredoxin UM012_0167. Chloroplast proteins were distributed homogeneously (30), non-homogeneously (38) or exclusively to either the pyrenoid (15) or chloroplast envelope (7; **Fig. 2B-C; Table S2**). Chloroplast proteins were further classified according to presence, depletion or enrichment in pyrenoid subdomains (**Fig. 2B-C; Table S2**). We evaluated if *Ulva* could provide novel or alternative insights into protein localization compared to *Chlamydomonas*. We focused on the 151 *Ulva* genes with a single ortholog in *Chlamydomonas*. For the 57 proteins with a localization in both organisms, 21 proteins showed the same spatial pattern, 23 were similar and for 13 the localization was different (**Table S3**). The similar category includes proteins with variations in sub-organelle localization and/or proteins that were localized in two cellular compartments in one organism but a single compartment in the other. Interestingly, 14 proteins with a similar localization pattern (61%) had a different sub-pyrenoid enrichment (see below, **Table S3**).

In summary, we used and optimized available molecular tools to enable protein-tagging screens at scale in a seaweed, which reveals diverse localizations. The enrichment for chloroplast-localized proteins enabled the discerning of fine sub-chloroplast features. We observe high expression in many transformed individuals, illustrating that *Ulva* represents an excellent model for functional biology in green algae. Broadly, orthologous *Ulva* and *Chlamydomonas* proteins localize to the same or similar locations, further suggesting that many pCCM components operate at similar spatial scales in chlorophytes.

### Tagging reveals the spatial organization of inorganic carbon delivery to Rubisco

Pyrenoid-based biophysical CCMs function by accumulating CO_2_ around Rubisco at higher concentrations than the surrounding environment. This is achieved by an interplay of Ci transport across membranes, the CA-catalyzed interconversion of CO_2_ and HCO_3_^-^, and compartmentalized pH to drive either the accumulation of membrane-impermeable HCO_3_^-^ or the release of CO_2_ for fixation by Rubisco (**Fig. 1D**; **Fig. 3**).^14,67^ The majority of genes involved in Ci delivery in *Chlamydomonas* are functionally characterized.^68–73^ We found *Ulva* orthologs for most core *Chlamydomonas* Ci delivery proteins, including HCO_3_^-^ transporters (HLA3, LCIA), HCO_3_^-^ channels (bestrophins), external CAs (CAH1/2), stromal CAs (LCIB/C) and the thylakoid CA (CAH3). By integrating our expression, localization and/or orthology data we can functionally assign *Ulva* proteins to the core Ci assimilation steps.

**Figure 3.**
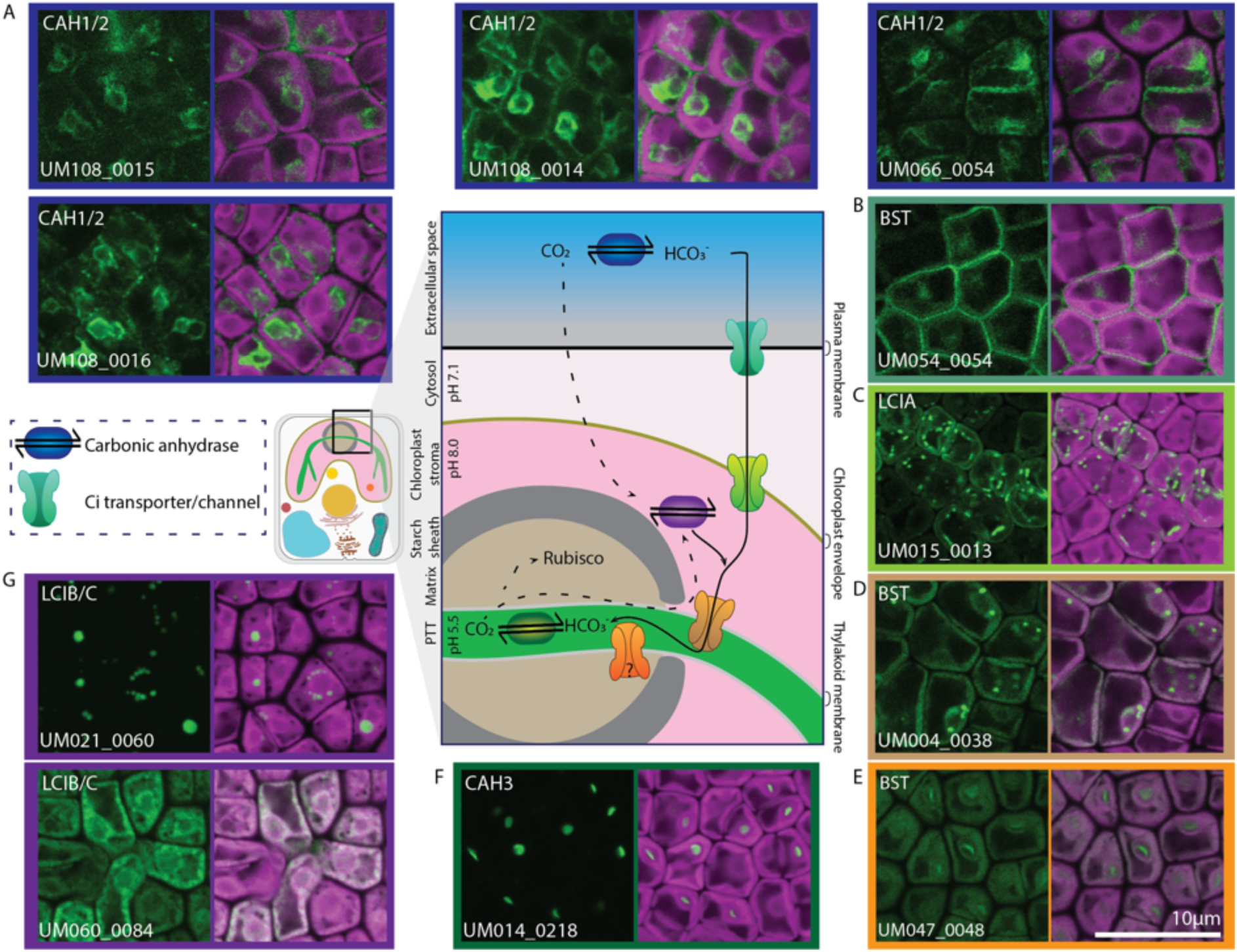
Tagging reveals spatial organization of inorganic carbon transport to Rubisco. Candidate inorganic carbon transporters and carbonic anhydrases are identified at different cellular localizations. A. Secreted carbonic anhydrases are CAH1/2 orthologs. B. Bestrophin (BST) is a putative Ci transporter at the plasma membrane. C. LCIA localizes to the chloroplast envelope. D. BST protein localizes to thylakoid convergence zones at the pyrenoid periphery. E. BST protein is specific to the PTT. F. CAH3 is specific to the PTT. G. LCIB/C localizes to the chloroplast stroma. Magenta: Chlorophyll autofluorescence; Green: YFP signal of tagged protein. Scale: 10 µm. See also **Fig. S3** and **Table S2**.

#### Inorganic carbon delivery to the chloroplast

Ci assimilation from the surrounding environment proceeds via diffusion across the boundary layer and cell wall to the plasma membrane. As *Ulva* is mostly found in the intertidal region, the available Ci species differ between immersion and emersion. When immersed in seawater under natural conditions (pH 8.1), dissolved HCO_3_^-^ is the major Ci molecule. In contrast, atmospheric CO^2^ is more abundant when *Ulva* is emersed.^74,75^ The capacity to continue photosynthesis and utilize two different Ci sources in two drastically different environments twice every day underpins the ability of *Ulva* to thrive in coastal ecosystems.^34,76,77^

Several CA families, denoted by Greek letters, are pCCM-associated.^78,79^ The *Ulva* genome encodes twelve CAs: six alpha-type CAs, three beta-type CAs and three gamma-type CAs. In eukaryotic algae, including *Ulva*, the gamma-type CAs are mitochondrial and their activity or role in pCCM is not clearly established; whereas alpha- and beta-type CAs play central roles at multiple cellular locations. In *Chlamydomonas* two periplasmic alpha-CAs, CAH1 and CAH2, catalyze the interconversion of CO_2_ with HCO_3_^-^.^73,80^ Early data using acetazolamide, a membrane impermeable CA inhibitor, suppressed photosynthesis supporting a similar role for external CAs in *Ulva* photosynthesis.^46,81^ Five alpha-type CAs are orthologs of *Chlamydomonas* CAH1/2, including three paralogs UM108_0014, UM108_0015 and UM108_0016 that share >98% protein sequence identity. These duplicated genes are lowly expressed and not identified in whole-cell MS, but we localized the proteins and an additional alpha-type CA (UM066_0054) to the secretory pathway and plasma membrane (**Fig. 3A**), with similar localization patterning to secreted proteins in *Chlamydomonas*.^16^ The fifth alpha-type CA, UM028_0029, is expressed and detected in MS data and also predicted to be secreted, but we did not observe YFP signal in the transformants. Based on these data, *Ulva* has at least two and up to five alpha-CAs expressed at the plasma membrane.

Depending on Ci availability and Ci species (e.g. CO_2_ vs HCO_3_^-^), uptake across the plasma membrane can be passive via CO_2_ diffusion or involve active HCO_3_- uptake. Three different transporter families could mediate the initial Ci uptake at the plasma membrane in *Ulva*. Inhibitor based studies and cross-reactivity of a plasma membrane protein with an antibody to the SLC4 family human red-blood-cell chloride/HCO_3_^-^ exchanger (AE1/SLC4A1) suggests that Ci uptake at the *Ulva* plasma membrane probably involves at least one anion exchanger.^82,83^ We identified two proteins (UM066_0035 and UM066_0036) that are predicted to be secreted and have sequence similarity to SLC4A1. SLC4 family transporters are associated with HCO_3_^-^ uptake in marine diatoms^84^ but are not involved in the freshwater *Chlamydomonas* pCCM. Instead, in *Chlamydomonas* two different transporters mediate Ci uptake across the plasma membrane: HLA3 and LCI1.^70,85^ The *Ulva* genome does not encode an LCI1 ortholog but we identified five putative HLA3 proteins that are predicted to be secreted. We were unable to clone these putative HCO_3_^-^ transporters, but UM139_0019 is likely the most important HLA3 given its transcript levels^62^ and abundance in our whole-cell proteome data (**Table S2; Table S4**). Bestrophins (BSTs) have also been identified as widely distributed pCCM-associated HCO_3_^-^ channels, with data supporting that they mediate HCO_3_^-^ transport across thylakoid membranes in green algae, hornworts and diatoms.^18,20,71^ Based on sequence similarity, we identified thirteen putative *Ulva* BSTs: five that are similar to thylakoid localized CrBST1-3 and eight that have closer sequence similarity with Cre11.g467611 that has no known CCM function. We cloned six and obtained localization data for four putative BSTs. Two localized to the chloroplast (see following section) and one to the plasma membrane (UM054_0054, **Fig. 3B**). UM022_0150 is predicted to be secreted and we observed signal that was not confined to an intracellular structure but found throughout the cell. We therefore cannot unambiguously assign a localization for this protein (**Fig. S3**). For the fifth BST (UM054_0055), that is also predicted to be secreted, there was no detectable YFP fluorescence. Although we did not detect these BSTs in the whole-cell proteome and their function needs to be confirmed, this is to our knowledge the first demonstration of an algal bestrophin, UM054_0054, that localizes to the plasma membrane.

After transport across the plasma membrane Ci needs to move across the chloroplast envelope into the chloroplast stroma. We identified a single ortholog (UM015_0013) of the HCO_3_^-^ channel LCIA.^72,86^ UM015_0013 localizes to the chloroplast envelope homogeneously but also in puncta enriched on the internal, cytoplasmic facing region of the chloroplast envelope (**Fig. 3C**). Taken together, our data provides an overview of inorganic carbon transport from the surrounding environment to the chloroplast in *Ulva*.

#### Inorganic carbon delivery to the pyrenoid and CO_2_-recapture

In all characterized pCCMs to date, stromal accumulated HCO_3_^-^ is proposed to be shuttled by BSTs into the lumen of thylakoids that traverse the pyrenoid. Our screen identified two pyrenoid-associated BSTs. UM004_0038 typically forms two clear puncta at the pyrenoid periphery, which align with the thylakoid convergent zones we see in our TEMs that potentially create a large thylakoid surface area for HCO_3_^-^ channeling into the PTT (**Fig. 1C- D**; **Fig. 3D**). UM047_0048 localizes to the PTT itself, forming a clear single band across the pyrenoid. This is a similar localization to *Chlamydomonas* CrBST4, which is associated with maintaining ion homeostasis in the thylakoid lumen, although its substrate is still unclear (**Fig. 3E**).^87^ In *Chlamydomonas* CrBST4 is targeted to the PTTs via an extended disordered C-terminal that contains multiple RBMs shared with the Rubisco condensing protein CrEPYC1.^87^ UM047_0048 also possesses a predicted disordered extension at the N-terminus (**Fig. S5**; see next section).

The low pH of the thylakoid lumen drives the conversion of HCO_3_^-^ to CO_2_, a process accelerated by a specialized CA enriched in the PTTs^16–18,20,88^, with the alpha carbonic anhydrase 3 (CAH3) performing this role in *Chlamydomonas*.^89^ We found a single ortholog of CrCAH3, UM014_0218, which was exclusively localized to the PTT (**Fig. 3F**). An alpha-type CA is also localized to the PTTs in hornworts^90^ supporting that the final HCO_3_^-^ to CO_2_ dehydration step is conserved across pCCMs in the Viridiplantae.

Released CO_2_ is then fixed by Rubisco in the pyrenoid. Integral to stromal HCO_3_^-^ delivery and pCCM efficiency in *Chlamydomonas* are the stromal beta-type CAs, LCIB and LCIC, that have a dual role of converting CO_2_ into HCO_3_^-^ (**Fig. 1D**):^68,91,92^ LCIB and LCIC form a dynamic complex that convert inward fluxing CO_2_ into HCO_3_^-^ at atmospheric CO_2_ levels but relocalize under severely limiting CO_2_ levels to the pyrenoid periphery where they are associated with converting CO_2_ leaking from the pyrenoid back into HCO_3_^-^.^68,91,92^ We identified three putative LCIB/C orthologs and localized two to the chloroplast: UM021_0060 formed various sized puncta associated with the pyrenoid periphery whilst UM060_0084 was diffuse throughout the stroma (**Fig. 3G**).

In conclusion, the spatial organization of inorganic carbon transport to the pyrenoid is largely conserved in green algae. Our data suggests a broader repertoire of CO_2_ and HCO_3_^-^ interconversion and transport proteins at the plasma membrane in *Ulva*. Although the precise involvement of the different components needs to be validated, we hypothesize that this extended set of components facilitates the frequent shift in environmental Ci availability in an intertidal environment.

#### Core Chlamydomonas pyrenoid structural components are absent in Ulva

The pyrenoid has been observed since the first microscopic observations of *Ulva*, when this structure was proposed to function in starch synthesis due to the surrounding starch sheath.^50,93–95^ Later, ultrastructural studies visualized the presence of starch plates around the pyrenoid and a single traversing thylakoid (**Fig. 1B**; **Fig. S1**).^48^ Here we provide insight into the molecular components that form this complex biomolecular condensate. In total, 68 of our pCCM target proteins were found to localize to the pyrenoid, with 15 proteins exhibiting exclusive localization to a pyrenoid subdomain. Additionally, 25 proteins were associated with the matrix, 18 with the periphery, one with the PTT, and 9 with a combination of sub-pyrenoid regions (**Fig. 2B**; **Fig. 4; Table S2**, see later section).

**Figure 4.**
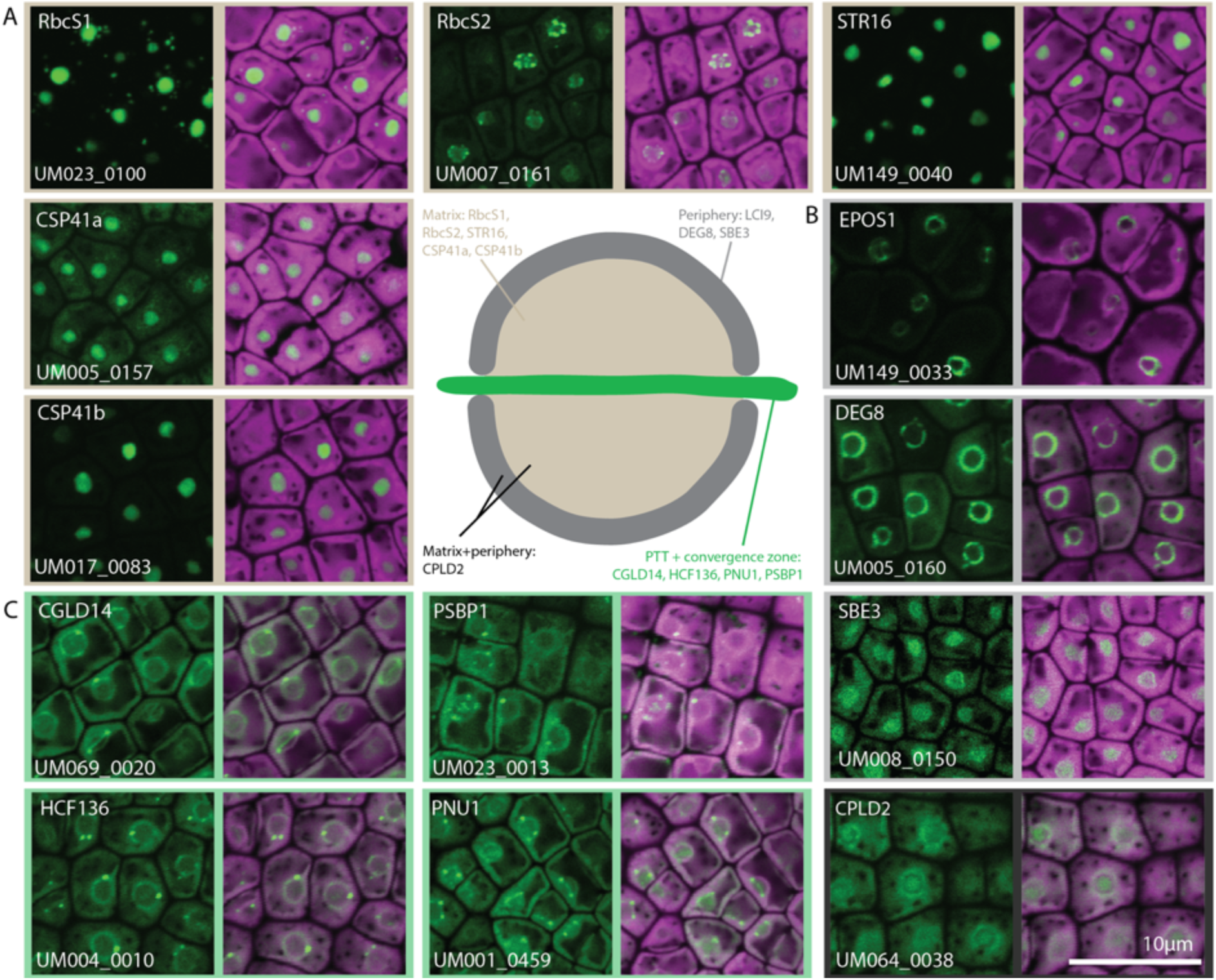
The pyrenoid has multiple subdomains. Although core structural proteins of *Chlamydomonas* pyrenoids are absent in *Ulva*, some orthologs of pyrenoid-exclusive proteins are associated with one or more pyrenoid subdomains (A) matrix, (B) periphery and/or (C) PTT) in *Ulva*. Magenta: Chlorophyll autofluorescence; Green: YFP signal of tagged protein. Scale: 10 µm. Also see **Table S5** and **Fig. S6**.

The pCCM functions to concentrate CO_2_ at the active site of Rubisco, a hexadecameric enzyme composed of 8 large subunits (RbcL) and 8 small subunits (RbcS). Whilst the single *Ulva* RbcL gene is chloroplast genome encoded; there are two nuclear encoded RbcS genes. RbcS1 (UM023_0100) and RbcS2 (UM007_0161) both localized to the pyrenoid matrix but also formed puncta but with subtle differences. RbcS1 was found throughout the matrix in addition to small puncta adjacent to the pyrenoid but external to the starch sheath. In contrast, RbcS2 formed puncta internal to the starch sheath and predominantly at the periphery of the matrix (**Fig. 4A; Fig. S6A**). Interestingly, RbcS2 is annotated as containing a C-terminus disordered extension. The recent identification that an RbcS extension drives pyrenoid assembly in hornworts^57^ suggested that this protein could have a similar role in *Ulva*. However, further analysis of available RNA-seq data indicates that RbcS2 is lowly expressed (∼3% compared to RbcS1) and relatively few reads map to the third and fourth exon of the gene (**Fig. S6B**) suggesting that either the gene model needs refinement, or that it undergoes alternative splicing that would result in a non-functional RbcS. Additionally, we see no corresponding peptides in our whole cell mass spectrometry data to the central region and the protein appears at a relatively low abundance (**Fig. S6C**), with high protein abundance being a prerequisite for a Rubisco condensing, pyrenoid assembly protein.^11^

Surprisingly, based on sequence conservation, no orthologs were identified for most genes involved in pyrenoid assembly in *Chlamydomonas* (**Table S5**). This includes the core structural pyrenoid components Essential Pyrenoid Component 1 (EPYC1), which condenses Rubisco to form the pyrenoid matrix; StArch Granules Abnormal 1 and 2 (SAGA1-2), which are associated with pyrenoid starch assembly and PTT formation^12,22,96,97^; Rubisco-binding membrane protein 2 (RBMP2), which regulates the formation of a reticulated region of the PTT^98^; and MIssing THylakoids 1 (MITH1), which is required for PTT formation.^97^ From the well-characterized proteins involved in pyrenoid assembly, only the ortholog of Element for Pyrenoid Organization of Starch 1 (EPOS1; UM149_0033) was identified, which also localized exclusively to the pyrenoid periphery (**Fig. 4B**). CrEPOS1 contains two CBM20 domains and forms a mesh-like structure around the pyrenoid in *Chlamydomonas*, where it mediates the formation of the starch sheath.^16,99^ We therefore next sought to understand how the *Ulva* pyrenoid is structurally assembled.

#### Identification of the Rubisco-condensing Ulva pyrenoid assembly protein

Our orthology-based approach failed to identify a candidate Rubisco-condensing protein with sequence similarity to EPYC1 or CsLinker (**Table S5**). Whilst a RbcS with a C-terminal extension was found in the genome, the lack of similarity to the self-condensing RbcS in hornworts^57^ and low transcriptional and protein abundance indicates that it is unlikely to drive *Ulva* pyrenoid assembly (see above; **Fig. S6**). With this, we hypothesized that *Ulva*, like *Chlamydomonas* and *Chlorella*, would possess an abundant intrinsically disordered repeat protein that drives Rubisco condensation to form the pyrenoid.

We used the sequence-homology-independent FLIPPer search pipeline^19^ to identify 8 proteins in the *Ulva* genome with the required features to perform Rubisco cross-linking: ≥3 repeats, largely disordered with a pI>8 (**Fig. S7**). Whole-cell mass spectrometry (MS) has demonstrated that the *Chlamydomonas* and *Chlorella* Rubisco-linker proteins are both abundant^19,100^ and, in agreement with *in vivo* quantification, *in vitro* condensation indicates that approximately equimolar concentrations of linker:Rubisco holoenzyme are required for efficient condensation.^19,101^ To determine the relative abundance of FLIPPer-identified candidates we performed MS on whole cells and on a crude pyrenoid enrichment. In both experiments we only detected one FLIPPer candidate, UM120_0037. In the whole-cell MS it was the 54th most abundant of 2895 detected proteins (**Fig. 5A-D, Table S4**). Whilst pyrenoid enrichment MS was contaminated with other organelles (nucleus) and large protein complexes (ribosomes), UM120_0037 was strongly enriched along with other pyrenoid components (*e.g.* RbcS, RCA and CPN60). We therefore proceeded with the characterization of UM120_0037 as a putative linker protein in *Ulva* and named it Seaweed *Ulva* Pyrenoid Assembly 1 (SUPA1). SUPA1 comprises 291 residues with 6 repeating sequences (R1-R6) which each contain a positively-charged predicted alpha-helical region we hypothesize are *Ulva*-specific RBMs. The helices are separated by low complexity, predicted disordered spacer sequences of ∼40 residues, except for the first spacer which is ∼20 residues (**Fig. 5E,F**).

**Figure 5.**
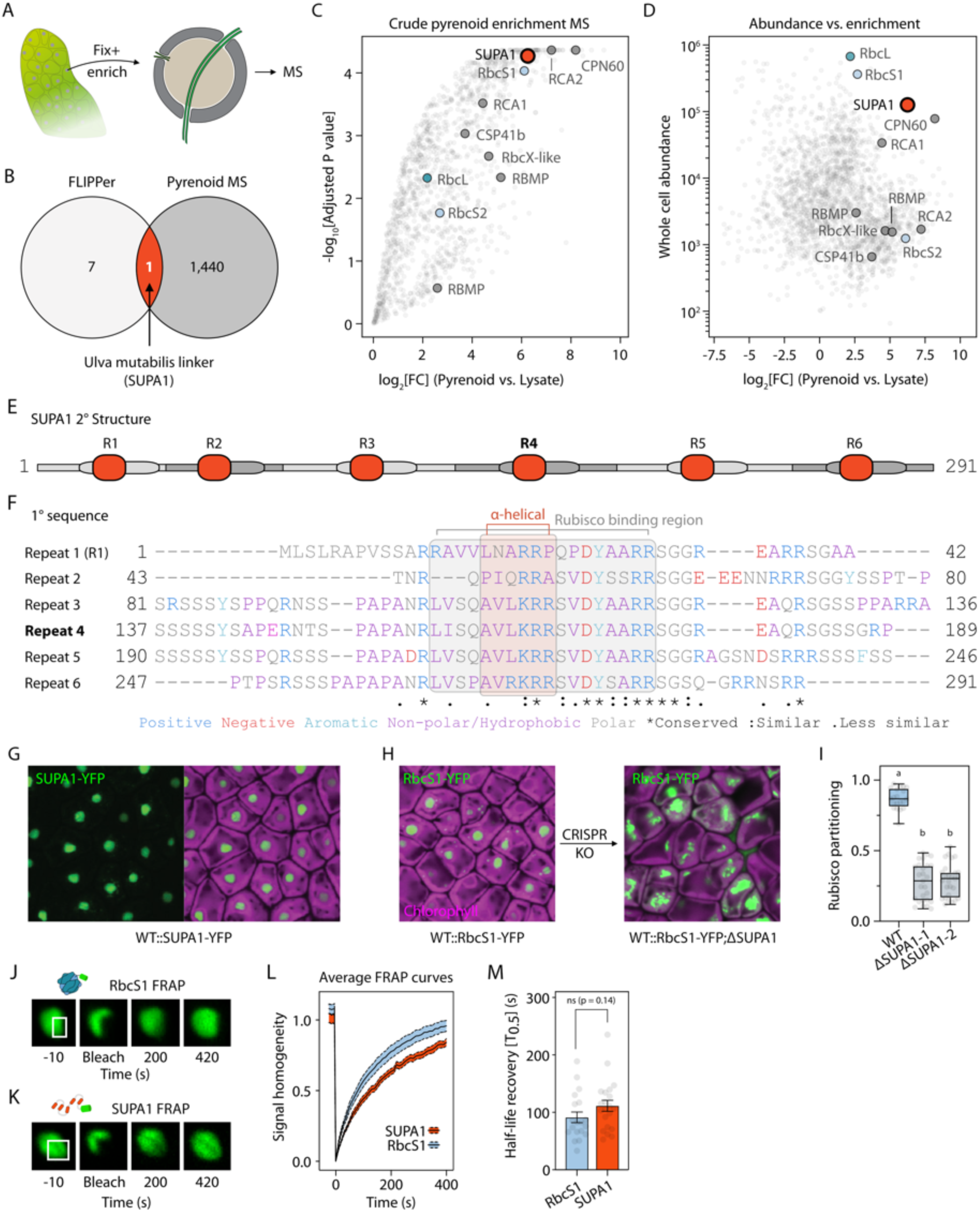
Identification and characterization of the SUPA1 *in vivo*. A. Schematic representation of pyrenoid enrichment from *Ulva* biomass. B. Venn diagram demonstrating identification of SUPA1 from FLIPPer and pyrenoid enrichment mass spectrometry experiments. C. Crude pyrenoid enrichment mass spectrometry experiment in which pyrenoid- localized and known Rubisco-interacting proteins are annotated. D. Pyrenoid enrichment vs whole cell abundance, with the same proteins annotated. E. Repeat and secondary structure of SUPA1 from subsequent experiments. F. Primary sequence of the repeated domains, colored by residue property. G. Localization of SUPA1-YFP in the pyrenoid of WT. H. CRISPR knockout of SUPA1 in a WT background expressing an additional copy of RbcS1 fused to YFP, demonstrating the relocalization of Rubisco in the absence of condensation by SUPA1. I. Quantification of Rubisco partitioning (from RbcS1-YFP signal) at the canonical pyrenoid position in WT and ΔSUPA1 CRISPR knockout lines. J. Snapshots of *in vivo* FRAP experiments completed on the pyrenoid of UmRbcS1-YFP lines. White box indicates the bleach region. K. *In vivo* FRAP experiments of SUPA1-YFP. L. Average FRAP recovery curves from *in vivo* experiments, where the dashed line indicates the s.e.m. and the solid line the mean signal homogeneity. M. Individual fitted *T*_0.5_ values from replicates (n = 19, 20).

#### SUPA1 condenses Rubisco into dynamic condensates

Fluorescent tagging confirmed that SUPA1 localized exclusively to the pyrenoid matrix, with a localization comparable to RbcS1 (**Fig. 5G**). To validate the role of SUPA1 in condensing Rubisco we generated CRISPR knockouts^26^ of SUPA1 in a RbcS1-YFP tagged strain. We targeted two independent sites in separate experiments, generating 4 and 7 individuals respectively (**Fig. 5H; Fig. S8**). Individuals from both lines showed a significant mislocalization of the RbcS1-YFP signal from the canonical pyrenoid position to a fragmented pattern throughout the chloroplast (Rubisco partitioning in the pyrenoid: Site 1, median 28%, IQR 15-39%; Site 2, median 30%, IQR 17-35%) relative to unedited control line (median 86%, IQR 81-94%). This is similar to the pattern observed for the *epyc1* knockout line in *Chlamydomonas*^12,102^, indicating that SUPA1 performs an analogous role and is essential to the condensation of *Ulva* Rubisco into the pyrenoid matrix (**Fig. 5G-I**).

In *Chlamydomonas*, it has been demonstrated that the pyrenoid possesses liquid-like properties, including internal mixing of matrix components over second-long timescales.^103^ Using our RbcS1-YFP and SUPA1-YFP tagged lines, we performed fluorescence recovery after photobleaching (FRAP) experiments *in vivo*, in which we monitored signal re-homogenization following bleaching. Using this approach, we observed full and comparable homogenization of the signal of both RbcS1 and SUPA1 (*T*_0.5_ = 91.32 ± 9.3 s s.e.m. and 111.4 ± 9.3 s respectively; **Fig. 5J-M**), albeit over longer timescales than observed for RbcS and EPYC1 in the *Chlamydomonas* pyrenoid (∼40 s and ∼20 s respectively).

To further validate the role of SUPA1 in Rubisco condensation, we purified Rubisco from *Ulva* thalli collected from the North Yorkshire coast and recombinantly produced and purified the full-length *Ulva* SUPA1 in *E. coli* (**Fig. S9; Fig. S10**). When mixed at concentrations close to those of Rubisco and linker proteins in characterized pyrenoid-containing species, we observed spontaneous demixing into micron-scale condensates, that was dependent on both components and sensitive to salt concentration (**Fig. 6A-D; Fig. S10**). In line with previous observations, titration droplet sedimentation assays of SUPA1 and Rubisco indicated a requirement for a ∼2-fold excess of SUPA1 to fully demix Rubisco.

**Figure 6.**
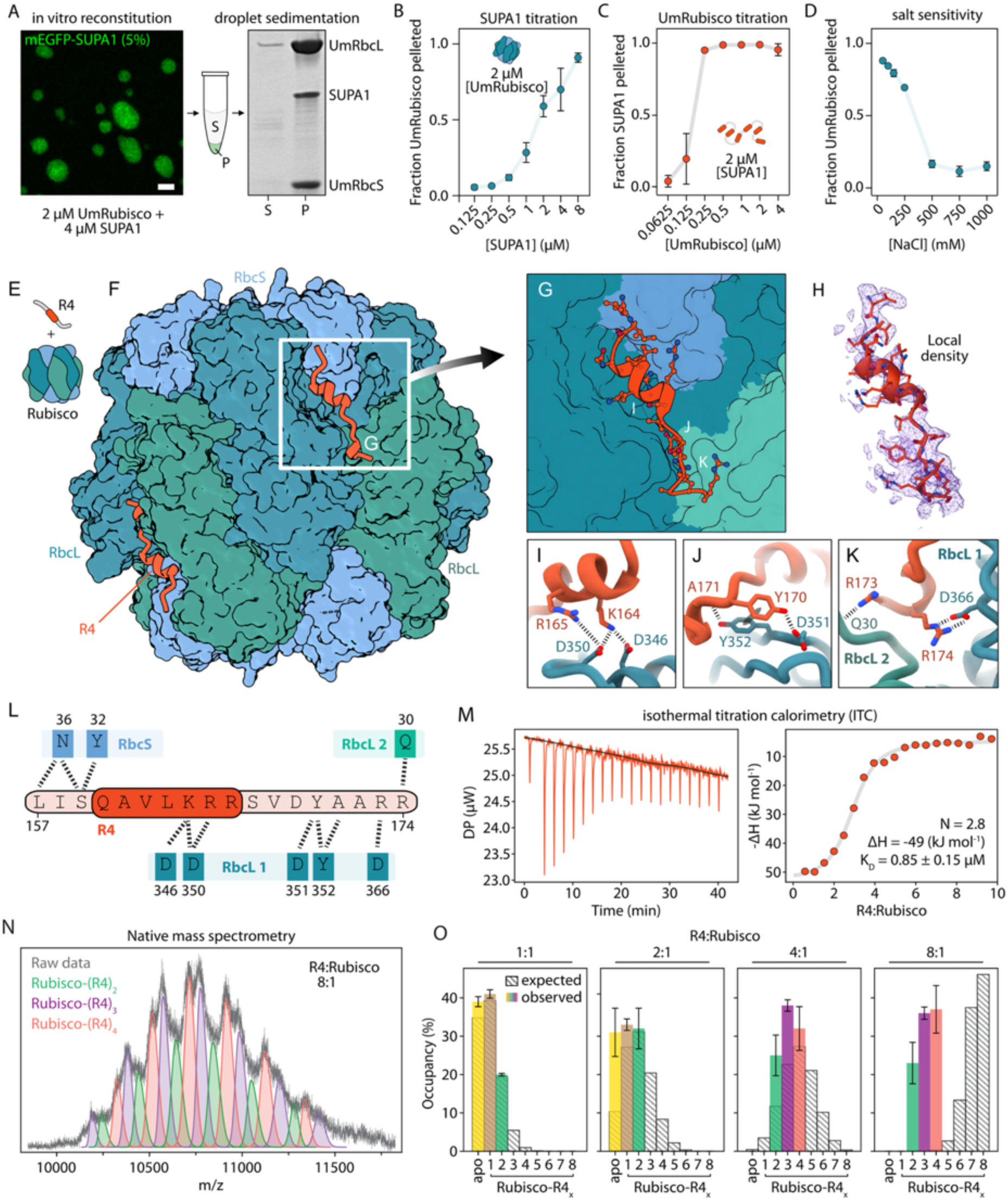
Biochemical, structural and biophysical characterization of Rubisco–SUPA1 interactions. A. Left: confocal fluorescence microscopy image of droplets in the *in vitro* reconstitution using SUPA1 and *Ulva* Rubisco. Scale bar = 5 µm. Right: SDS-PAGE analysis of the pellet (P) and supernatant (S) fractions following sedimentation of droplets. This experiment format was used to collect data in B, C and D. B. Titration droplet sedimentation assays with fixed UmRubisco concentration. See also **Fig. S10.** C. Titration droplet sedimentation assays with fixed SUPA1 concentration. See also **Fig. S10**. D. Titration droplet sedimentation assays with SUPA1 and UmRubisco at fixed concentrations (4 and 1 µM respectively) at varying salt concentrations. See also **Fig. S10**. E. Schematic representation of the SUPA1 R4 fragment and UmRubisco used for cryo-EM experiments. F. Overall structure of Rubisco with bound SUPA1 R4 peptide (orange), shown in surface/cartoon representation, revealing binding along an extended surface groove, RbcS is shown in lighter blue, RbcLs in dark blue/green. G. Close-up view of the SUPA1 R4 binding site. The peptide adopts an elongated conformation with a short α-helical segment embedded within the surface groove. The peptide interacts with three different subunits of Rubisco; two RbcLs and one RbcS. H. Local Cryo-EM map of the R4 fragment. I. Electrostatic interactions between K164/R165 of SUPA1 R4 and D346/D350 of RbcL 1. J. Interactions of SUPA1-R4 Y170 with RbcL 1. K. Interactions of the second group of basic residues of SUPA1-R4 (R173/R174) with RbcL 1 and RbcL 2. L. Map of the interactions between a single SUPA1-R4 fragment and the two RbcLs and one RbcS of UmRubisco. The helical region is highlighted, with interactions between residues shown by dashed lines. M. Isothermal titration calorimetry (ITC) analysis of SUPA1-R4 binding to Rubisco. Left: raw heat exchange data. Right: integrated heats fitted to an independent sites binding model, yielding an apparent dissociation constant *K*_D_ = 0.85±0.15μM with a stoichiometry N = 2.8. N. Native mass spectrometry of Rubisco incubated with SUPA1-R4 at an 8:1 R4:Rubisco molar ratio. Distinct species corresponding to Rubisco bound to 2, 3, or 4 SUPA1-R4 molecules are observed; no higher-order complexes are detected. See also **Fig. S14**. O. Quantification of Rubisco–R4 stoichiometries from four independent native mass spectrometry experiments performed at R4:Rubisco ratios ranging from 1:1 to 8:1. Colored bars indicate experimentally observed species, while dashed background bars represent the expected distribution, assuming eight independent Rubisco binding sites with the measured affinity. The discrepancy indicates limited site accessibility, consistent with overlapping binding sites.

#### Structural basis of SUPA1 binding to Rubisco

All Rubisco linker proteins characterized to date have revealed distinct interaction modes with Rubisco to fulfil their function.^19,104,105^ Cryogenic electron microscopy (cryo-EM) resolved the structure of *Ulva* Rubisco in complex with a 52-residue fragment corresponding to the fourth repeat (R4) of SUPA1, centered on the conserved sequence predicted to constitute the RBM (**Fig. 5E-F**). The reconstruction reached a resolution of 1.8–2.2 Å, with the R4- binding region resolved to ∼2.0 Å (**Fig. S11**), enabling unambiguous modeling of the central 18 residues and their interactions with the Rubisco surface (**Fig. 6E-L**).

R4 binds within an extended surface groove formed at the interface of two RbcLs and one RbcS such that each peptide simultaneously engages three of the 16 subunits of the Rubisco holoenzyme (**Fig. 6E-G**). Two RbcLs form the structural core of the complex, while one RbcS contributes to shaping the binding pocket. Within the binding groove, R4 adopts an elongated conformation interrupted by a short α-helix (**Fig. 6F-H**). There was no evidence of a helical element in 2D NMR spectra of the isolated R4 fragment (**Fig. S12**), supporting a previously undescribed folding-upon-binding mechanism for Rubisco condensation.

The interface combines hydrophobic and electrostatic contacts similar to linker interactions described in *Chlorella* and *Chlamydomonas*. Binding is mediated by two clusters of basic residues of the peptide (K164/R165 and R173/R174; **Fig. 6G-L**), which engage complementary acidic patches on the Rubisco surface and anchor the peptide within the groove. A conserved aromatic residue (Y170) further stabilizes the complex through hydrophobic and π-stacking interactions.

Despite the presence of multiple binding grooves on the Rubisco surface created by its D4 symmetry, inspection of the cryo-EM density revealed a fundamental geometric constraint: density was continuous across the symmetry interface at the intersection of two adjacent SUPA1 sites (**Fig. S11E-F**). Attempts to model two linkers into adjacent sites created unavoidable steric clashes, demonstrating that these binding sites spatially overlap. We therefore interpreted this continuous density as an artefact arising from symmetry averaging of two partially occupied sites that could not be disentangled with classification, most likely because of the small size of the linker. This geometric constraint implies that not all symmetry-related binding grooves can be occupied at the same time, effectively reducing the stoichiometry of SUPA1 binding per Rubisco holoenzyme. Consistent with this interpretation, the R4 region exhibited elevated B-factors (median 131.3 A^2^ for SUPA1 chain vs 57.7 A^2^ overall) and reduced occupancy (average 0.82), indicative of partial and/or dynamic binding at these sites (**Fig. S11G**).

We next quantified the Rubisco–R4 interaction using isothermal titration calorimetry (ITC). ITC measurements revealed a clear binding signal (**Fig. 6M**) and fitting the integrated heats to an independent-sites binding model yielded an apparent dissociation constant of *K*_D_ = 0.85 ± 0.15 μM with a stoichiometry of *N* = 2.8 R4 peptides per Rubisco complex. This substoichiometric binding is consistent with the structural observation of overlapping binding sites. Further, the *K*_D_ is greater than two orders of magnitude higher than cognate single RBM binding to *Chlamydomonas* and *Chlorella* Rubisco^19,105,106^, where there are eight simultaneously-available binding sites on Rubisco. To validate the structural determinants of binding, we examined R4 mutants targeting key interacting residues. Mutation of the N-terminal basic pair (K164A/R165A) completely abolishes detectable binding in ITC experiments (**Fig. S13A**). Similar loss of interaction was observed for Y170D and R173A/R174A mutants (**Fig. S13B,C**), confirming that both electrostatic anchoring and aromatic interactions are essential for complex formation, which was also confirmed by the abolition of stable complex formation in native PAGE experiments (**Fig. S13D**).

Finally, we assessed the stoichiometry of R4 binding using native mass spectrometry. At a R4:Rubisco molar ratio of 8:1, distinct species corresponding to Rubisco bound to two, three, or four R4 molecules were detected, but no higher-order complexes were observed (**Fig. 6N; Fig. S14**). Quantification across four independent experiments and a range of R4:Rubisco ratios (1:1 to 8:1) confirms this limited occupancy (**Fig. 6O; Fig. S14**). While accurate quantification of partially overlapping mass spectral peaks requires careful interpretation, the experimentally observed distributions deviate strongly from those expected for eight independent binding sites with the measured affinity. Even allowing for substantial uncertainties in peak integration, the absence of higher-order species and the pronounced under-occupancy are inconsistent with a model of fully independent sites, instead supporting a scenario in which only a subset of sites is accessible at any given time due to spatial overlap.

Together, these biochemical, structural and biophysical data establish that SUPA1 assembles Rubisco into a dynamic condensate through multivalent sequence-specific interactions that force inter-Rubisco cross-linking via imposing geometric constraints that limit the number of available Rubisco binding sites. The reduced Rubisco binding site availability of the *Ulva* SUPA1-Rubisco system is potentially compensated for by the increased RBM binding affinity compared to the *Chlorella* and *Chlamydomonas* systems.^19,105,106^ Further, it highlights how oligomeric architecture can regulate ligand accessibility independently of local binding affinity.

### Pyrenoid structural components contain a Rubisco-binding motif

In *Chlamydomonas*, assembly of the pyrenoid with its ultrastructural features (traversing thylakoids and surrounding starch sheath) is controlled by a number of structural protein components which contain RBMs similar to those found in the linker protein EPYC1.^22^ To see whether a similar RBM-based pyrenoid assembly/organization framework exists in *Ulva* we used an iterative MEME suite approach based on the Rubisco-binding sequence of the SUPA1 protein to search the *Ulva* translated genome. Using this approach, we identified 4 additional proteins that shared high confidence variants of the RBM: UM003_0241, UM004_0280, UM016_0026, UM047_0002 (**Fig. S15A-B**), 3 of which contained 7 or more RBMs and another with only a single RBM. In line with our structural characterization, the key Rubisco-interacting residues were highly conserved between motifs, with more variation observed outside these regions. Although not identified by the MEME analysis, subsequent manual analysis revealed two partial RBM matches in the PTT-localized BST UM047_0048 in its disordered N-terminal domain (**Fig. 3; Fig. S15C**). The number of *Ulva* proteins containing the SUPA1 RBM is approximately an order of magnitude lower than the number of *Chlamydomonas* proteins that contain the EPYC1 RBM^22^, although speculative this may be due to the lower structural complexity of the *Ulva* pyrenoid requiring less proteins to be targeted to the pyrenoid.

We obtained imaging data from all 4 additional proteins that contain RBMs, which all localized to the pyrenoid. UM004_0280, a largely disordered protein with no other annotated domains, was distributed similar to SUPA1 throughout the pyrenoid matrix (**Fig. 7A**). This protein probably plays an accessory role in Rubisco condensation since it is lowly expressed and not detected in whole cell MS (**Table S2**), so we named it Pyrenoid *Ulva* Matrix Associated1 (PUMA1). Two of the RBM-containing proteins contained additional domains that suggested a role in association with other structural features of the pyrenoid. We named UM016_0026, which contains 7 RBMs and a PGRL1 domain, Rubisco-binding PGRL1 Domain-containing protein1 (RPGD1; **Fig. 7A; Fig. S16**). The C-terminal PGRL1 domain suggests that RPGD1 is a cyclic electron flow (CEF)/ PSI-associated protein.^107,108^ RPGD1 displayed non-homogeneous localization in the pyrenoid, consistent with its presence in the traversing thylakoid, but in some cases was also seen at the pyrenoid periphery. It is tempting to speculate that the enrichment of RPGD1 in and at the periphery of the pyrenoid is to regulate CEF that is the principle energetic driver of pCCMs.^109^ UM047_0002 contains 4 RBMs with an N-terminal syntaxin-like domain, and a coiled-coil with structural similarity to membrane-associated proteins (ESCRT-III, VIPP, t-SNARE; **Fig. 7A; Fig. S16**), suggesting a role in PTT remodeling.^110^ We name this protein Rubisco-binding Syntaxin-Like Domain-containing protein 1 (RSLD1), its signal was enriched at the periphery of the pyrenoid matrix, consistent with a localization within, or adjacent to the matrix-associated starch sheath. Surprisingly, we did not identify proteins containing both RBMs and characterized starch-binding domains analogous to SAGA1 and SAGA2 that are required for localizing the starch sheath to the pyrenoid in *Chlamydomonas.*^22,96,111^ Starch sheath assembly may proceed via an alternative mechanism in *Ulva,* or alternatively one of the RBM-containing proteins may interact with a protein(s) that meditates starch sheath formation and matrix attachment. In similar fashion, RBM-containing SAGA1 interacts with MITH1 to mediate PTT formation in *Chlamydomonas*.^97^ UM003_0241 is a RbcX-like Rubisco assembly chaperone that contains a single C-terminal RBM and localizes to the pyrenoid matrix and in peripheral puncta similar to RbcS1. In *Chlamydomonas*, Rubisco activase (RCA) a AAA+ chaperone that removes inhibitory sugar phosphates, is able to partition into EPYC1-Rubisco droplets with this interaction possibly mediated by an RBM.^112^

**Figure 7.**
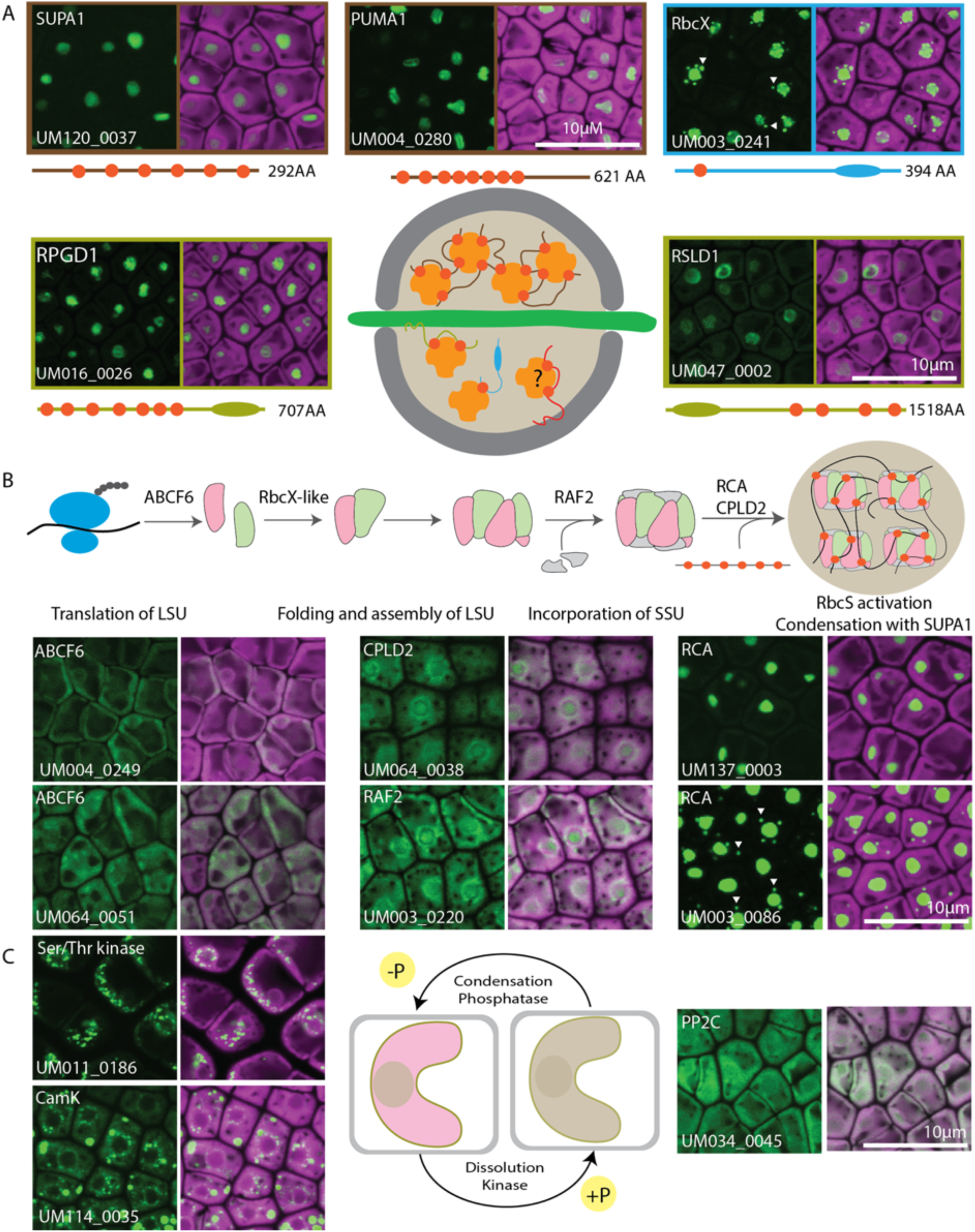
Proteins binding Rubisco and affecting condensation, assembly and regulation. A. RBM-containing proteins are exclusive to pyrenoid subdomains. Red circles indicate RBMs, blue and green ovals additional protein domains. All RBM-containing proteins in this figure are boxed in a color that represents their subclass: Orange: Rubisco SSU; Brown: Rubisco linkers; Green: thylakoid tethering proteins RPGD1 and RSLD1; Blue: RBM-protein with additional RbcX protein domain. We did not identify protein containing RBM and association to the starch sheath (red). Magenta: Chlorophyll autofluorescence; Green: YFP signal of tagged protein. White arrowheads indicate puncta at pyrenoid periphery. Scale: 10 µm. B. Proteins involved in Rubisco translation, assembly and activation are associated with the pyrenoid. Magenta: Chlorophyll autofluorescence; Green: YFP signal of tagged protein. White arrowheads indicate puncta at pyrenoid periphery. Scale: 10 µm. C. Two Kinases and a phosphatase are putatively involved in regulation of condensation and dissolution of the pyrenoid. Magenta: Chlorophyll autofluorescence; Green: YFP signal of tagged protein. Scale: 10 µm.

#### Conservation of SUPA1

Homologs of both previously characterized Rubisco linker proteins in the green lineage are apparently only present within their respective taxonomic orders (Chlamydomonadales and Chlorellales for EPYC1 and CsLinker, respectively). In line with this, within the Ulvophyceae, we observed homologs of SUPA1 only in species of the Ulvales order, including those of the Ulvaceae, Ulvellaceae, Kornmanniaceae and Bolbocoleonaceae families (**Table S6**). We also identified homologous SUPA1 fragments in reads of the transcriptomes of three species of rhodophytes, but not in any assembled genomes. It remains to be seen whether the presence of these reads is an artefact or not.

### Rubisco likely assembles before integrating into the pyrenoid matrix

Since the assembly of the pyrenoid largely depends on interactions of proteins with Rubisco^22^, the correct assembly and activation of the holoenzyme is crucial. Rubisco holoenzyme assembly is a tightly controlled complex multi-chaperone assembly process.^114^ We have localized two Rubisco assembly chaperones, Rubisco Assembly Factor 2 (RAF2; UM003_0220) and RBM-containing RbcX-like (UM003_0241; **Fig. 7A-B**). RAF2 likely mediates proper folding of nuclear-encoded RbcS and makes these subunits available to the plastid-encoded LSU for holoenzyme assembly (**Fig. 7B**).^115,116^ In line with this function, *Ulva* RAF2 is localized broadly in the chloroplast, but enriched at the pyrenoid periphery and in the matrix. RbcX-like mediates the generation of LSU Rubisco oligomers and is enriched in the pyrenoid matrix but also in puncta at the periphery. Interestingly, RbcS1 shows a similar localization at the pyrenoid periphery (**Fig. 4A; Fig. S6**), suggesting that holoenzyme assembly may take place at the pyrenoid periphery before Rubisco is integrated into the pyrenoid.

In addition to assembly chaperones, Rubisco requires chaperones for maintenance with RCA playing a central role.^117^ Interestingly, one RCA localizes to the matrix and in high-expressing independent lines in peripheral puncta (UM003_0086), a second RCA (UM137_0003) is matrix-exclusive (**Fig. 7B**). One of the inhibitory sugars, xylulose-1,5-biphosphate (XuBP), is removed by phosphatase CbbY (CPLD2, UM064_0038) that is expressed in the chloroplast and matrix enriched (**Fig. 7B**). Protein translation in the *Chlamydomonas* chloroplast is known to occur in a specific translation zone near the pyrenoid.^118^ Also in *Ulva* two orthologs of a ribosome-binding enzyme ABCF6 (UM004_0249 and UM064_0051) are partially enriched adjacent to the pyrenoid (**Fig. 7B**). Combined, our observations suggest that Rubisco holoenzyme assembly and activation occur in the pyrenoid periphery, possibly in close proximity to the location of LSU translation. Notably, SUPA1 is not found in the peripheral puncta, enforcing that only complete Rubisco holoenzymes may be condensed into the pyrenoid.

### Putative regulators of pyrenoid dynamics

Although the morphology of a pyrenoid looks simple under a light microscope, the complex dynamics of this structure during cellular processes like cell division was already appreciated in the first microscopic observations of *Ulva*.^93,94^ The rapid (dis)assembly of condensates is often mediated by (de)phosphorylation of key proteins. Rubisco linkers can be phosphorylated under low CO_2_^119^ and a kinase is known to disrupt the interaction between Rubisco and its main linker in *Chlamydomonas*.^120^ Interestingly, two chloroplast-localized proteins, UM011_0186 and UM147_0014, are putative serine/threonine kinases (**Table S1; Fig. 7C**). UM011_0186 localizes in puncta to the pyrenoid periphery. We also identified a putative calmodulin-dependent protein kinase (UM114_0035) that localizes to the pyrenoid periphery in puncta (**Fig. 7C**). Several other protein kinases are present in the *Ulva* pCCM-associated proteins but were not cloned. However, only one of these (UM018_0146) is predicted to localize to the chloroplast (**Table S2**).

Since phosphorylation is a key regulator of pyrenoid dissolution, it is likely that removing phosphorylation leads to recondensation. In our screen we identified a chloroplast protein phosphatase 2c (PP2C) family member, UM034_0045, that localizes throughput the chloroplast but is found in the pyrenoid (**Fig. 7C**). Interestingly, UM034_0045 contains a disordered region similar to CO_2_-sensing PP2Cs that form condensates upon changes in CO_2_ levels.^121^

Taken together, we have identified candidates that might regulate rapid dynamics of the *Ulva* pyrenoid. Altering the function of the *Chlamydomonas* kinase results in a chloroplast containing multiple pyrenoids.^120^ This phenotype occurs naturally in *Ulva* and was one of the morphological descriptors to determine species identity in the pre-molecular era.^122,123^ Further studies are needed to evaluate if the presence or activity of proteins that mediate post-translational modifications can explain this morphological variation.

### Proteins associate to *Ulva* pyrenoid subdomains

In addition to our finding that orthologs of core structural proteins are absent in *Ulva*, we wanted to understand how conserved the localization of 41 known pyrenoid-exclusive *Chlamydomonas* proteins^13^ are in *Ulva*. 29 of the genes have at least one ortholog in *Ulva*, and we localized at least one of these orthologs for 22 targets (see below, **Fig. 4**; **Table S5**). In addition to structural pyrenoid proteins like EPYC1 and SAGA1/2, further missing orthologs in *Ulva* include proteins that are less well-characterized, such as Cre16.g648400, Cre02.g143635, Cre03.g172700, PsbP-like protein (PSPB4), Structural Maintenance of Chromosomes 7 (SMC7), Cre09.g394547, Cre09.g415600 and Cre09.g394510 (**Table S5**).

Surprisingly, only eight proteins were exclusively localized to the same pyrenoid subdomain in both *Ulva* and *Chlamydomonas*: RbcS1-2, Rubisco Activase, CAH3 and the orthologs of EPOS1, SULFURTRANSFERASE 16 (STR16; UM149_0040) and Chloroplast Stem-loop Binding Protein of 41 kDa A-B (CSP41a-b; **Fig. 4A-B**; **Fig. 8; Table S5**). STR16 is a matrix-exclusive rhodanese (thiosulfate sulfurtransferase) domain containing protein^53^ whose exact role is still unclear. Interestingly, the rhodanese domain of *Chlamydomonas* RBMP2 is necessary for the PTT extension in the matrix.^98^ CSP41a (UM005_0157) and CSP41b (UM017_0083) both localized to the pyrenoid matrix (**Fig. 4A**; **Fig. 8**). CSP41 proteins are best characterized for binding and stabilizing chloroplast mRNAs coding for photosynthetic proteins and rRNAs in a redox-dependent manner.^22,124^ However, CSP41 proteins have endonuclease activity^125^ and could play a role in degrading DNA or oxidized RNA that has been shown to accumulate in the pyrenoid in *Chlamydomonas* but also the green seaweed *Caulerpa.*^126,127^

**Figure 8.**
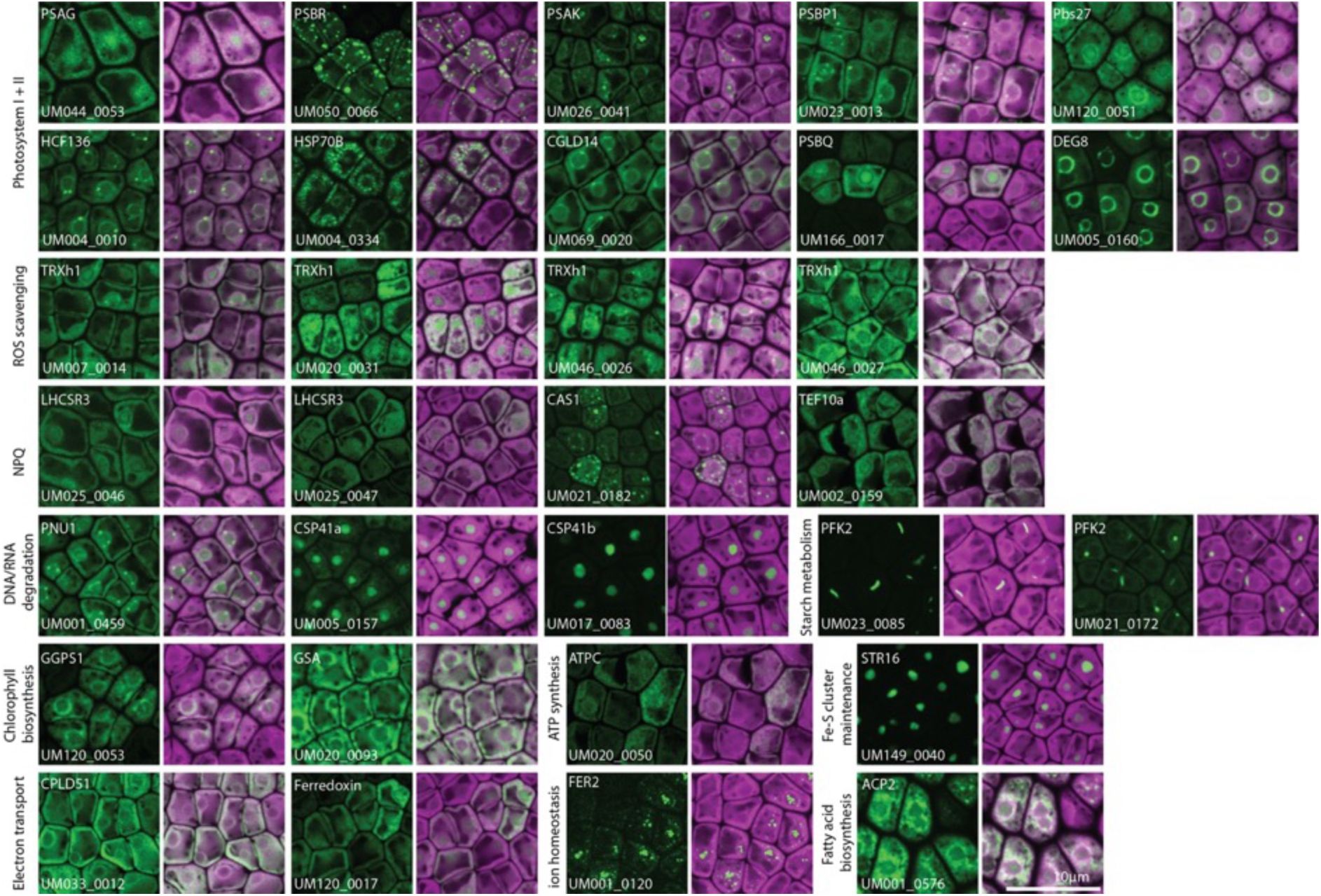
Other pathways are associated with the pyrenoid. Proteins involved in photosynthesis, electron transport, NPQ, ROS scavenging, ion homeostasis, DNA/RNA degradation, ATP synthesis; starch metabolism, FE-S cluster maintenance and fatty acid biosynthesis are present or enriched at pyrenoid subdomains. Magenta: Chlorophyll autofluorescence; Green: YFP signal of tagged protein. Scale: 10 µm.

As discussed, there are substantial architectural differences between the multiple pyrenoid tubules found in *Chlamydomonas* and the singular PTT of *Ulva*. Interestingly, several proteins exclusively localized to the pyrenoid tubules in *Chlamydomonas*, are present at the pyrenoid periphery and enriched at the presumed thylakoid convergence zones in *Ulva*: Pyrenoid nuclease 1 (PNU1, UM001_0459), Conserved in the green lineage and diatoms 14 (CGLD14, UM069_0020), DegP-type protease (DEG8, UM005_0160), High chlorophyll fluorescence 136 (HCF136, UM004_0010) and PS II subunit P1 (PSBP1; UM023_0013; **Fig. 4C**; **Fig. 8**). The precise mode of action of these proteins is largely unknown; PNU1 is likely involved in RNA metabolism while CGLD14, DEG8, HCF136 and PSBP1 are proteins associated with PSII activity.^21^ The observation that several *Chlamydomonas* pyrenoid tubule proteins are localized to the pyrenoid periphery in *Ulva* suggests that potential secondary pyrenoid functions outside of CO_2_-fixation associated with the *Chlamydomonas* pyrenoid^53^ may utilize the thylakoid convergence zones. This could be dependent on the large membrane surface area of the thylakoid convergence zones providing a functional equivalence of the pyrenoid tubules and reticulated region of the *Chlamydomonas* pyrenoid. Independently, the frequent observation of protein enrichment at the thylakoid convergence zones further suggests that this is an important regulatory hub (**Fig. 4C**; **Fig. 8; Table S2**).

#### The pyrenoid periphery connects carbon capture to metabolism, photosynthesis and regulation

It is increasingly appreciated that the vicinity of the pyrenoid is an important site for a variety of processes downstream of carbon capture like the Calvin–Benson–Bassham (CBB) cycle and L-serine biosynthesis.^21^ Here, we provide evidence for carbon catabolic pathways associated with the pyrenoid. Two phosphofructokinases (PFK2, UM001_0120 and UM021_0172) localize adjacent to the pyrenoid periphery in elongated puncta (**Fig. 8**). Chloroplast PFKs regulate starch metabolism, suggesting that these proteins could be involved in regulation of the starch sheath at the pyrenoid periphery. Interestingly, *Chlamydomonas* PFK2 localizes to similar elongated structures and interacts with pyrenoid periphery proteins EPOS1 and SBE3.^16,99^

Cellular processes are dependent on energy generated during photosynthesis. Photosystems are synthesized in and localized to the translation zone in*Chlamydomonas*.^16,128,129^ In line with these observations, several *Ulva* proteins associated with photosynthesis were enriched at the periphery and often matrix of the pyrenoid (**Fig. 8**). Besides known pyrenoid-associated proteins of Photosystem I (PSI; PSAG, UM044_0053 and PSAK, UM026_0041) and Photosystem II (PSII; Pbs27 (UM120_0051), PSBQ (UM166_0017), PSBR (UM050_0066) and PSBP1), we also localized putative PSII-associated proteins HSP70B (UM004_0334), DEG8, HCF136 and CGLD14 to the matrix and/or periphery (**Fig. 4**; **Fig. 8**). Similarly, ATP synthase component ATPC (UM020_0050), proteins involved in electron transport CPLD51 (UM033_0012) and Ferredoxin (UM120_0017) and proteins involved in non-photochemical quenching such as light-harvesting complex stress-related 3 proteins (LHCSR3, UM025_0046 and UM025_0047), Calcium Sensor 1 (CAS1; UM021_0182) and TEF10a (UM002_0159) are present or enriched at the pyrenoid matrix or periphery (**Fig. 8**). Finally, two proteins involved in chlorophyll biosynthesis (Glutamate-1-semialdehyde aminotransferase (GSA: UM020_0093) and Geranylgeranyl pyrophosphate synthase (GGPS1: UM120_0053) and an acyl-carrier protein involved in fatty acid biosynthesis (UM001_0576) are enriched at the periphery (**Fig. 8**).

Since the pyrenoid is an important hub for photosynthesis, maintaining a redox balance is critical. Thioredoxins are considered redox transmitters because a conserved dithiol group of thioredoxin proteins is reduced and this reductive power is subsequently transferred to its specific target proteins, for example the light-dependent regulation of CBB cycle enzymes.^130^ We cloned six thioredoxin h-type genes and four (TRXh1, UM007_0014, UM020_0031, UM046_0026 and UM046_0027) localized to the periphery and matrix (**Fig. 8**).

Taken together, our data suggests that photosynthetic processes, catabolic and anabolic pathways and redox scavenging are enriched at the pyrenoid periphery.

#### The pyrenoid matrix favors oxygen-sensitive reactions

Several chloroplast proteins rely on metal cofactors and anions. Ion homeostasis is crucial for many photosynthetic processes including photosystem and Rubisco function.^131^ Fe is the prominent transition metal for protein complexes in the photosynthetic electron chain and Fe^2+^ ligates proteins in PSI and II, cytochrome *b*6/*f*, Fe–Sulfur [Fe–S] clusters and ferredoxin.^132^ These proteins mediate crucial metabolic reactions in the oxygen-rich environment of the chloroplast, but paradoxically Fe-S cluster-dependent metabolism is sensitive to oxygen.^133^ In addition to the matrix localization of Fe-S cluster protein Ferredoxin and the rhodanese-domain protein STR16 mentioned above, we observed a clear enrichment of a ferritin subunit (FER2, UM023_0085) in the matrix (**Fig. 4**; **Fig. 8**). Similarly, the pyrenoid is enriched in Fe-S assembly and Fe-S-containing proteins in *Chlamydomonas*, and the high CO_2_:O_2_ ratio in the pyrenoid is proposed to favor oxygen-sensitive biological reactions.^53^ Ferritin sequesters iron that likely acts as storage for Fe-S assembly and other photosynthesis-related proteins in the vicinity of the pyrenoid. Taken together, these observations suggest a spatial regulation of chloroplast ion homeostasis and oxygen-sensitive reactions in the pyrenoid.

## Perspective

We have integrated large-scale protein tagging with biochemistry and structural biology to provide a comprehensive overview of pCCM carbon fixation in a seaweed. The most comprehensive mapping to date has been done in the single cell terrestrial/freshwater alga *Chlamydomonas*. Our work is of comparable scale but applies it to a marine intertidal multicellular seaweed, highlighting core similarities and major differences between pCCM spatial organization, pyrenoid assembly, function and evolution within and outside of the green lineage (**Fig. 9A-B**). Efficient stable transformation remains rare in algae, and particularly rare in seaweeds, due to transgene silencing.^60,134^ Our work shows that *Ulva* is amenable to screening large collections of constructs and thereby overcomes a crucial throughput barrier previously only possible in *Chlamydomonas* for the eukaryotic algae. The observed high transgene expression and success rate of functional constructs further consolidates *Ulva* as a transformative, functional biology model.^39^

**Figure 9.**
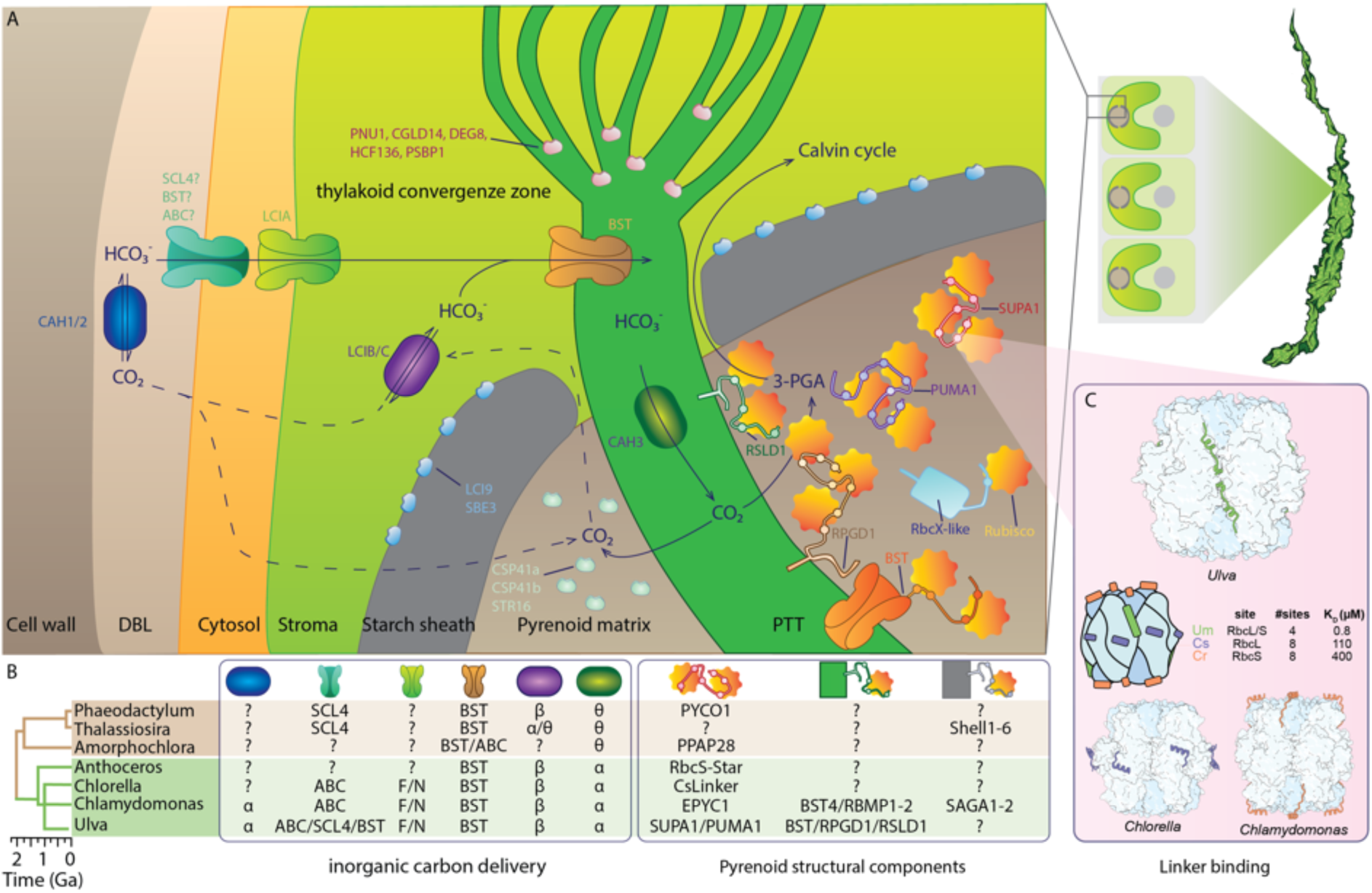
Molecular model and evolution of the ulvophyte CCM. A. Summary of the *Ulva* pCCM pathway with the extracellular (blue), chloroplast (purple) and PTT (green) carbonic anhydrases; carbon transporters at the plasma membrane (blue), chloroplast envelope (green) and thylakoid (orange); and pyrenoid structural proteins SUPA1 (red), PUMA1 (purple), BST (orange), RbcX-like (blue), RSLD1 (green) and RPGD1 (light orange) binding to Rubisco (yellow/orange). Also, protein associated with pyrenoid periphery (blue), matrix (green) and thylakoid convergence zone (pink) are indicated. B. The evolution of the pathway is illustrated for proteins involved in inorganic carbon delivery, pyrenoid structural proteins and linker binding to Rubisco. C. Unique binding locations and characteristics of green algae Rubisco linker proteins. The location, number of binding sites and dissociation constant K_D_ is shown for the linker of *Chlamydomonas*, *Chlorella* and *Ulva*.

Our elucidation of the CO_2_-fixing pathway in *Ulva* supports the hypothesis that the inorganic carbon transport components of the CCM (e.g. Ci transporters, channels and CAs) are likely of a common, ancient origin in the green lineage (**Fig. 9 A-B**). In stark contrast, our identification and characterization of SUPA1 and RBM containing proteins suggests that pyrenoid assembly differs significantly, and supports that Rubisco condensation and subsequently pyrenoid functionalization likely emerged more recently (**Fig. 9B-C**).^11,135^ Comparison of SUPA1, with EPYC1 and CsLinker indicates that the high-level architecture of Rubisco linker proteins within the core-chlorophytes, composing of five or six RBMs separated by predicted flexible disordered spacer regions, is conserved but that the properties of the linkers and emergent properties of their pyrenoids differ significantly. Through our multi-discipline approach, we have revealed that biomolecular condensation of *Ulva* Rubisco is mediated via the unique mechanism of SUPA1 binding to the large and small subunits. Whilst there are eight potential binding sites only four can be simultaneously occupied due to binding resulting in steric clashes preventing adjacent site binding. This contrasts to EPYC1 and CsLinker that respectively bind their cognate RbcS and RbcL at eight binding sites (**Fig. 9C**).^19,105^ Another unique feature of the *Ulva* system is a structural change of the RBM upon Rubisco binding, with the central RBM region unfolded in solution but forming a helical conformation upon binding. Potentially compensating for the reduced number of binding sites is the higher RBM affinity of SUPA1 compared to EPYC1 and CsLinker, that in turn may impact material properties, such as viscosity as suggested through slower FRAP recovery rates. Collectively, this supports that Rubisco condensation, the core defining feature of a pyrenoid^135^, has convergently evolved in the core-chlorophytes through multivalent linker proteins but the emergent biophysical properties and co-adopted accessory proteins show significant differences. These differences in material properties and accessory proteins may explain the large diversity in pyrenoid ultrastructure between lineages.

Although we uncovered a comprehensive molecular map of pyrenoid-based seaweed carbon fixation, our dataset relies heavily on ortholog identification based on sequence similarity. For example, the pyrenoid also contains a large number of proteins without RBMs in *Chlamydomonas.*^53^ With our library of tagged lines, additional *Ulva*-specific pCCM components can be revealed, *e.g.* by determining their protein interactions using more unbiased methods like affinity purification and proximity labeling. These methods might shed light on the mechanisms of Rubisco tethering to the starch sheath and regulators of pyrenoid dissolution-condensation.

The green seaweeds (Ulvophyceae) display an interesting diversity in cytology and habitat preference (marine, brackish and even terrestrial), including lineages without pyrenoids.^54,136,137^ Since pCCMs are the product of convergent evolution that occurred after the diversification of most lineages, this variability likely translates to pCCM variations. For example, terrestrial lineages likely rely more on CO_2_ diffusion and less on active HCO_3_^-^ transport compared to aquatic organisms, similar to pyrenoid-containing hornworts.^20^ The identification of additional linkers or alternative mechanisms to condensate Rubisco promises to further reveal principles underlying the convergent evolution of pyrenoids. The green seaweeds could therefore become important model organisms to better understand the molecular variation and evolution of carbon fixation pathways.

A large focus of understanding pyrenoid-based CCMs has been with a view to translating these mechanisms into C_3_ crop plants to improve photosynthetic yields.^14,138,139^ Our work provides a powerful insight into how a pCCM is assembled and functions in a multicellular system. Combined with the recent advances in understanding hornwort pCCM structure and function^20,57^, these advances are providing a framework for the requirements to functionalize engineered pyrenoids in plants.^14,139,140^ Additionally, the structurally simpler organization of the *Ulva* pyrenoid compared to the model template of *Chlamydomonas*, with a single planar PTT, two starch plates and inferred reduced structural protein set, and validated multicellular functionality make an *Ulva*-based pCCM engineering strategy an exciting prospect.

In summary, seaweeds are keystone organisms that play fundamental roles in marine ecosystems and global biogeochemical cycles, but until now have been recalcitrant to robust large-scale genetic manipulation. Removing this limitation in *Ulva* now allows the detailed elucidation of biological pathways for the first time in multicellular algae. Beyond a fundamental understanding of the fixation of inorganic carbon into algal biomass, the work we present here is a starting point to understand why *Ulva* thrives in coastal ecosystems worldwide and how it can dominate during blooms.

## Resource availability

### Materials availability

The authors declare that the data supporting the findings of this study are available within the paper and its Supporting Information.

All vector maps are available at https://vectorvault.vib.be/ulva-mutabilis.

### Data and code availability

Proteomics data are deposited in MassIVE (ID: MSV000102502) with ProteomeXchange identifier PXD081181.

Electron density maps were deposited in EMDB with accession code EMD-58973. The corresponding coordinates are deposited in the PDB with accession code 32LL.

MBG-Fit is available on Github (https://github.com/cdegut/MsMBG-Fit). FLIPPer is available on Github (https://github.com/james-r-barrett/FLIPPer). Code for the integrated *Ulva* dashboard and dataset is available on Github (https://github.com/james-r-barrett/ulva_db).

Protein localizations are annotated on ORCAE (https://bioinformatics.psb.ugent.be/orcae/overview/Ulvmu).

## Acknowledgements

J.Blo., A.S.C., E.C.V. and O.D.C. are indebted to Research Foundation – Flanders (FWO) for funding (Senior Research Project G015623N, postdoctoral fellowship 12T3418N and ERC StG runner-project G0AHR24N).

J.Blo. and T.B.J. thanks Ghent University Special Research fund (postdoctoral fellowship BOF20/PDO/016 and starting grant BOF/STA/202209/016) and VIB Tech Watch Project DeRisking Funding.

J.Blo. and O.D.C. thank the Gordon and Betty Moore Foundation (Symbiosis in Aquatic Systems Initiative #9329).

TEM imaging was performed with the help of Anneke Kremer and Michiel De Bruyne at the VIB BioImaging core Ghent.

J.B., C.D., M.H., H.D., M.J.P., and L.C.M.M. were funded by EPSRC grant EP/W024063/1 as part of the York Physics of Pyrenoids Project (YP3). We would also like to thank the YP3 consortium and Mackinder Lab for fruitful discussions.

J.B. was supported by BBSRC studentship (BB/M0111511a).

J.B., H.D. and L.C.M.M. were supported by a CTRF grant (AP23-1_023), Bill and Melinda Gates Foundation/ Foreign Commonwealth and Development Office funding (INV-054558).

L.C.M.M. was funded by UKRI-Future Leaders Fellowship (MR/T020679/1 + MR/Y034074/1); BBSRC-NSF/BIO (BB/S015337/1 and BB/Y000323/1)\

J.N.B. was supported by a UKRI Future Leader Fellowship (MR/ T040742/1).

## Authors contributions

J.B.: Performed pyrenoid enrichment and whole cell MS experiments, identified and cloned SUPA1 and variants, performed in vitro experiments, analyzed and interpreted data, created figures and contributed to writing the manuscript.

C.D.: Purified Rubisco samples for cryoEM and native Mass Spec. CryoEM, ITC and native MS data collection. Processing and analysis of cryoEM, ITC and MS data.

A.S.C.: Performed cloning. Did SUPA1 targeted knock-out. Analyzed and interpreted the data. Created figures.

M.R.H: Purified SUPA1 samples for CryoEM and native MassSpec. H.D.: Provided bioinformatic support.

A.S.: Optimized SUPA1 and Rubisco purifications. E.C.V: Provided bioinformatic support. Created figures.

R.B.: MS data collection and supervised MS data analysis. T.B.J.: Designed and supervised the study. Provided resources. O.D.C.: Designed and supervised the study. Provided resources. J.H.Bot.: MS and TEM data collection.

M.J.P.: Designed and supervised the study. Provided resources. J.N.Bla.: Supervised the cryo-EM data collection and processing.

L.C.M.M.: Designed and supervised the study. Provided resources. Supported with manuscript structure and writing.

J.Blo.: Designed and supervised the study. Performed the cloning. Did the microscopy. Analyzed and interpreted the data. Created the figures. Wrote the manuscript (lead).

## Declaration of interests

The authors declare no competing interests.

## STAR methods

### *Ulva* growth and cultivation

All transgenic *Ulva* lines were generated using the *Ulva mutabilis/compressa* Føyn “*slender*” lab strain, a descendant of the original isolates from the South Atlantic coast of Portugal.^50^ *Ulva* is maintained as haploid gametophytes and cultivated in a tissue culture chamber under long-day conditions (16-h light:8-h dark; 21°C; approx. 75 mE m^-2^ sec^-1^; Spectralux Plus NL-T8 36W/840/G13 fluorescent lamp) in standard Petri dishes (150x25 mm, SPL Life Sciences) containing 50-100 mL synthetic *Ulva* Culture Medium (UCM).^141,142^ *Ulva* was grown and parthenogenetically propagated as previously described.^143^

### Identification of pCCM-related genes

We identified orthologous genes using the Integrative Orthology Viewer of PLAZA^144,145^ using *Chlamydomonas* gene IDs as input. Each *Ulva* ortholog was identified using at least one evidence: Orthologous gene families (ORTHO), Tree-based orthologs (TROG) and/or Best-Hits-and-Inparalogs (BHIF; **Table S1; Table S2**).

FLIPPer^19^ was used with default settings (**Fig. S7**). To identify RBM-containing proteins using the MEME suite, an initial motif was built using the Multiple Em for Motif Elicitation (MEME) program with the following settings (-mod anr -nmotifs 3 -minw 6 -maxw -30 -objfun classic -markov_order 0). A subsequent search of the *Ulva* proteome using the Motif Alignment and Search Tool (MAST) program was used to identify proteins with similar motifs, which were then compiled and used for a final MEME search to identify high confidence motifs (motif P-value < 1e-10; **Fig. S15**).

### Molecular cloning

Vector assembly is based on the GreenGate cloning system and our existing molecular toolkit for *Ulva.*^25,146^ Sequences for entry modules were obtained via gene synthesis (Neochromosome) or PCR amplification of the target DNA sequence using primers that contain c. 20-bp overlap with the entry module using Q5 polymerase under standard conditions and including the High GC Enhancer (NEB; **Table S7**). All *Ulva* gene sequences were cloned from genomic DNA, isolated via CTAB^147^ or OmniprepTM for Plant (G-Biosciences).

After gel electrophoresis, the amplicons were gel purified (Zymoclean Gel DNA Recovery Kit; Zymo Research) and mixed with the predigested (BsaI; NEB) entry plasmid and NEBuilder® HiFi DNA Assembly Master Mix (NEB) for Gibson assembly. After assembly (15–60 min at 50°C), the cloning reactions were transformed into chemically competent DH5a *Escherichia coli* cells and plated on LB + 100 µg ml^-1^ Carbenicillin. PCR-mediated silent mutations were introduced to remove internal BsaI recognition sites in 75 genes (**Table S2; Table S7**). The destination vector pGG-pUM140_0016-BE-NOST-PIBT3 was previously generated.^25^

For a Golden Gate reaction, 100 ng of each entry and destination vector were assembled in one reaction mix containing 10 U BsaI-HF-v2 (NEB), 200 U T4 Ligase (NEB), 1 mM ATP (Thermo Scientific), and 1x Cutsmart buffer (NEB). The Golden Gate reaction was run for 30 cycles (16°C for 3 min and 37°C for 3 min), followed by 50°C for 5 min and 80°C for 5 min. Approximately 2.5 µl of the assembly mix was transformed into DH5a *E. coli* cells and plated on LB + 25 µg ml^-1^ Kanamycin. Vectors were validated using Sanger sequencing (Mix2seq; Eurofins) or Oxford Nanopore whole-plasmid sequencing (Plasmidsaurus or Eurofins) and restriction digest, typically using NcoI (Promega), PstI (Promega), or XhoI (NEB). Vector sequences generated in this study (**Table S7**) are available at https://vectorvault.vib.be/ulva-mutabilis.

Untagged SUPA1 and R4 sequences were synthesized by TWIST bioscience in pET28a expression vectors. Mutant variants of R4 were created using *in vivo* assembly (IVA)^148^ in which primers contained nucleotide substitutions to introduce the appropriate residue substitutions (**Table S8**).

### *Ulva* transformation

*Ulva* gametes were transformed using the established Polyethylene glycol (PEG)-mediated protocol.^59,142^ For each transformation, 3-5 µg of DNA was used per plasmid. Plasmids were purified using a GeneJet Plasmid Miniprep kit according to manufacturer’s recommendations (Thermo Scientific). 50 µg mL^-1^ Phleomycin (Invivogen) was added 72 h after transformation.

Resistant germlings develop parthenogenetically from the transformed gametes. One additional volume of 40 mL selective medium was added on top after two weeks. After three to four weeks of cultivation, resistant individuals were imaged using confocal microscopy. For each transformation experiment, no-vector controls were included to confirm proper selection.

### Confocal imaging

Thalli of resistant individuals were used for imaging. Detailed imaging of cells was performed using an Olympus Fluoview (FV1000) confocal microscope with FluoView software (FV10-ASW 4.2). The following excitation and emission settings were used: YFP, 515 nm excitation with 530–545 nm emission and chlorophyll, 559 nm excitation with 650–750 nm emission. Laser intensity was adapted depending on expression variation between individuals. 3-29 (average 10) independent transformants were imaged per construct (**Table S2**). Individuals were imaged with a 60X W objective using 1x and 3x digital zoom. All observed protein localizations (**Table S2; Fig. S3**) were annotated on the *Ulva mutabilis* reference genome on ORCAE: https://bioinformatics.psb.ugent.be/orcae/overview/Ulvmu.

We determined internal mixing of stably transformed *Ulva* expressing SUPA1-YFP or RbcS1-YFP using Fluorescence Recovery After Photobleaching (FRAP). Imaging was done on a PerkinElmer Ultraview spinning-disk system, attached to a Nikon Ti inverted microscope, and operated using Volocity software (V2.1). Images were acquired on an ImagEM CCD camera (Hamamatsu C9100-13) using frame-sequential imaging with a 100x O NA 1.2 objective. FRAP was done on the spinning disk system using the Ultraview PhotoKinesis unit. Bleaching was performed using a 2×2 µm ROI using 100% of the 488 laser for 3s, images were acquired 5 seconds before and every 10 seconds up to 400 seconds after bleaching. Analysis of FRAP experiments was performed according to Kumar et al.^102^, in which individual replicates were fitted to obtain half-life recovery values.

### Generation of SUPA1 CRISPR lines

The mutant lines were generated using Cas9 RNP-mediated CRISPR targeted insertion^26^ with some modifications. Two guide RNAs targeting SUPA1 were designed with Geneious Prime software (version 2023.0.1) using the nuclear genome^149^ as an off-target database (**Table S8**). The purified crRNAs, trans-activating crRNA (tracrRNA), and HiFi S.p. Cas9 Nuclease V3 (Integrated DNA Technologies) were assembled into a RNP complex with a Cas9:gRNA 1:1 molar ratio in NEBbuffer 3.1 (New England Biolabs). As a donor, we amplified the blasticidin resistance cassette (pUM056_0044-BSD-tRbcS)^26^ by PCR using Q5 polymerase (NEB) and included 25 bp homology arms to the target sites (**Fig. S8**). The amplicon was purified (DNA Clean and Concentrator-25 kit; Zymo Research). 8 µl of RNP solution at a concentration of 1 µg/µl of Cas9 was co-transfected with 5 µg of BSD donor DNA in gametes of transgenic lines expression RbcS1-YFP. Putative mutant individuals were selected using 50 µg ml^-1^ Blasticidin (Invivogen).

After three weeks, multiple resistant individuals were observed. To confirm the insertion of the donor DNA at the target site, genomic DNA was extracted from selected individuals via CTAB.^147^ Primers were designed to amplify both the target site and either the left or right homology arms of BSD (**Table S8; Fig. S8**). The amplicons were Sanger sequenced (Mix2seq; Eurofins).

### Transmission Electron Microscopy

Thalli of one-month old *Ulva* were cut into smaller pieces and immersed in a fixation solution of 4% paraformaldehyde and 2.5% glutaraldehyde in 0.1M sodium-cacodylate buffer for 3h, rotating at room temperature. The solution was later replaced with fresh fixative and samples were left rotating overnight at 4°C. After washing with buffer, samples were post-fixed in 1% OsO4 with K3Fe(CN)6 in 0.1 M sodium-cacodylate buffer, pH 7.2, on ice. After washing with ddH2O, samples were placed in 1% uranyl acetate in water for 1h, in the dark, rotating at room temperature. Subsequently the samples were dehydrated through a graded ethanol series, followed by embedding in Spurr’s resin.^150^ After trimming, ultrathin sections of a gold interference colour (90 nm) were cut using an ultra-microtome (Leica EM UC7) and collected on formvar-coated copper slot grids. Sections were post-stained with 1% uranyl acetate and 3% lead citrate in a Leica EM AC20. They were imaged with a JEM 1400plus transmission electron microscope (JEOL, Tokyo, Japan) operating at 80 kV.

### Gene expression analysis

As an approximation of RNA expression, the FKPM values over the different timepoint analyzed in ^62^ were averaged. For mapping RNA sequencing reads to RbcS1 and RbcS2, RNA sequences were obtained from NCBI (ERR2722103; ERR2722112-ERR2722114; ERR27221-ERR2722135) using SRA Toolkit-3.2.0. Both paired reads (46.2 Gbp) and unpaired reads (33.1 Gbp) were mapped to the CDS of the genes using BWA-0.7.18^151^ using default settings. Files were processed, mapped reads extracted, and coverage calculated using SAMtools-1.21.^152^

### Whole cell and pyrenoid-enriched proteomics

Thalli of *Ulva* sp. were collected from an intertidal zone at Robin Hood’s Bay, North Yorkshire and maintained in natural seawater sparged with ambient air under ∼20 µE light until processing (2 days). For lysis, material was frozen in liquid nitrogen and ground to a fine powder in a pestle and mortar pre-cooled with liquid nitrogen. For whole cell mass spectrometry, the powder was dissolved in 5% SDS, 100 mM TEAB, pH 8.5 buffer and sonicated (3 min processing time, 3 s pulses at 30% amplitude using a micro-tip and Misonix S-4000 sonicator). The lysate was centrifuged at 30,000 *g* for 30 minutes at 4 °C and the supernatant extracted for analysis. For mass spectrometry samples were digested with trypsin using a sTrap-mediated digestion procedure. Peptides were analysed by LC-MS, with elution from a 15 cm Performance C18 column over a 30 SPD EvoSep gradient. Spectra were acquired in PASEF-DIA mode using a Bruker timsTOF HT mass spectrometer. Data were searched using Spectronaut against the *Ulva mutabilis* chloroplast and nuclear proteome databases, appended with common proteomic contaminates. Results were filtered to protein q<0.01 and a minimum of two peptides per accepted protein. Summed DIA intensities were extracted using Spectronaut. Samples were analysed in triplicate, and the summed intensities across replicates are reported (**Table S4**).

For pyrenoid-enrichment, the powder following grinding was dissolved in 2% formaldehyde in 30 mM HEPES-KOH pH 8.0 and stirred for 30 minutes at room temperature. Fixing was quenched by addition of and equal volume of 2 M Tris-HCl pH 8.0. The resulting mixture was passed through Miracloth to remove unbroken thallus material and centrifuged at 2,500 *g* for 20 minutes at room temperature. The pellet was resuspended in 2 mL of pyrenoid isolation buffer (50 mM Tris-HCl, 0.2 mM EDTA, 0.5% (w/v) Triton X-100, pH 8.0) and the pyrenoid fraction enriched by centrifugation at 20,000 *g* for 1 minute at room temperature. The resulting pellet was resuspended in 1 mL of pyrenoid isolation buffer and loaded onto a 9 mL 100% Percoll cushion. Following centrifugation, the pellet at the bottom of the cushion was washed twice with 1 mL of pyrenoid isolation buffer. Proteins were eluted from the pyrenoid fraction with 5% SDS, 100 mM TEAB, pH 8.5 buffer. The total protein concentration in the elution was determined using a Pierce BCA protein assay kit and normalized to the protein concentration in the ‘top’ fraction of the Percoll gradient for analysis (**Fig. S10**). For mass spectrometry, samples were digested as before, and peptides were separated over a 60 SPD EvoSep gradient with elution from an 8 cm Performance C18 column. Mass spectra were acquired in PASEF-DIA mode using a Bruker timsTOF HT mass spectrometer. Data were searched using Spectronaut against the same database and identifications were filtered as before. Relative abundance testing was performed using limma via FragPipe-Analyst. Sample minimum imputation was applied and the Hochberg and Benjamini approach was used for multiple test correction. Samples were analyzed in triplicate. One lysate sample was excluded from the differential abundance analysis due to detection of ∼⅓ of the proteins compared to the other two replicates (**Table S4**).

### Purification of Rubisco from *Ulva*

Thalli of *Ulva intestinalis* were collected from Staithes, North Yorkshire, UK and processed within two hours of collection. ∼200 g wet weight material was lysed in a blender in 200 mL of Rubisco lysis buffer (50 mM Tris-HCl, 100 mM NaCl, 0.5 mM EDTA, 5 mM DTT at pH 8.0) for 15 minutes at 4 °C. The lysate was centrifuged at 4,000 × *g* for 15 minutes twice to remove unbroken material and cell debris, after which the solution was centrifuged at 50,000 × *g* for 30 minutes at 4 °C. 200 mg mL^-1^ ammonium sulphate was added to the supernatant and stirred for 2 hours at 4 °C before centrifugation at 30,000 × *g* for 10 minutes at 4 °C. A further 190 mg mL^-1^ ammonium sulfate was added to the supernatant and stirred as before. The protein was pelleted and resuspended in anion exchange buffer A (50 mM Tris-HCl, 50 mM NaCl, 1 mM DTT at pH 8.0) and dialyzed overnight against the same buffer at 4 °C. The solution was centrifuged after dialysis at 90,000 × *g* for 30 minutes at 4 °C to remove remaining membrane fragments, before application to an XK-26 column packed with SOURCE 15Q resin. A step elution to 15% of anion exchange buffer B (50 mM Tris-HCl, 1 M NaCl, 1 mM DTT at pH 8.0) was used to elute non-specific proteins, after which a gradient to 37.5% of anion exchange buffer B over 20 column volumes was used to elute Rubisco. Rubisco fractions were concentrated using 10,000 kDa MWCO concentration columns (Merck) and subjected to size exclusion chromatography using a Superdex 200 PG 26/600 column and size exclusion buffer (50 mM Tris-HCl, 50 mM NaCl, 1 mM DTT, 5% glycerol (w/v) at pH 8.0). Rubisco fractions were analyzed by SDS-PAGE, concentrated, snap-frozen and stored at −70 °C prior to use.

### Purification of SUPA1 variants from *E. coli*

*E. coli* BL21(DE3) cells harboring relevant plasmids (Twist Bioscience) were grown to OD_600_ of 0.5-0.8 Luria Broth or ^15^NH_4_Cl minimal media for the NMR samples and induced with 1 mM IPTG overnight at room temperature. Cell pellets were lysed by sonication in denaturing lysis buffer (6 M urea, 500 mM NaCl, 50 mM Tris-HCl, 25 mM imidazole), before centrifugation at 50,000 × *g* for 30 minutes at 4 °C. The supernatant was applied to 5 mL HisTrap FF columns (Cytiva) equilibrated with lysis buffer. Unbound proteins were removed by washing with lysis buffer prior to refolding on the column by washing with HisTrap buffer A (50 mM Tris-HCl, 500 mM NaCl, 25 mM imidazole at pH 8.0). Bound protein was eluted with a gradient to HisTrap buffer B (50 mM Tris-HCl, 500 mM NaCl, 500 mM imidazole at pH 8.0) over 20 column volumes. Fractions containing SUPA1 variants were pooled and cleaved overnight with either Thrombin from bovine plasma or TEV protease. Following cleavage, samples were buffer-exchanged into HisTrap buffer A and a reverse nickel affinity purification was performed, in which the flowthrough was collected. The collected fractions were concentrated and subjected to size exclusion chromatography using a Superdex 75 PG 16/600 column equilibrated with linker size exclusion buffer (50 mM Tris-HCl, 500 mM NaCl, 5% glycerol (w/v) at pH 8.0). For the mEGFP-tagged SUPA1, the protein was purified under non-denaturing conditions and without the cleavage and reverse IMAC.

### Droplet sedimentation assays

Droplet microscopy and sedimentation assays were performed in volumes of 5 µL in a buffer of 50 mM Tris-HCl, 50 mM NaCl at pH 8.0. Rubisco was added first in all cases and droplets were allowed to form for 15 minutes at room temperature prior to sedimentation at 10,000 *g* for 10 minutes at 4 °C, before analysis by SDS–PAGE. Band intensity was quantified in Fiji.

### Single-particle cryo-EM data collection and image processing

*Ulva* Rubisco and the R4 fragment were mixed in a buffer of 25 mM Tris, 25 mM K acetate pH 8.0, 50 mM sorbitol, 4 mM Mg, 2 mM CaCl2 at a final concentration of 0.5 μM and 75 μM respectively and incubated at room temperature for 10 minutes. 2.5 µL of sample was applied to R1.2/1.3 Cu 300-mesh grids (Quantifoil) that were treated with air plasma for 120 s at 0.6 Torr of air in Harrick microwave Plasma cleaner PD-001. Grids were blotted for 8 seconds with a blot force of −10 at 4 °C and 95% relative humidity before plunge-freezing in liquid ethane, using an FEI Mark IV Vitrobot system (ThermoFisher).

Data were collected on a 300 kV Krios cryo-electron microscope equipped with a Gatan K3 detector at the Diamond Light Source. Automated data collection was performed using EPU. A nominal magnification of x105,000 and electron fluence of 60 e^−^ Å^−2^ with a calibrated pixel size of 0.825 Å was used during collection in which each exposure was 2.6 seconds. The 100 μm objective aperture was inserted, the C2 aperture was 70 μm, and energy filter slit width 20 eV. A range of defocus values were used (−1.8, −1.6, −1.4, −1.2, −1.0, −0.8 μm).

Relion5 was used for data processing and 3D reconstruction.^153,154^ 3,200 micrographs were used, each consisting of 194 frames, and during motion correction the frames were grouped in batches of 10 for alignment. MotionCor2 was used in Relion for motion correction before CTF estimation with CTFFIND4^155^ assuming a spherical aberration of 2.7 mm. 1.8 M particles were autopicked and subjected to 2D classification following extraction with 4x binning. Particles from Rubisco classes were extracted with 2x binning and used for *ab initio* reconstruction. The resulting map was used as a reference for 3D classification with C1 symmetry. A single 3D class was used for auto-refinement with D4 symmetry (462,744 particles), after extraction with no binning. CTF refinement and Bayesian polishing were performed in Relion^156^ and a 15 Å low-pass filtered mask, expanded by 2 pixels with a soft edge of 6 pixels was used for solvent masking and resolution estimation. The final map has EMD ID 58973.

### Single-particle cryo-EM model building, fitting and refinement

Initial coordinates were built into the map using AlphaFold3^157^ generated model and the sequences of the Rubisco large and small subunits and that of R4. Rigid fitting with Fourier filtering was performed in Moorhen (www.moorhen.org). The model was refined in Phenix^158^, ServalCat^159^ and ISOLDE^160^ and Moorhen and validated using MolProbity.^161^ Figures were created in Moorhen and molecular contacts assessed with PDBePISA.^162^ The final model has been deposited on the PDB with id: 32LL.

### NMR spectroscopy

Spectra were recorded as described in ^19^, with the exception that the data was collected at either 4 °C or 24 °C and with R4 protein concentration of 1.4 mM.

### Native PAGE gel-shift assay

Rubisco and R4 variants were mixed in 5 µL volumes containing 0.5 µM Rubisco and 26 µM of each variant and incubated at room temperature for 30 minutes in a buffer of 50 mM Tris-HCl, 500 mM NaCl. 1.6 μL of loading buffer (80mM Tris-HCl pH 8.0, 200mM NaCl, 40% glycerol) was added before loading onto 4–20% Mini-PROTEAN TGX gels. Electrophoresis was completed for 4h at 100V at 4°C. The solubility tags were not removed for these experiments, allowing visualization of the unbound R4 fragment in the gel.

### Native Mass Spectrometry of Linker bound to Rubisco

Samples were all buffer exchanged in 150 mM Ammonium Acetate at pH 6.8 using Bio-Rad Micro Bio-Spin P6 columns (Bio-Rad, Hercules, CA, USA). Rubisco concentration was kept at 3.5 μM for all experiment and R4 fragment concentration varied accordingly from 3.5 μM to 28 μM. Measurements were carried out using a Waters Synapt G2-Si mass spectrometer (Waters Corporation, Manchester, UK) equipped with a nano-electrospray ionization source. Nano electrospray tips were pulled in house from thin-walled borosilicate glass capillaries (i.d. 0.78 mm, o.d. 1.0 mm) (Sutter Instrument Co., Novato, CA, USA) using a flaming/brown micropipette puller (Sutter Instrument Co., Novato, CA, USA). A positive potential of 1.3–1.5 kV was applied to the solution using a thin platinum wire (d. 0.125 mm) (Goodfellow, Huntingdon, UK). Other non-default instrument settings include: sampling cone voltage 150–200 V, source offset voltage 150 V, collision voltage 10 V, trap gas flow 6-8 ml min−1, source temperature 40 °C.

Data analysis and relative species estimation were performed using MassLynx (Waters Corporation) and multi-bi-gaussian deconvolution of the spectra using MsMBG-Fit (https://github.com/cdegut/MsMBG-Fit).

### Isothermal Titration Calorimetry (ITC)

Isothermal titration calorimetry (ITC) experiments were performed using a MicroCal Auto-iTC200 instrument (Malvern Panalytical). All samples were dialyzed against the same buffer consisting of 50 mM Tris-HCl, 50 mM NaCl, and 0.5 mM TCEP at pH 8.0. The sample cell was loaded with 6.5 μM Rubisco, and the syringe was filled with 300 μM of the respective linker samples. Titrations were carried out at 25 °C. Raw thermograms were integrated using MicroCal PEAQ-ITC Analysis software, and the resulting binding isotherms were fitted using an independent-site binding model.

## Supplemental Information

The supplemental information for this manuscript consists of 8 tables and 16 figures.

**Table S1.** Identification of *Chlamydomonas* orthologous genes in *Ulva*.

**Table S2.** Summary of cloning and localization.

**Table S3.** Conservation of localization between *Chlamydomonas* and *Ulva*. **Table S4.** Summary of whole cell (F103) and purified pyrenoid (F107) proteomic experiments.

**Table S5.** Conservation of *Chlamydomonas* pyrenoid-localized proteins in *Ulva*.

**Table S6.** Conservation of SUPA1.

**Table S7.** Complete list of generated vectors (entry, destination and expression) and primer sequences.

**Table S8.** Sequences related to SUPA1 characterization

**Figure S1.** Chloroplast and pyrenoid ultrastructure of *Ulva compressa*.

**Figure S2.** Cloning and transgene expression statistics.

**Figure S3.** Protein localizations of tagged *Ulva* pCCM-related genes.

**Figure S4.** Comparison of observed and predicted localizations.

**Figure S5.** Identification of *Ulva* BST4 functional analog.

**Figure S6.** RbcS2 is lowly expressed in *Ulva*. **Figure S7.** FLIPPer analysis of *Ulva* proteins. **Figure S8.** Genotyping of SUPA1 mutant lines. **Figure S9.** Pyrenoid enrichment from *Ulva* biomass.

**Figure S10.** *In vitro* characterisation of SUPA1-Rubisco condensation. **Figure S11.** Single particle cryo-EM data processing and interpretation **Figure S12.** NMR analysis of the 53-residue RBM linker fragment.

**Figure S13.** ITC and native PAGE characterization of SUPA1-R4 binding-deficient mutants.

**Figure S14.** Raw native mass spectrometry data corresponding to Fig. 6N.

**Figure S15.** Identification of RBM-containing proteins.

**Figure S16.** Putative membrane-associated pyrenoid proteins.

## Notes

### Competing Interest Statement

The authors have declared no competing interest.

