## Supplemental Figures for "A spatial atlas of the seaweed CO_2_-fixation machinery reveals a unique Rubisco condensation mechanism"

Supplemental Information

The supplemental information for this manuscript consists of 8 tables and 16 figures.

**Table S1.** Identification of *Chlamydomonas* orthologous genes in *Ulva*.
**Table S2.** Summary of cloning and localization.
**Table S3.** Conservation of localization between *Chlamydomonas* and *Ulva*.
**Table S4.** Summary of whole cell (F103) and purified pyrenoid (F107) proteomic experiments.
**Table S5.** Conservation of *Chlamydomonas* pyrenoid-localized proteins in *Ulva*.
**Table S6.** Conservation of SUPA1.
**Table S7.** Complete list of generated vectors (entry, destination and expression) and primer sequences.
**Table S8.** Sequences related to SUPA1 characterization

**Figure S1.** Chloroplast and pyrenoid ultrastructure of *Ulva compressa*.
**Figure S2.** Cloning and transgene expression statistics.
**Figure S3.** Protein localizations of tagged *Ulva* pCCM-related genes.
**Figure S4.** Comparison of observed and predicted localizations.
**Figure S5.** Identification of *Ulva* BST4 functional analog.
**Figure S6.** RbcS2 is lowly expressed in *Ulva*.
**Figure S7.** FLIPPer analysis of *Ulva* proteins.
**Figure S8.** Genotyping of SUPA1 mutant lines.
**Figure S9.** Pyrenoid enrichment from *Ulva* biomass.
**Figure S10.** *In vitro* characterization of SUPA1-Rubisco condensation.
**Figure S11.** Single particle cryo-EM data processing and interpretation
**Figure S12.** NMR analysis of the 53-residue RBM linker fragment.
**Figure S13.** ITC and native PAGE characterization of SUPA1-R4 binding-deficient mutants.
**Figure S14.** Raw native mass spectrometry data corresponding to Fig. 6N.
**Figure S15.** Identification of RBM-containing proteins.
**Figure S16.** Putative membrane-associated pyrenoid proteins.

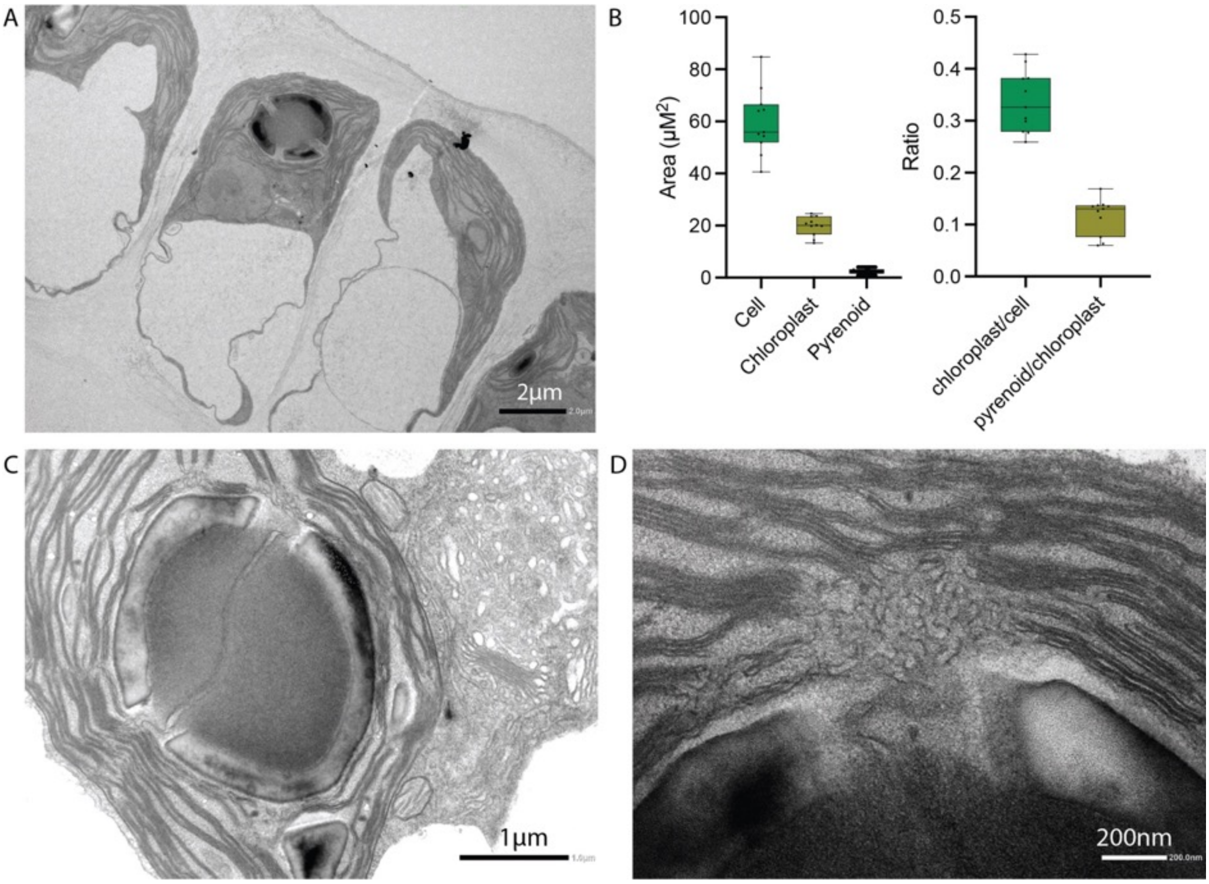


**Figure S1.** Chloroplast and pyrenoid ultrastructure of *Ulva compressa*.

A. Representative image of *Ulva* cells containing a single, cup-shaped chloroplast. Scale: 2 μm.
B. Area of *Ulva* cells (μm$^2$), chloroplasts and pyrenoids (left), the relative size of chloroplast per cell and pyrenoid per chloroplast (right; n=11).
C. Detail of the pyrenoid with a single PTT and surrounded by two starch plates. Scale: 1 μm
D. Detail of the thylakoid convergent zone at the pyrenoid periphery. Scale: 200 nm.

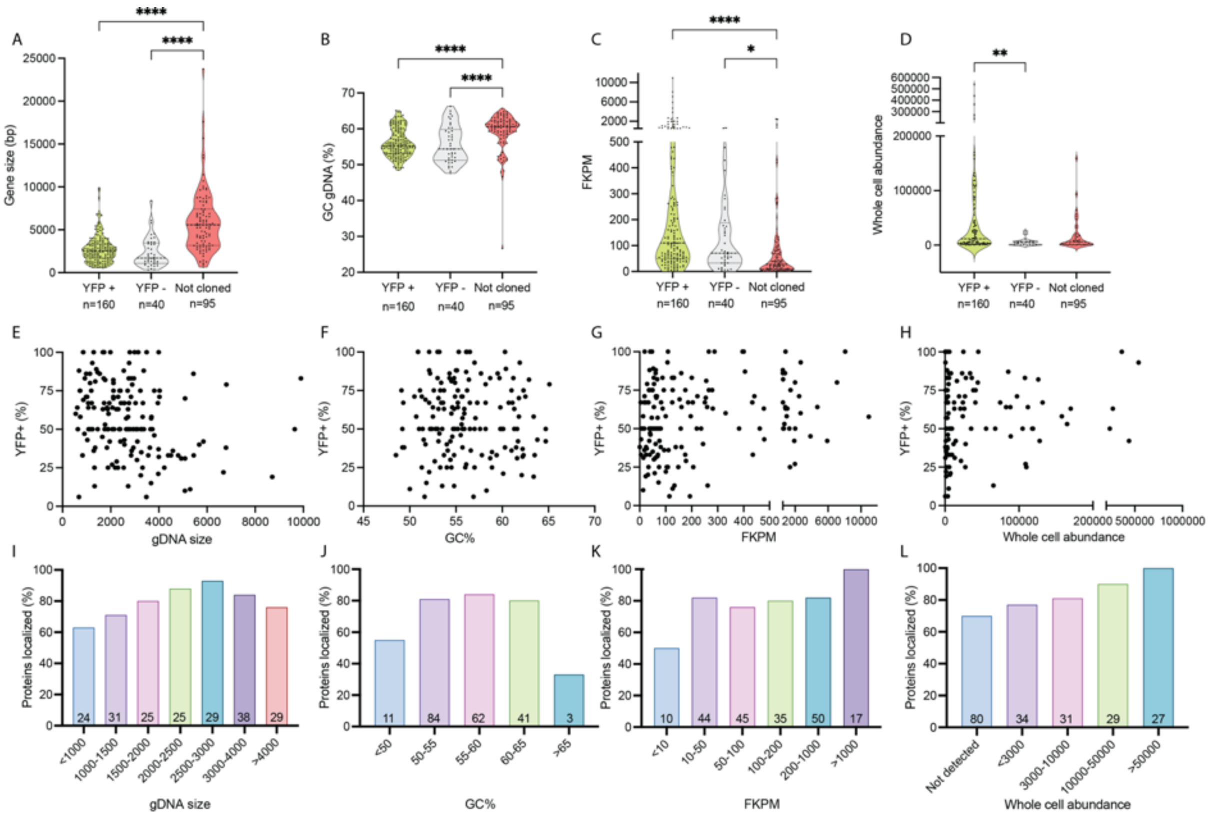


**Figure S2. Cloning and transgene expression statistics.**

Effect of (A) gene size, (B) GC %, (C) expression (expressed in Fragments Per Kilobase of transcript per Million mapped reads (FPKM)) and (D) whole cell protein abundance on PCR amplification success. Genes that were cloned are further subdivided in constructs where protein localization data was obtained (YFP+) or not (YFP-). Significant differences (Kruskal-Wallis test): p>0,05 *, p>0,01 ** and p>0,0001 ***.

Effect of (E) gene size, (F) GC %, (G) gene expression and (H) whole cell protein abundance on number of independent transformants with YFP signal, indicated as % of screened individuals.

Effect of (I) gene size, (J) GC %, (K) gene expression and (L) whole cell protein abundance on number of proteins localized, indicated as % of proteins where localization was observed relative to all proteins within certain bins. Number of proteins per bin is indicated in each bar plot. See also **Table S2**.

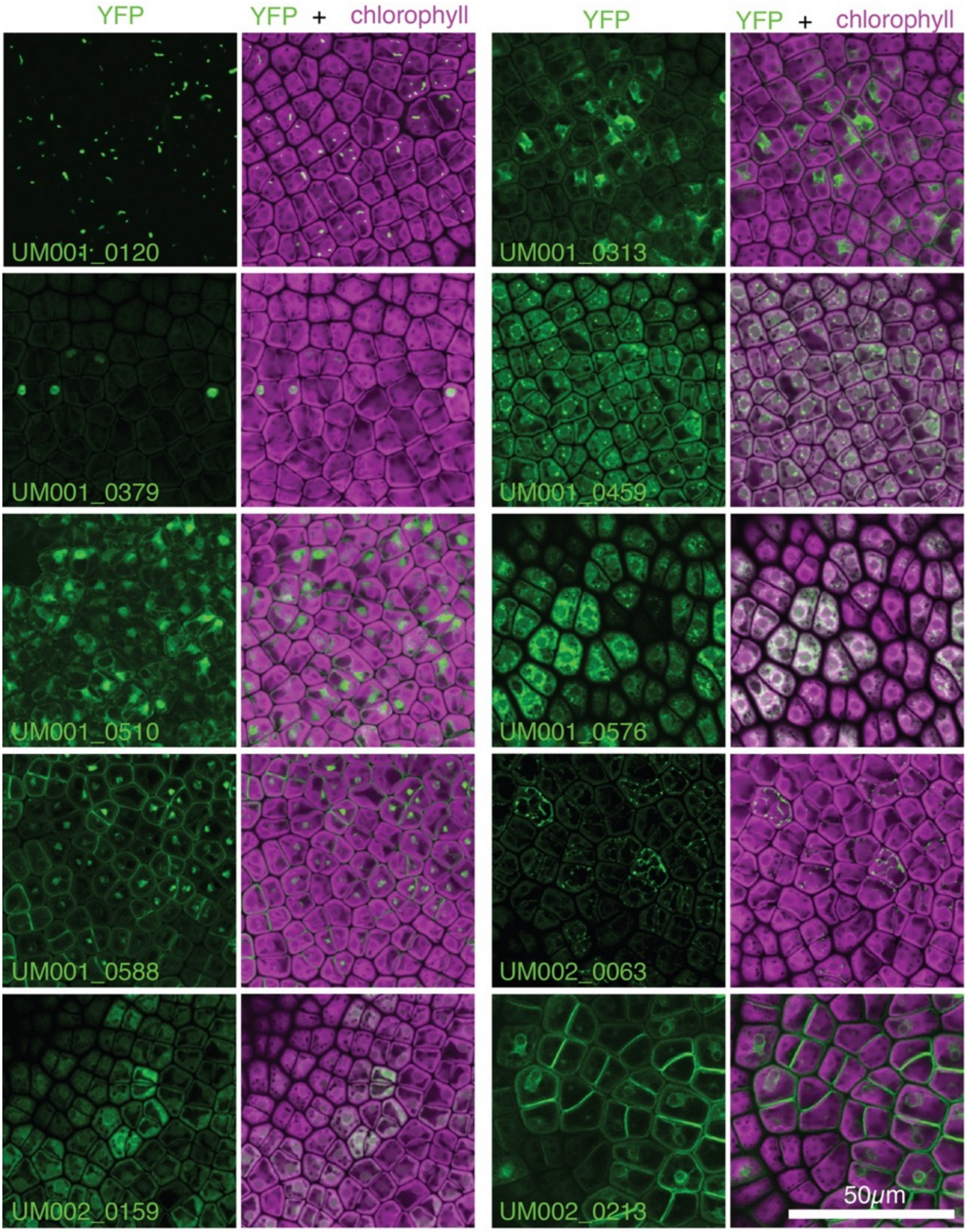


Figure S3 1/16

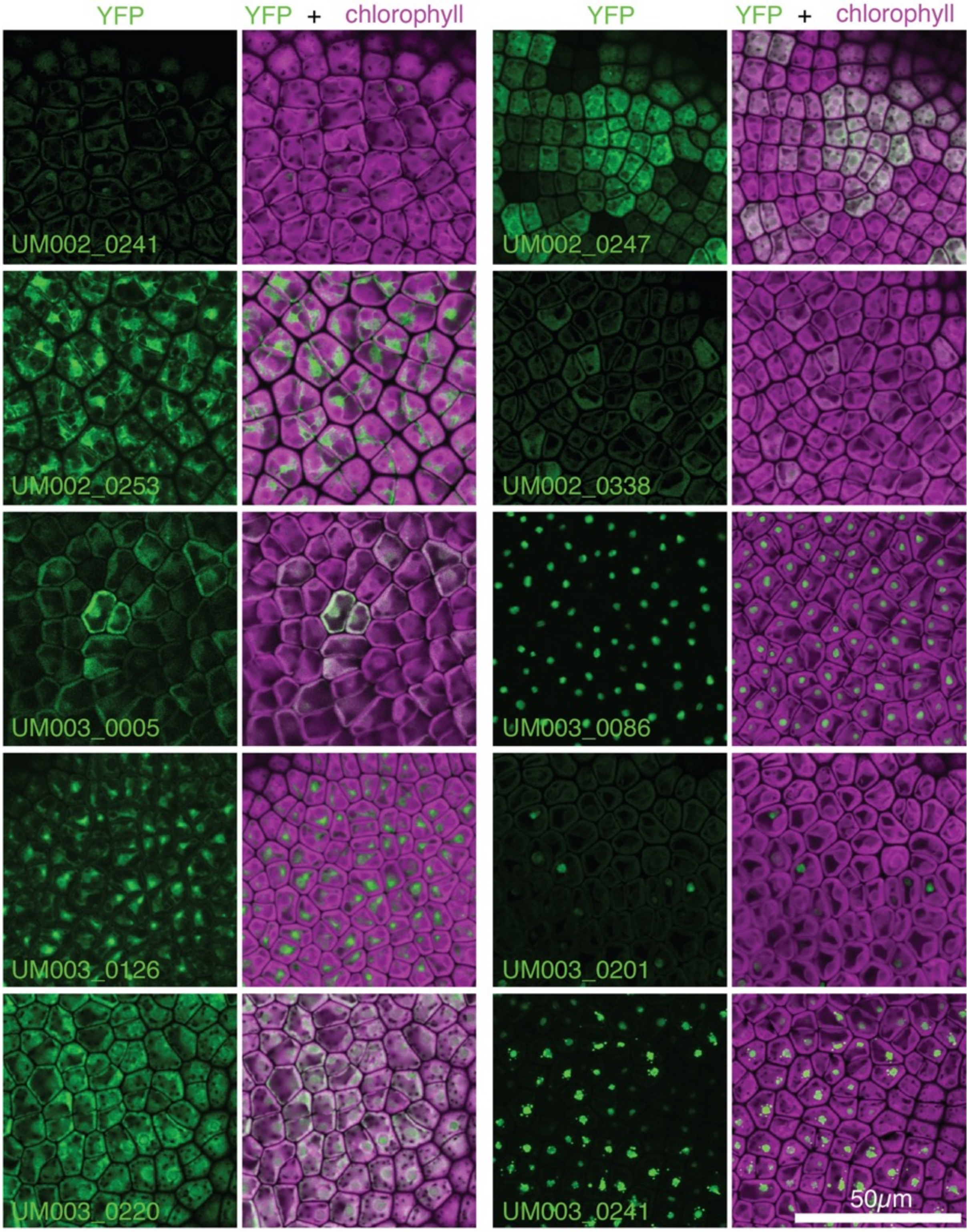

Figure S3 2/16

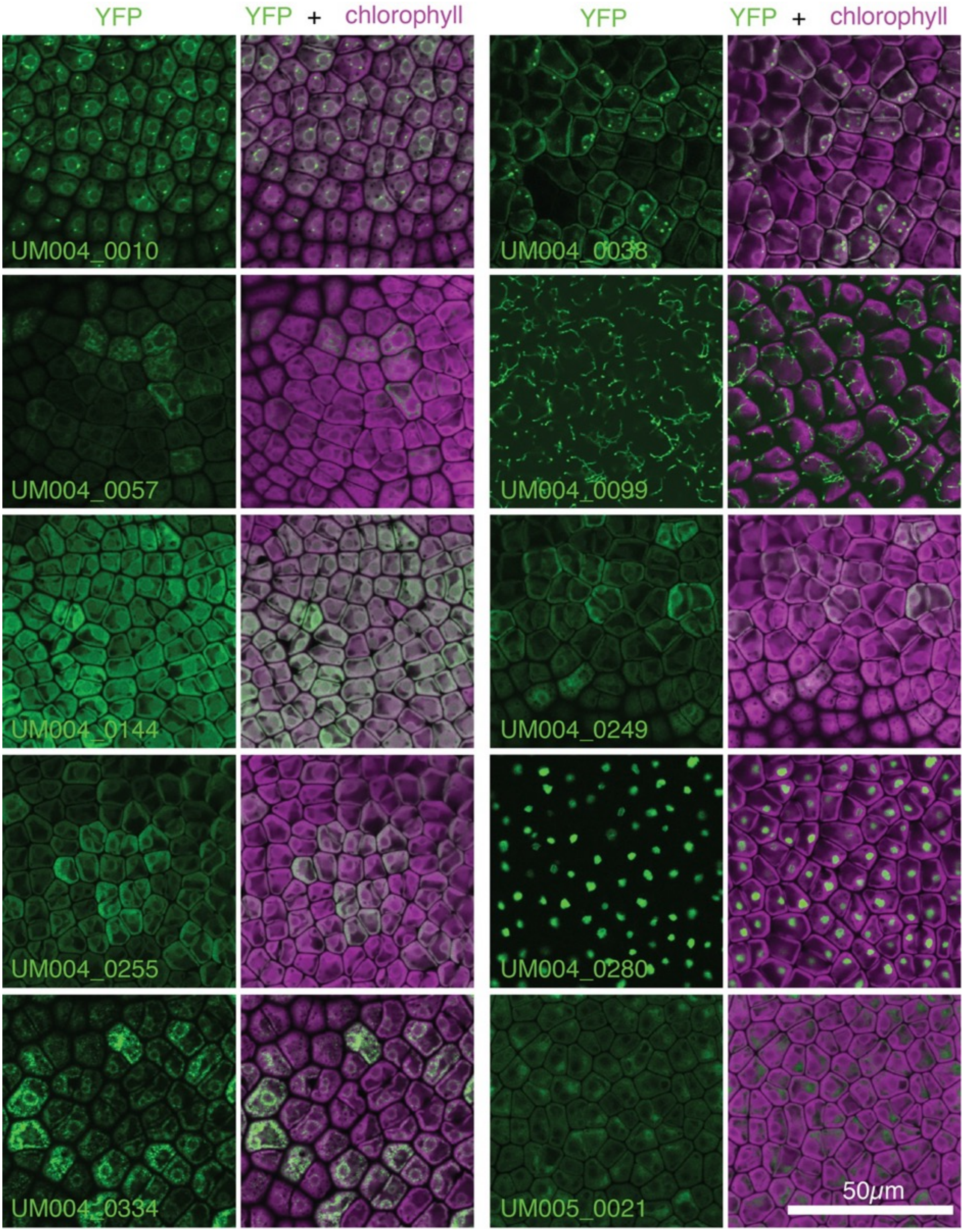


Figure S3 3/16

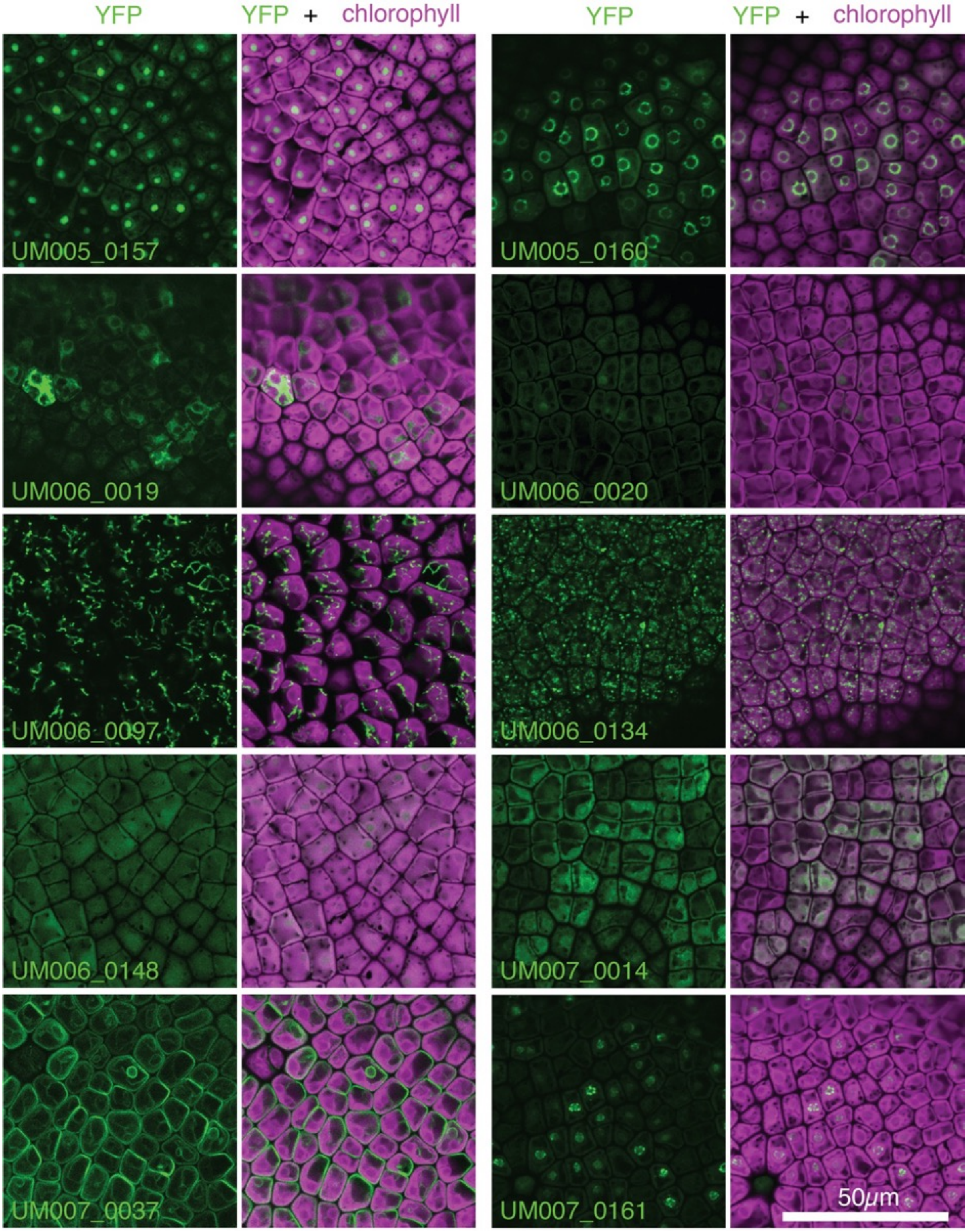


Figure S3 4/16

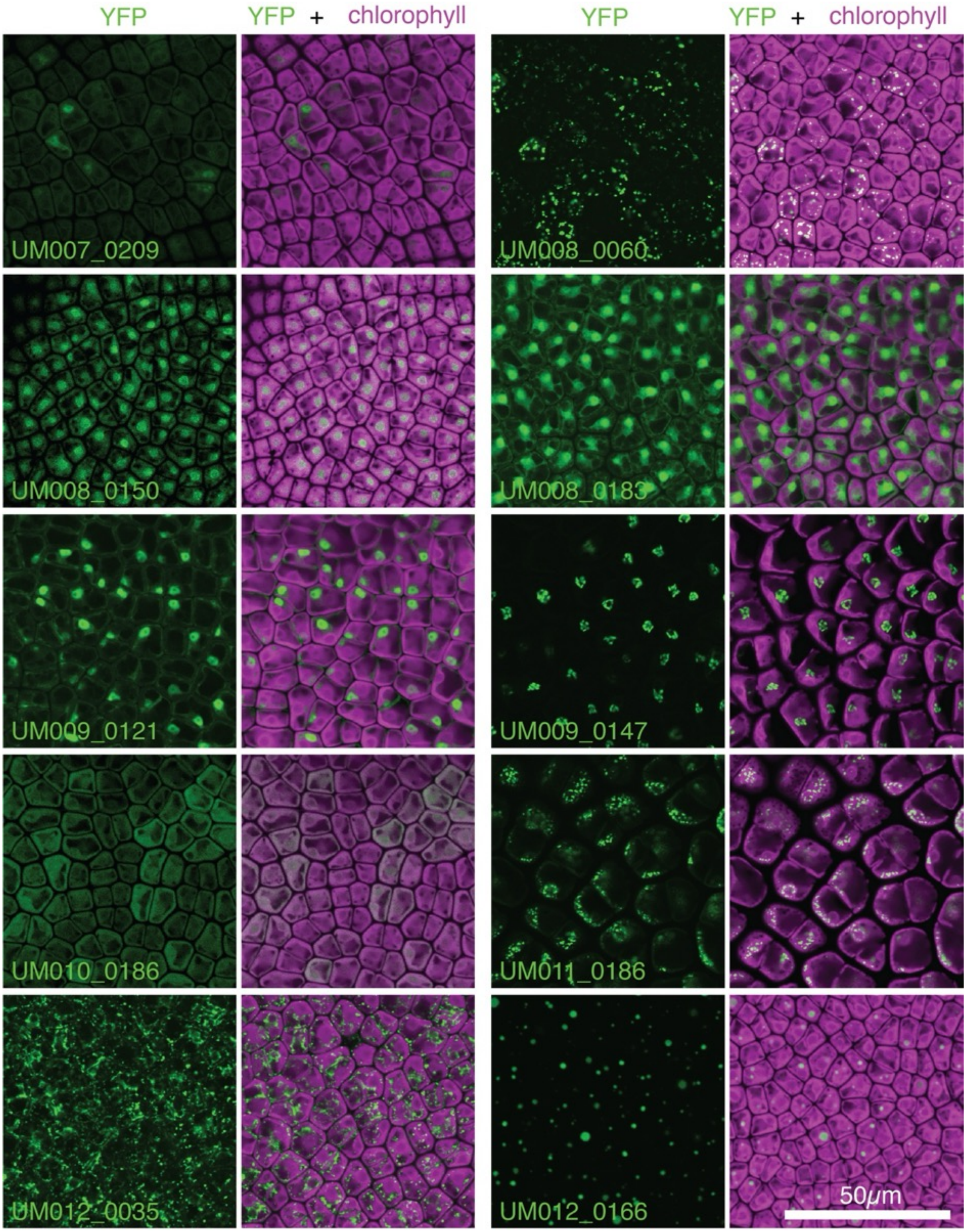


Figure S3 5/16

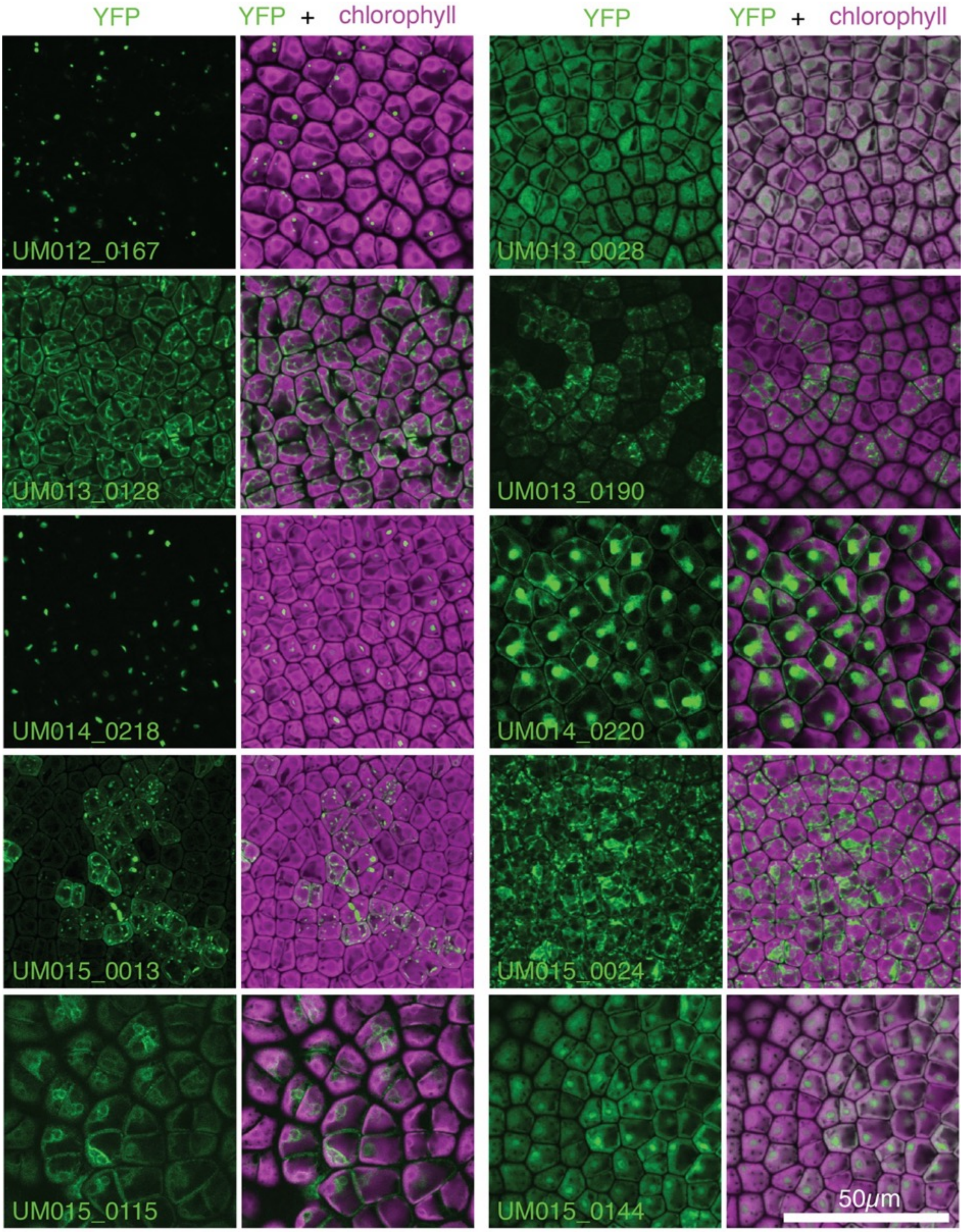


Figure S3 6/16

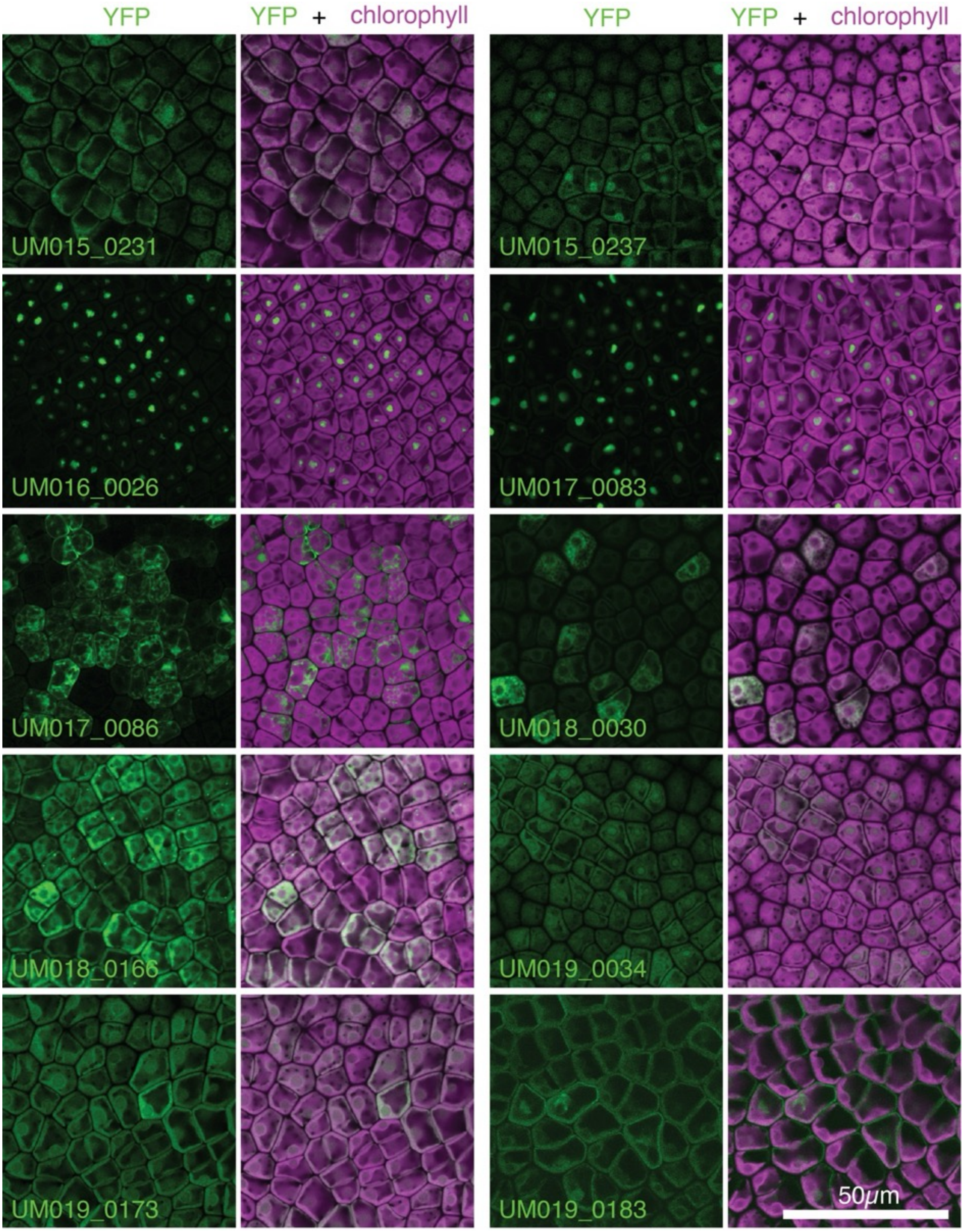


Figure S3 7/16

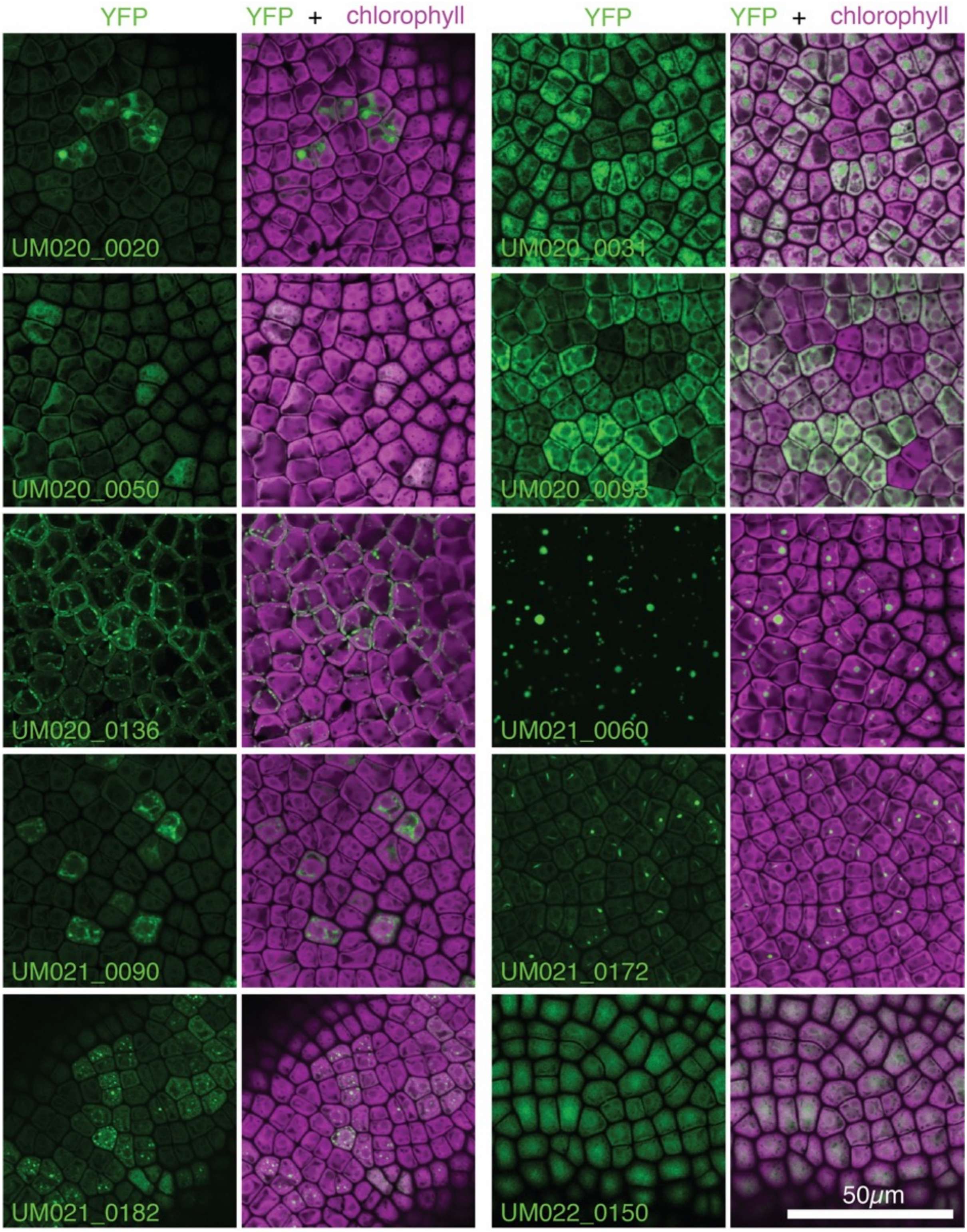


Figure S3 8/16

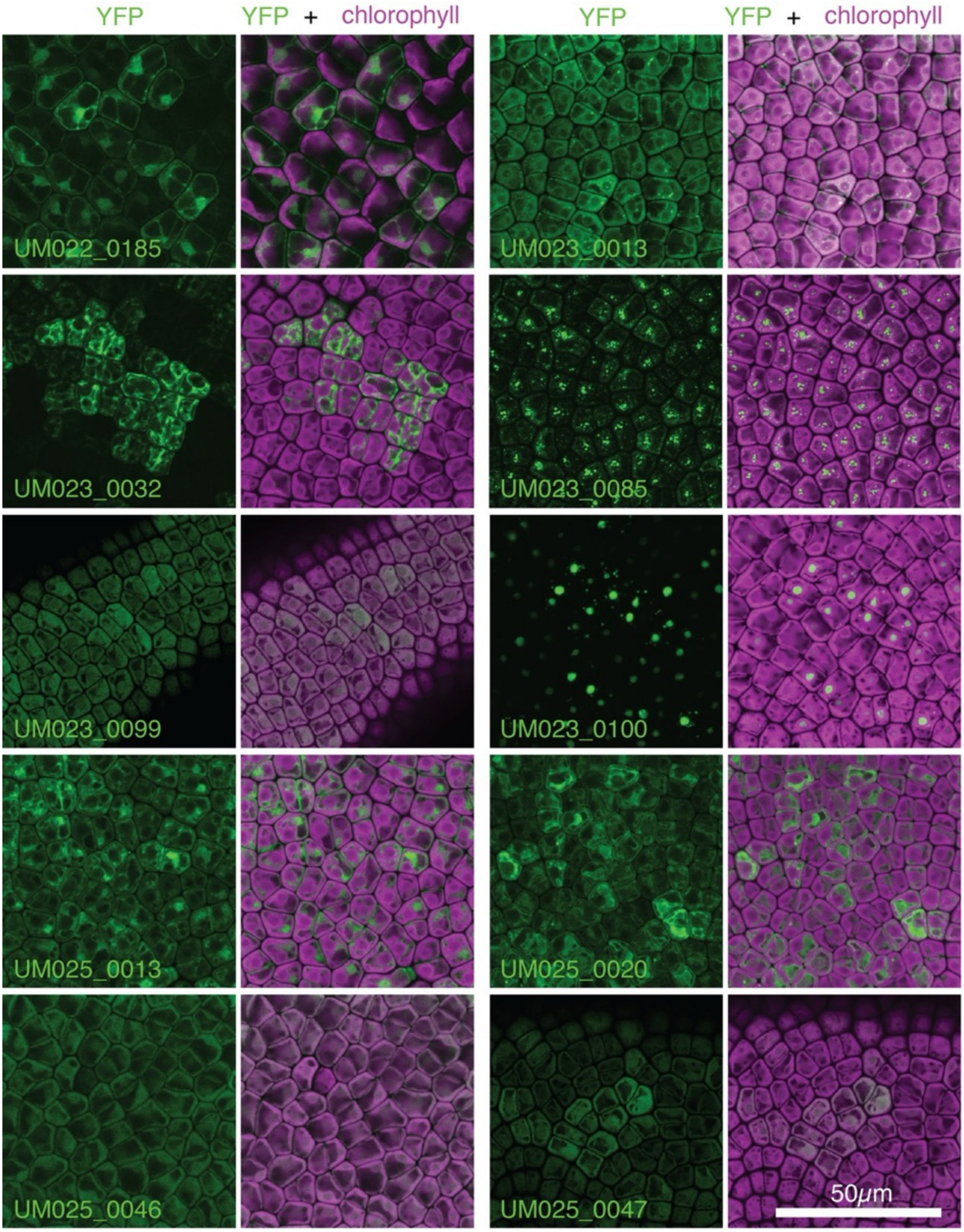


Figure S3 9/16

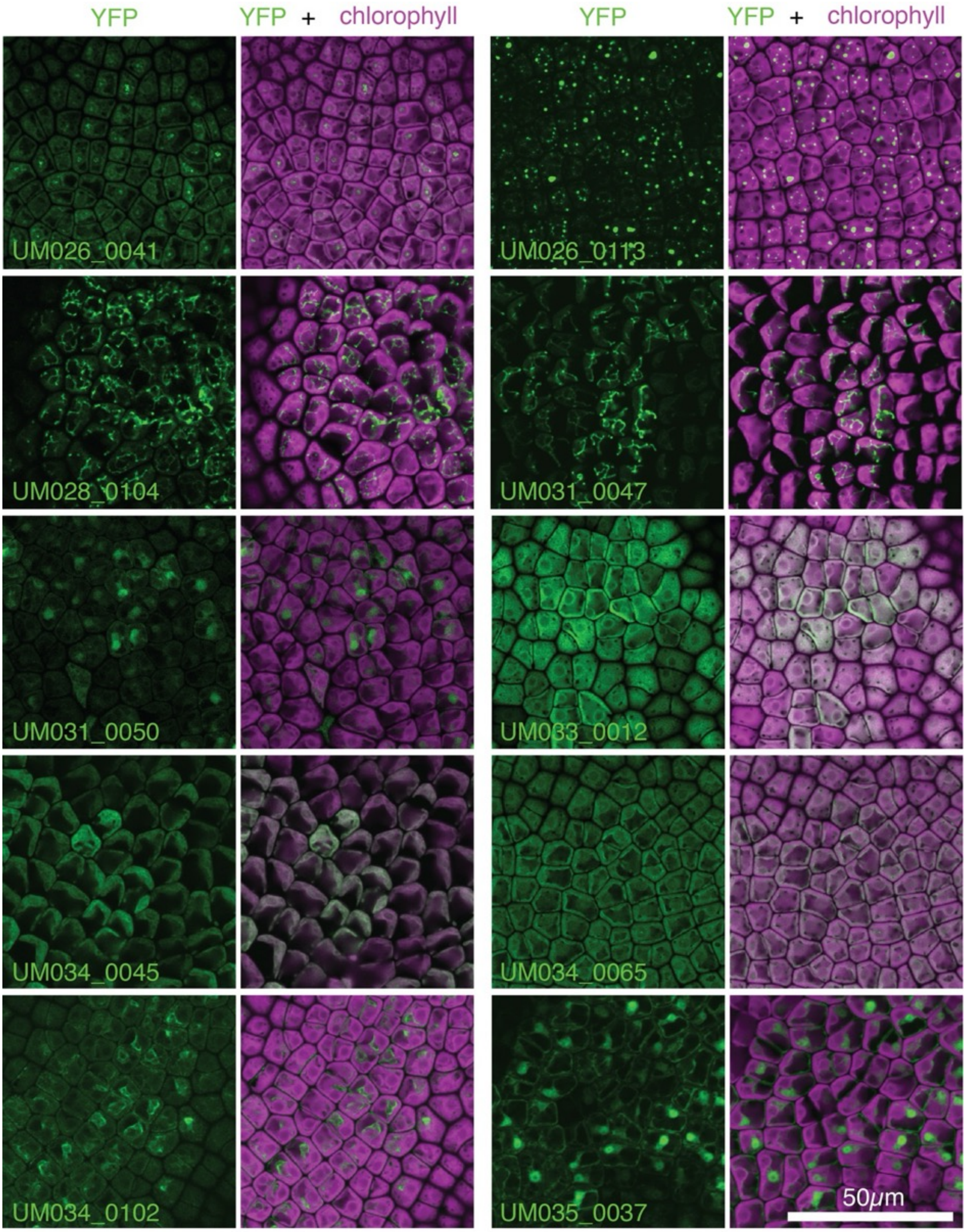


Figure S3 10/16

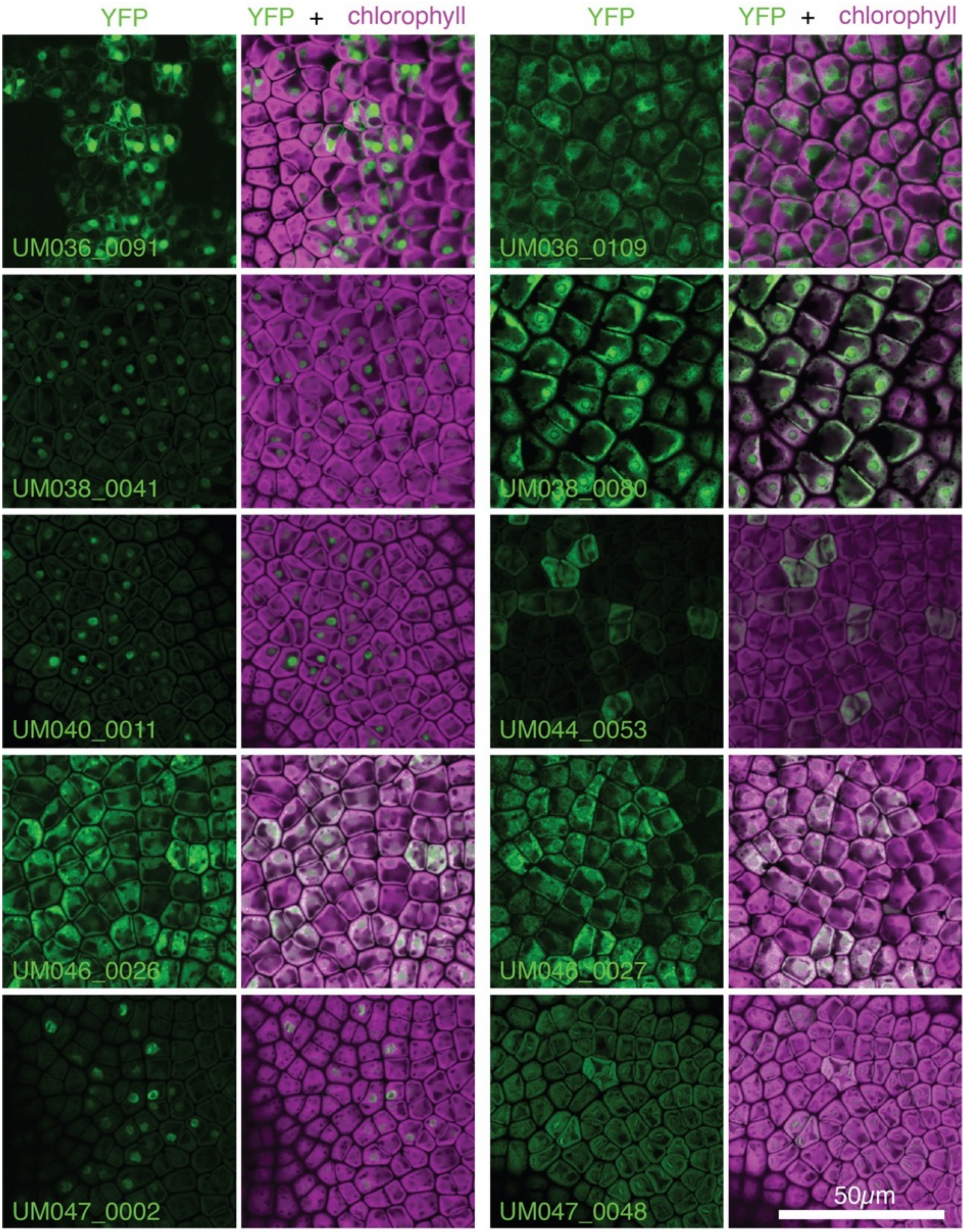


Figure S3 11/16

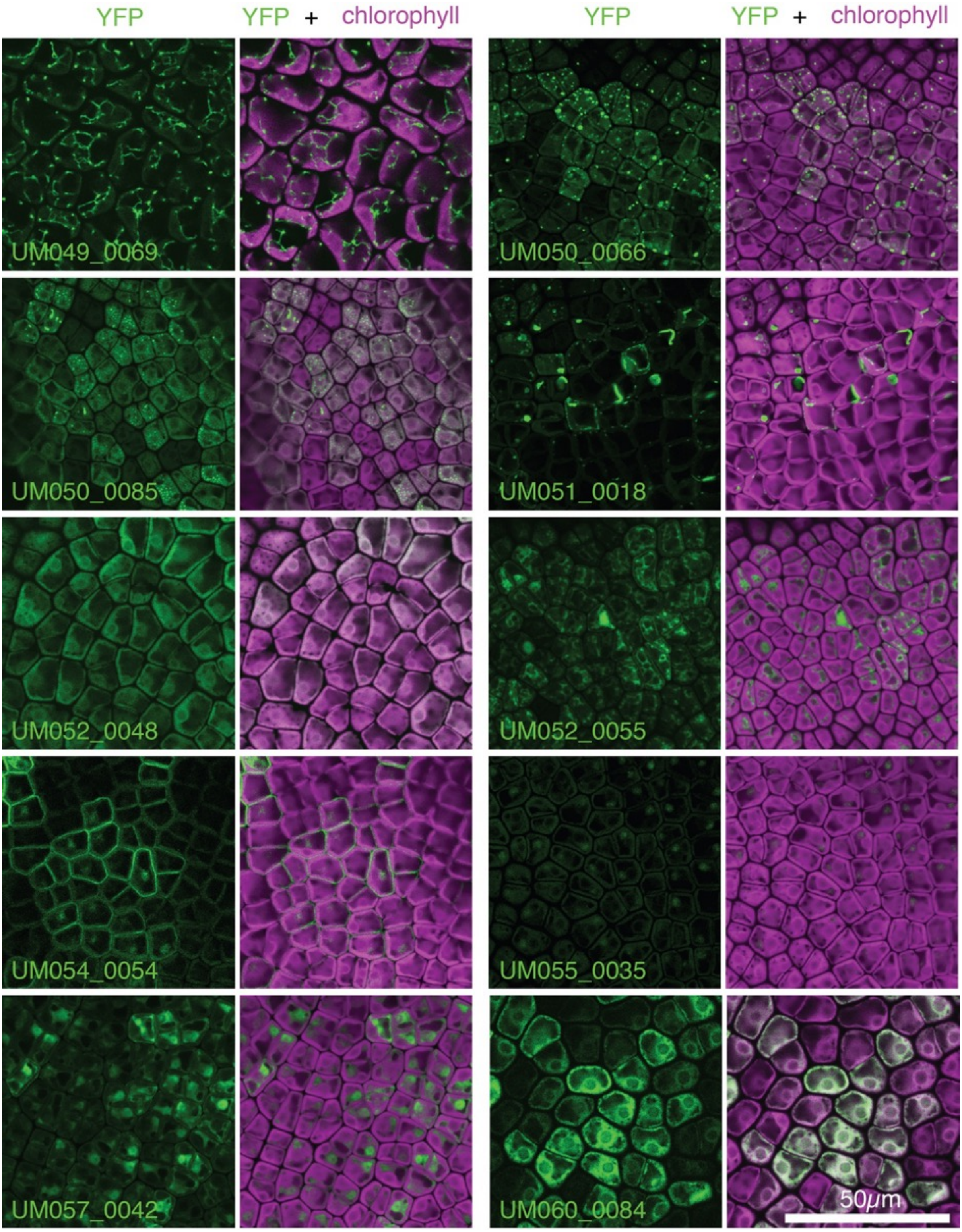


Figure S3 12/16

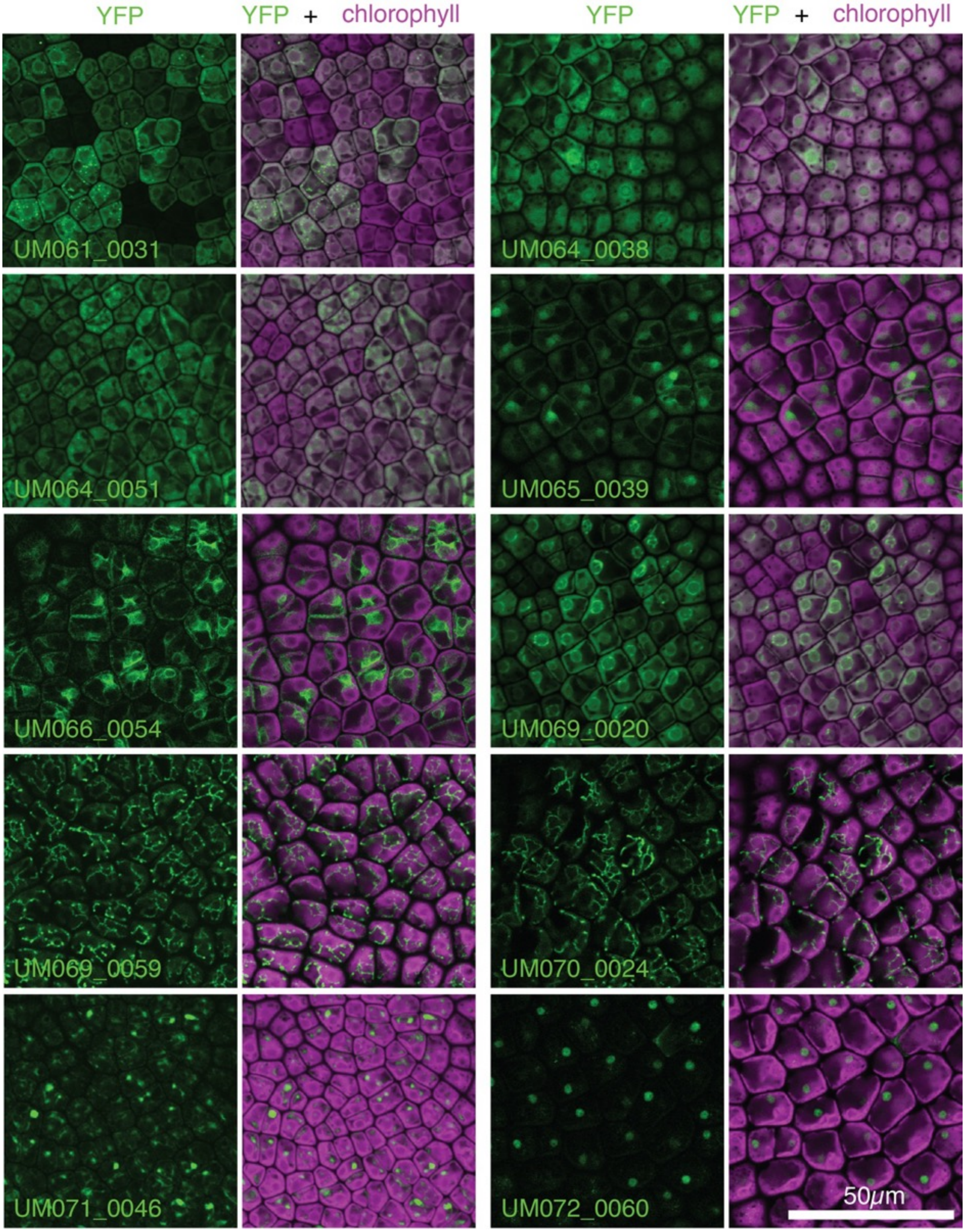

Figure S3 13/16

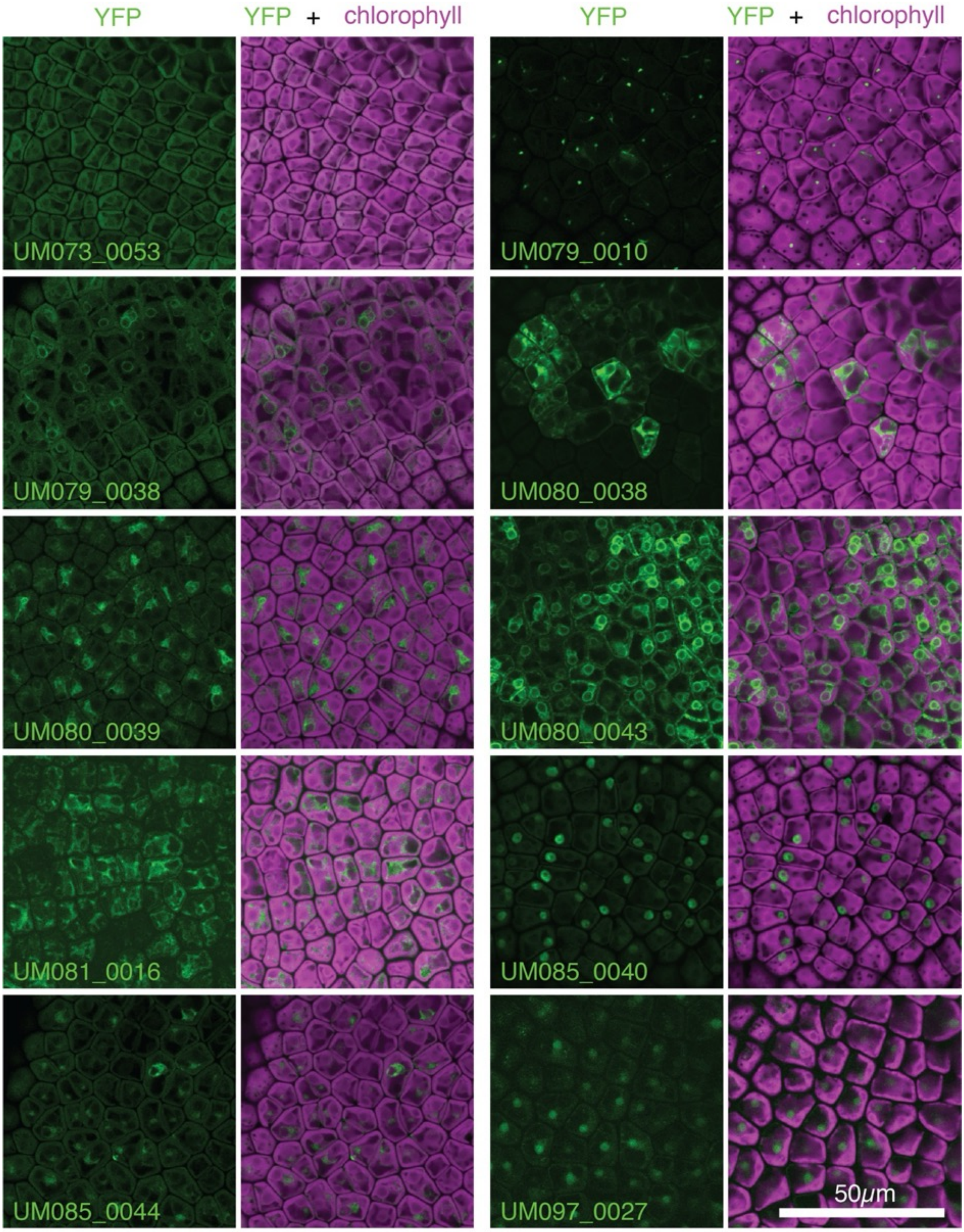


Figure S3 14/16

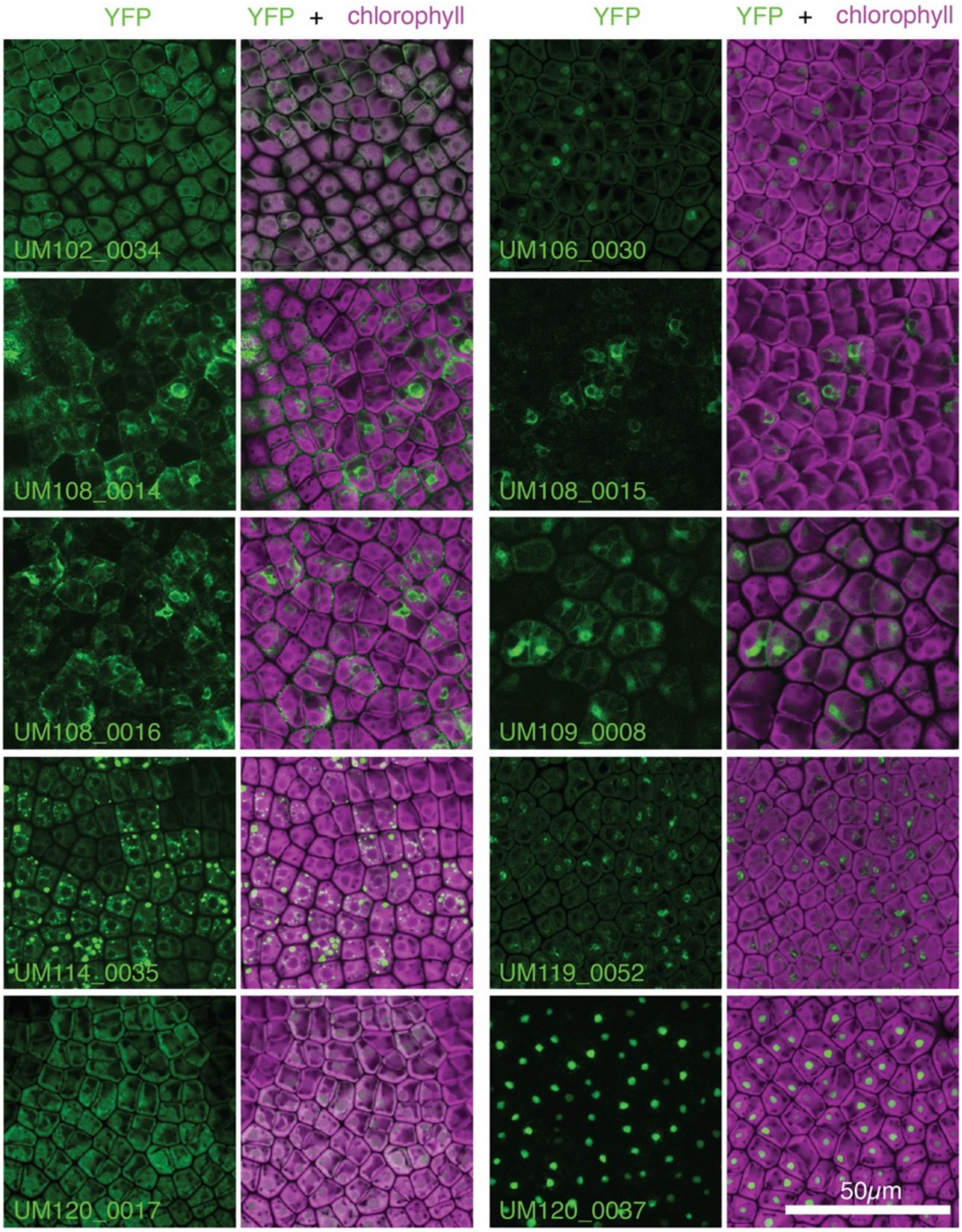

Figure S3 15/16

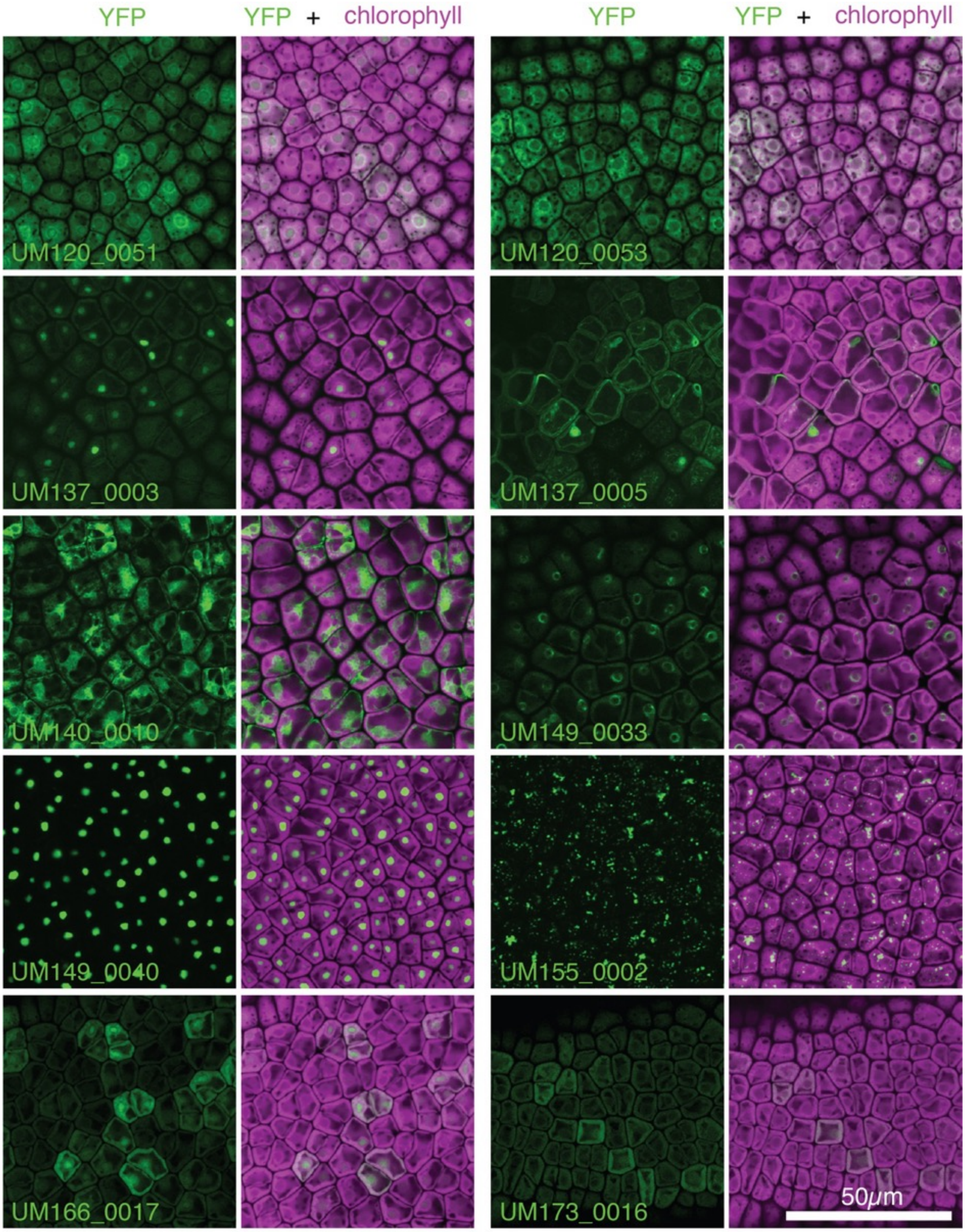


Figure S3 16/16

**Figure S3. Protein localizations of tagged *Ulva* pCCM-related genes.**
For each tagged line a representative individual is selected. Magenta: Chlorophyll autofluorescence; Green: YFP signal of tagged protein. Also see **Table S2.** Scale: 50 μm.

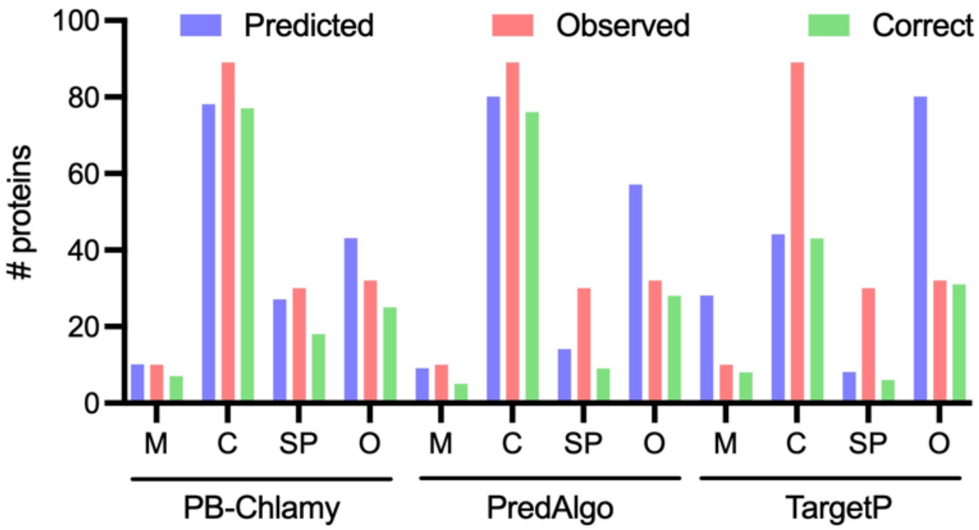


**Figure S4. Comparison of observed and predicted localizations**

Protein localizations predicted by PB-Chlamy, PredAlgo and TargetP, with the number of observed and correctly predicted proteins per category: M: mitochondria; C: chloroplast; SP: secretory pathway; O: other localization. Also see **Table S2**.

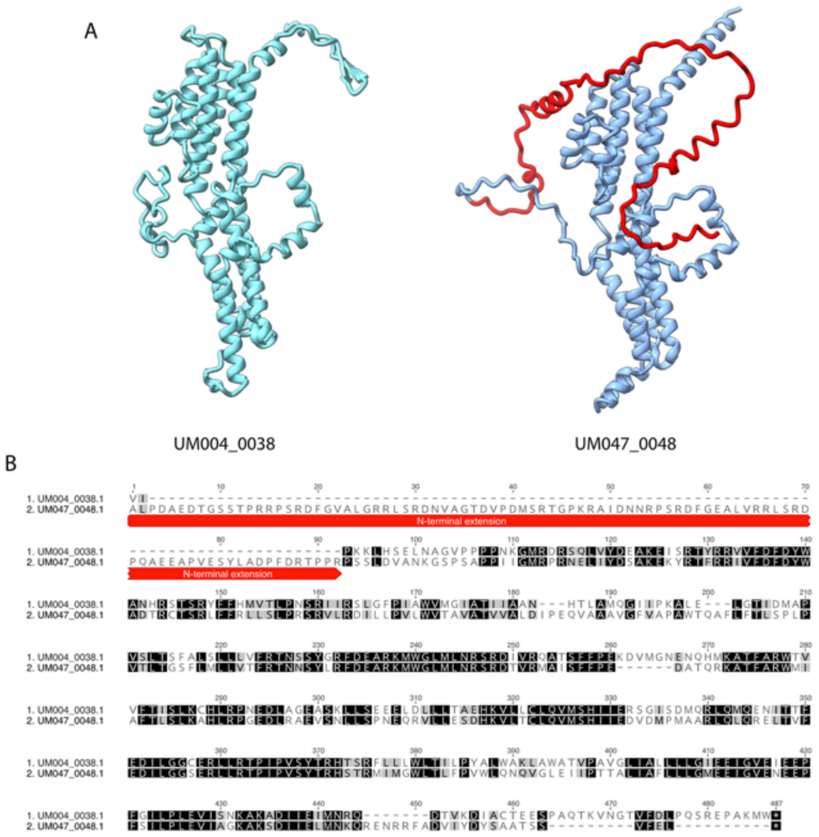


**Figure S5. Identification of *Ulva* BST4 functional analog.**

A. AlphaFold3 models of the pyrenoid-localized Bestrophin monomers. The N-terminal extension of UM047_0048 is shown in red.

B. MAFFT alignment of the predicted mature BST sequences, with the N-terminal extension annotated.

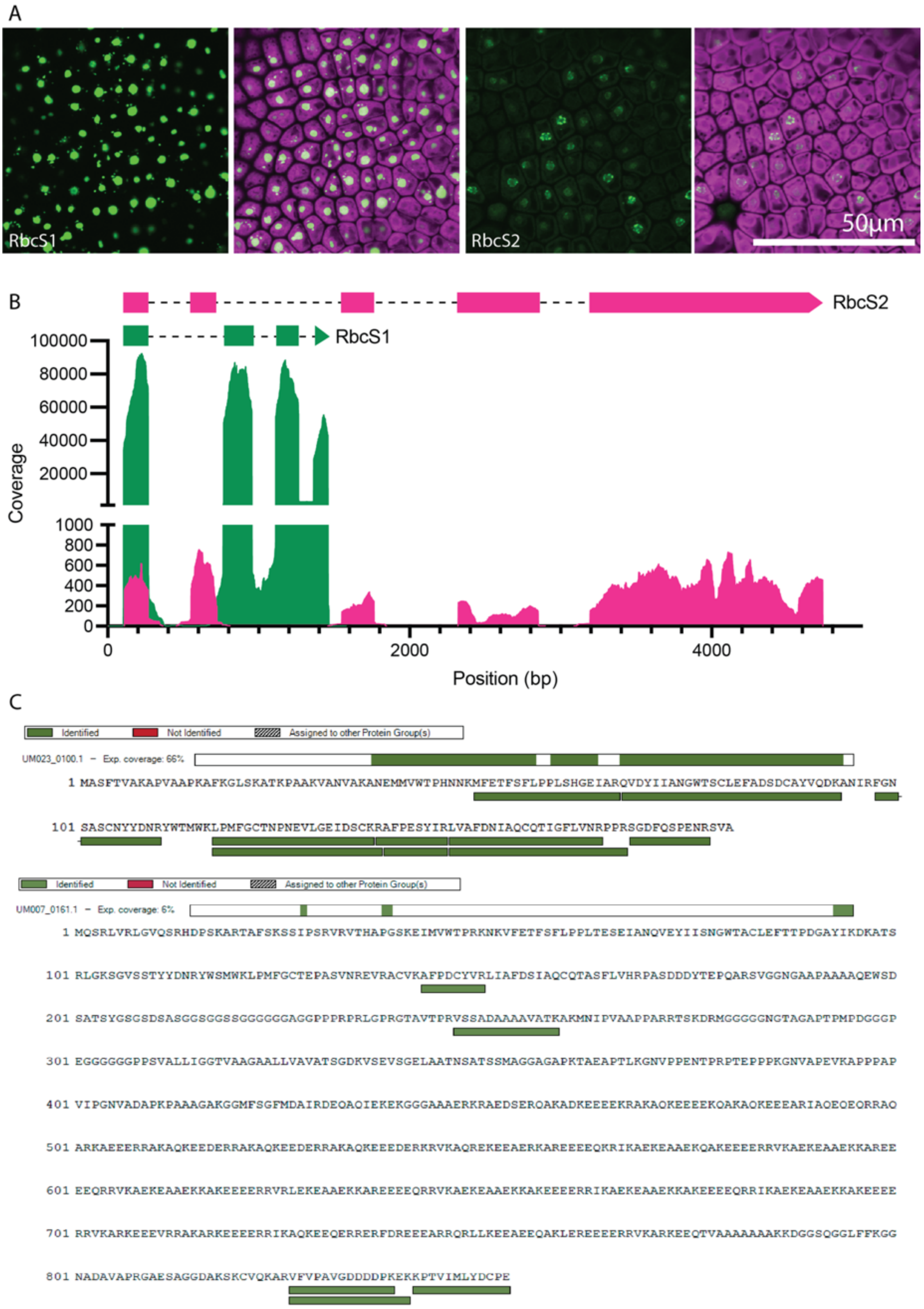


**Figure S6. RbcS2 is lowly expressed in *Ulva*.**

A. Localization of tagged RbcS1 and RbcS2. Magenta: Chlorophyll autofluorescence; Green: YFP signal of tagged protein. Scale: 50 µm.
B. RNA reads mapped to RbcS1 (green) and RbcS2 (magenta) gene model.
C. Peptides mapped to RbcS1 (top) and RbcS2 (bottom).

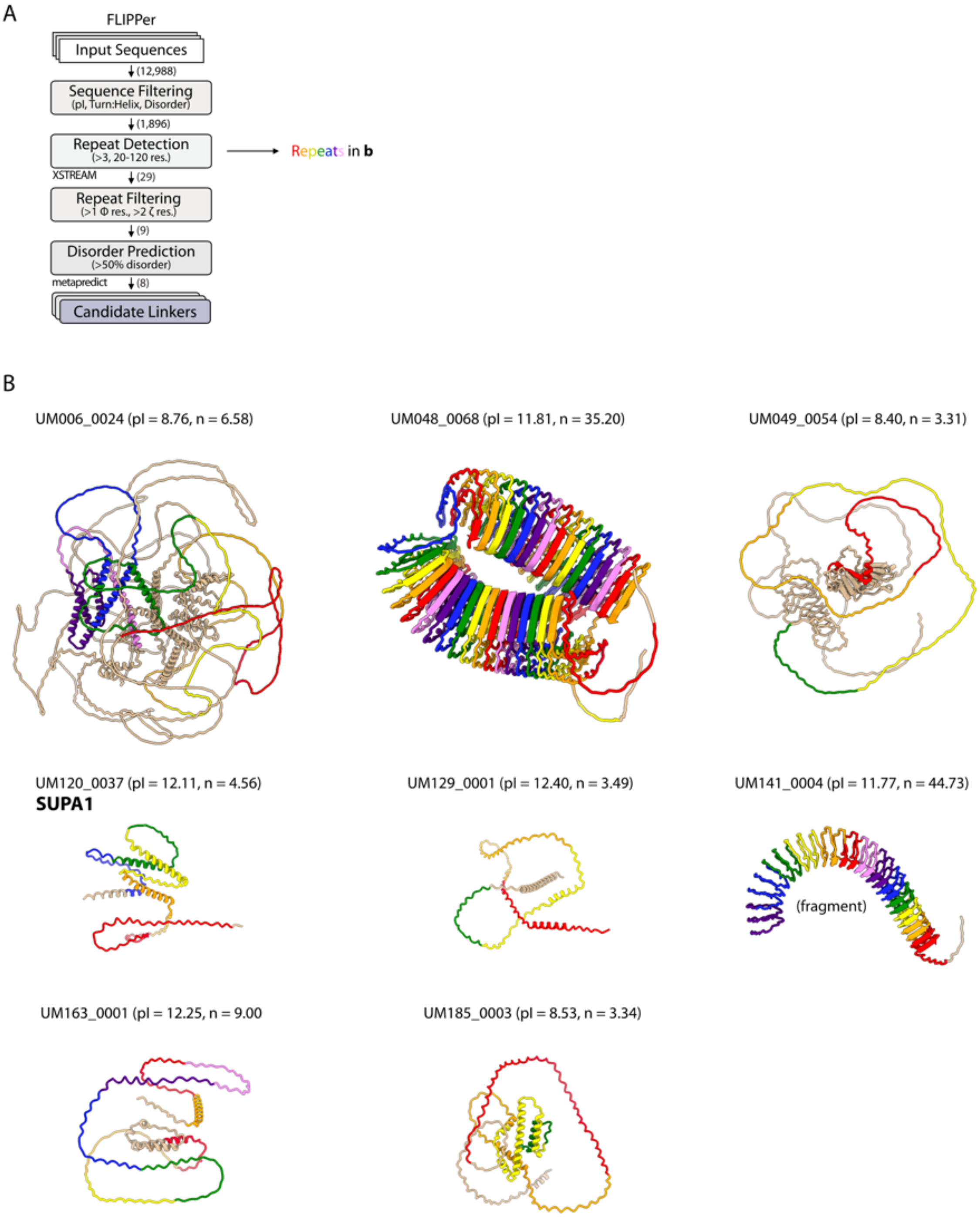


**Fig. S7. FLIPPer analysis of *Ulva* proteins.**

A. Schematic representation of the FLIPPer pipeline, with the programmes used at each stage indicated. The number of sequences remaining after each filtering step of the *Ulva mutabilis* genome is indicated. Acronyms: pI, isoelectric point; res., residue; Φ, hydrophobic; ζ, electrostatic. The repeats detected by XSTREAM are annotated in b.

B. Alphafold 3 models of the 8 candidate proteins identified by FLIPPer. The protein structures are colored according to the repeats identified by XSTREAM from N- to C-termini in the rainbow (red, orange, yellow, green, blue, indigo, violet). The indicated number of repeats is also shown associated with each protein.

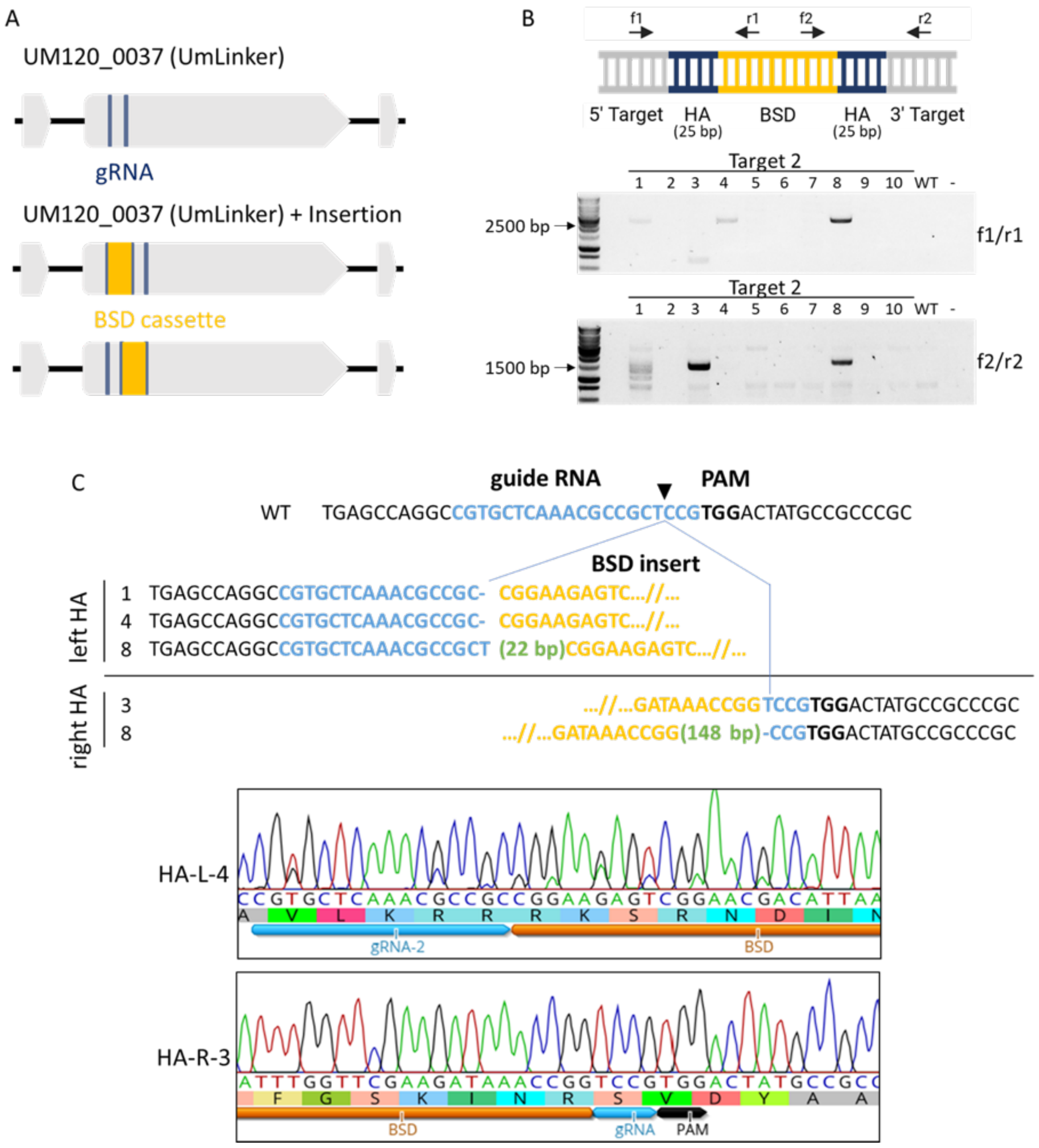


**Figure S8. Genotyping of SUPA1 mutant lines.**

A. Gene model of UM120_0037 (SUPA1) and schematic representation of the expected Blasticidin resistance expression cassette (BSD) knock-in outcomes. Exons in gray, introns in black lines, gRNAs in blue, BSD in yellow.

B. Genotyping PCR analysis of the template insertion in the SUPA1 target 2. Primers amplify the left or right homology arm (HA) (blue), genomic DNA (gray) and the BSD cassette (yellow).

C. Sanger sequencing results showing the correct insertion of the BSD template (yellow) in the expected cut site (black arrow) of either or both the left or right HA, visualized using Geneious software. The events in sample 8 include additional insertions (green) corresponding to duplications of DNA sequences from upstream or downstream of the target site.

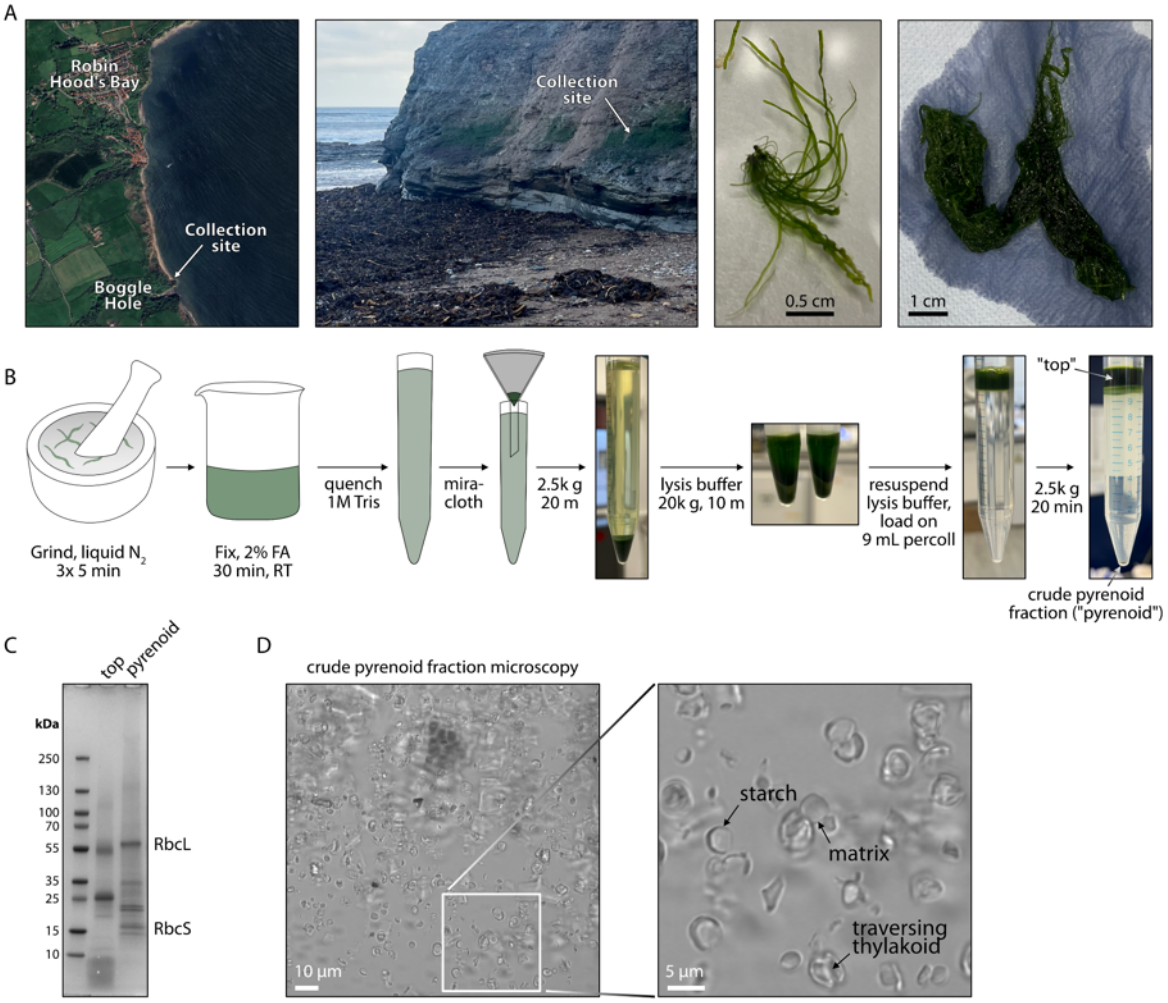


**Figure S9. Pyrenoid enrichment from *Ulva* biomass.**

A. Collection location at Robin Hood's Bay, North Yorkshire, UK. An example individual is shown, alongside the biomass processed for the experiment.
B. Pyrenoid enrichment protocol showing key steps.
C. SDS-PAGE analysis of final crude pyrenoid fraction used for mass spectrometry experiments, as well as "top" fraction from Percoll gradient.
D. Brightfield microscopy of crude pyrenoid fraction. Enlarged image shows presumed sub-pyrenoid structures.

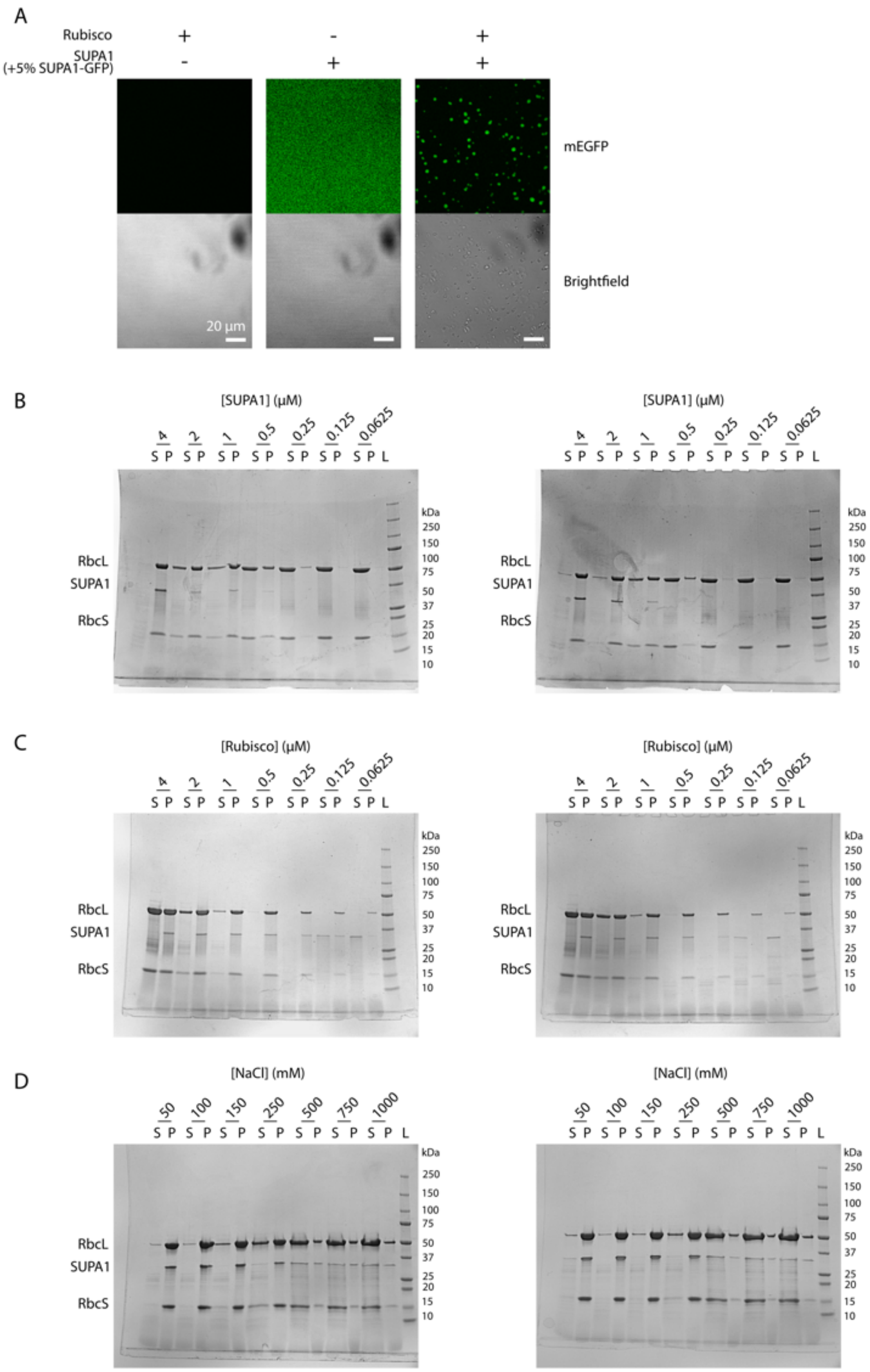


**Figure S10.** ***In vitro*** **characterization of SUPA1-Rubisco condensation.**

A. Confocal microscopy of heterotypic Rubisco phase separation by SUPA1. Rubisco and SUPA1 do not form droplets alone, but form droplets when mixed. 5% SUPA1-GFP was included in SUPA1 samples for visualization. Scale bar = 20 µm.
B. SDS-PAGE gels for fixed 2 µM Rubisco concentration droplet sedimentation assays with titrated concentration of SUPA1. S = supernatant, P = pelleted. Used for quantification presented in Fig. 6B.
C. SDS-PAGE gels for fixed 2 µM SUPA1 droplet sedimentation assays. Used for quantification presented in Fig. 6C.

D. SDS-PAGE gels for droplet sedimentation assays completed at fixed 4 µM SUPA1 to 1 µM Rubisco ratio with increasing NaCl concentration. Used for quantification presented in Fig. 6D.

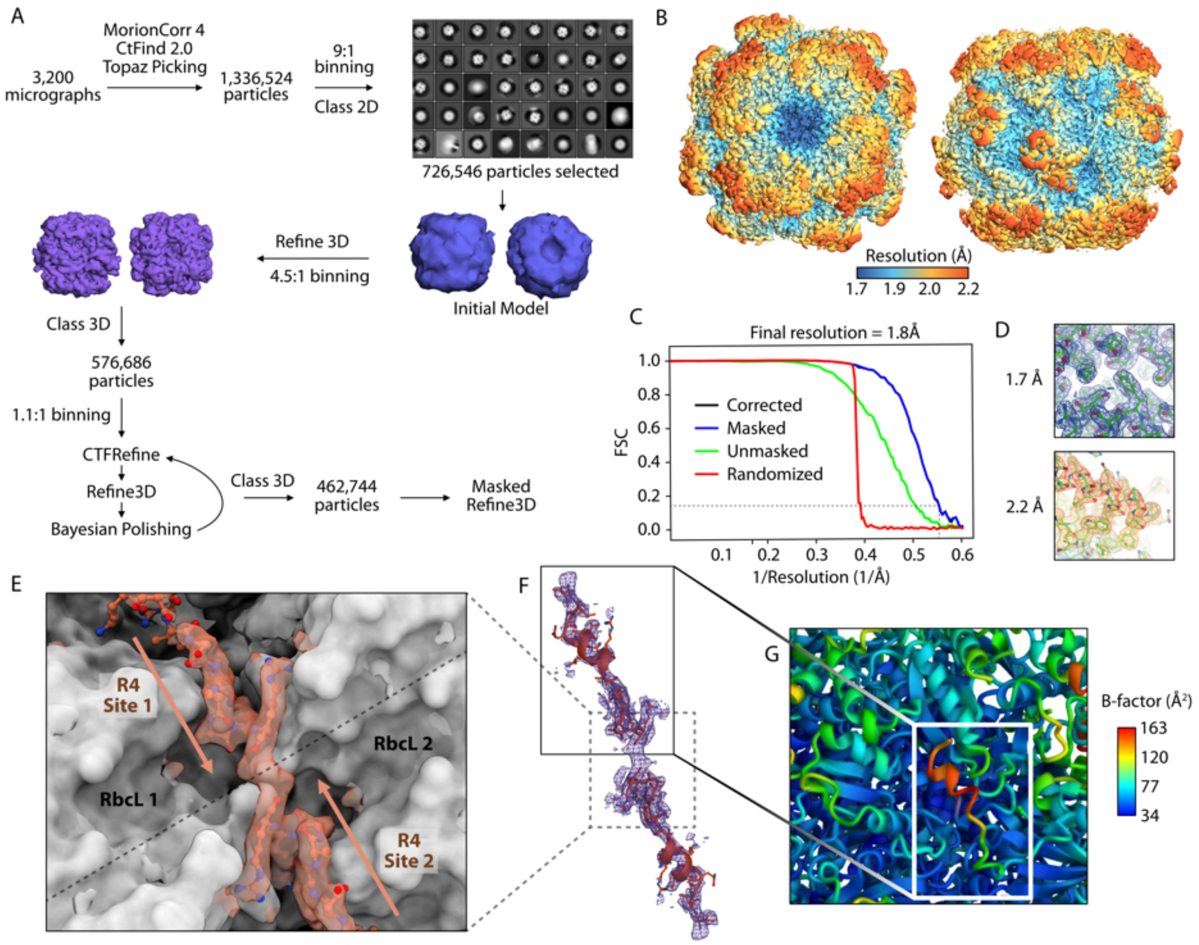


**Figure S11. Single particle cryo-EM data processing and interpretation**

A. Data processing workflow.
B. Density map colored by local resolution.
C. Fourier shell correlation (FSC) curve showing the resolution estimate for the map with FSC cut-off of 0.143 (dashed lines).
D. Example density of the highest and lowest resolution regions of the map.
E. Continuous density across the symmetry interface (dashed line). Models of R4 in site 1 and site 2 reveal steric clashes across the interface.
F. Local density across the two R4 sites.
G. Cartoon representation of a single R4 site colored by estimated B-factor.

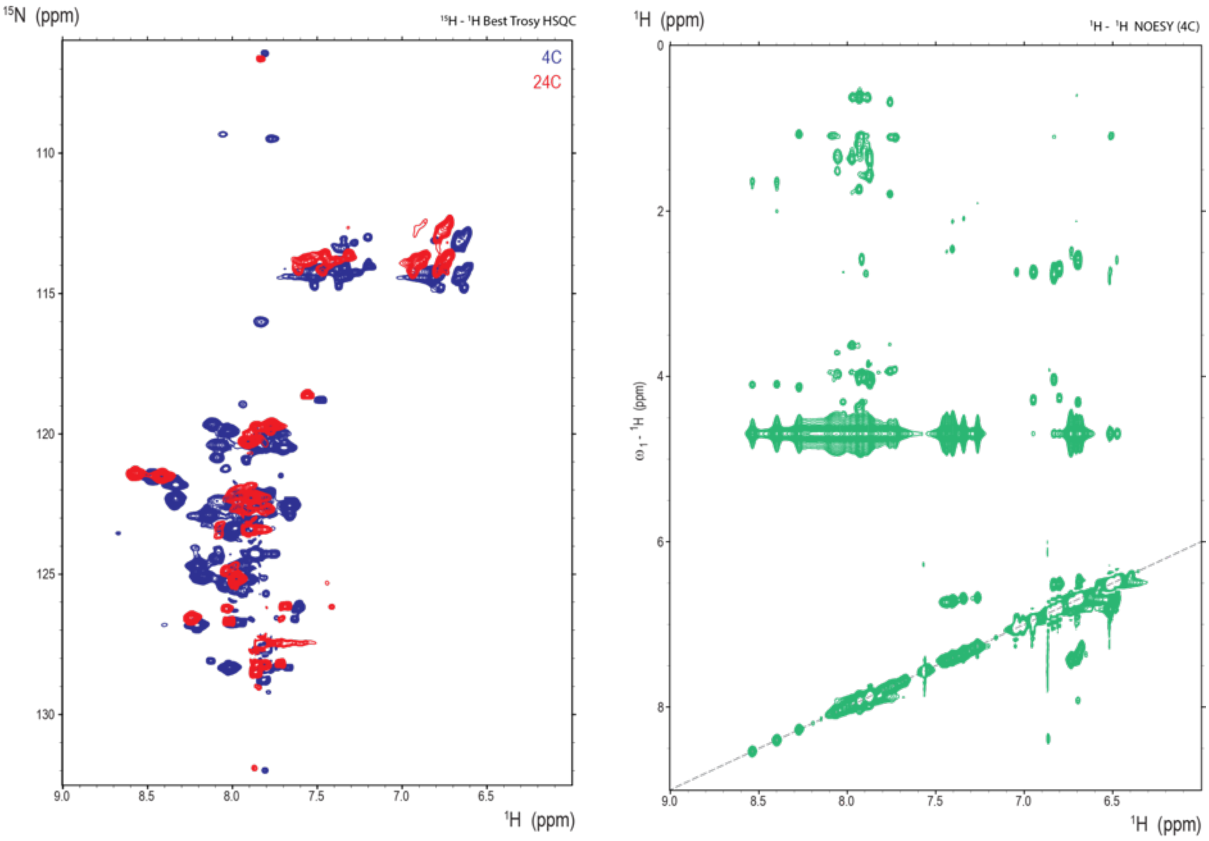


**Figure S12. NMR analysis of the 53-residue RBM linker fragment.**

**(Left)** 2D ($^1$H,$^{15}$N) BEST-TROSY HSQC spectra of the RBM linker fragment recorded at 4 °C (blue) and 24 °C (red). At 4 °C, the spectrum is well defined, with approximately 58 cross-peaks, consistent with the expected number of backbone amides for the linker fragment. The limited dispersion in the proton dimension (~1.5 ppm) is characteristic of an intrinsically disordered protein.

**(Right)** 2D ($^1$H,$^1$H) NOESY spectrum of the linker fragment. No off-diagonal cross-peaks are observed in the amide region (7.5-8.5 ppm, based on Left) that would indicate α-helical structure. In addition, no medium- or long-range NOEs are observed between side-chain proton resonances, further supporting the absence of stable secondary structure.

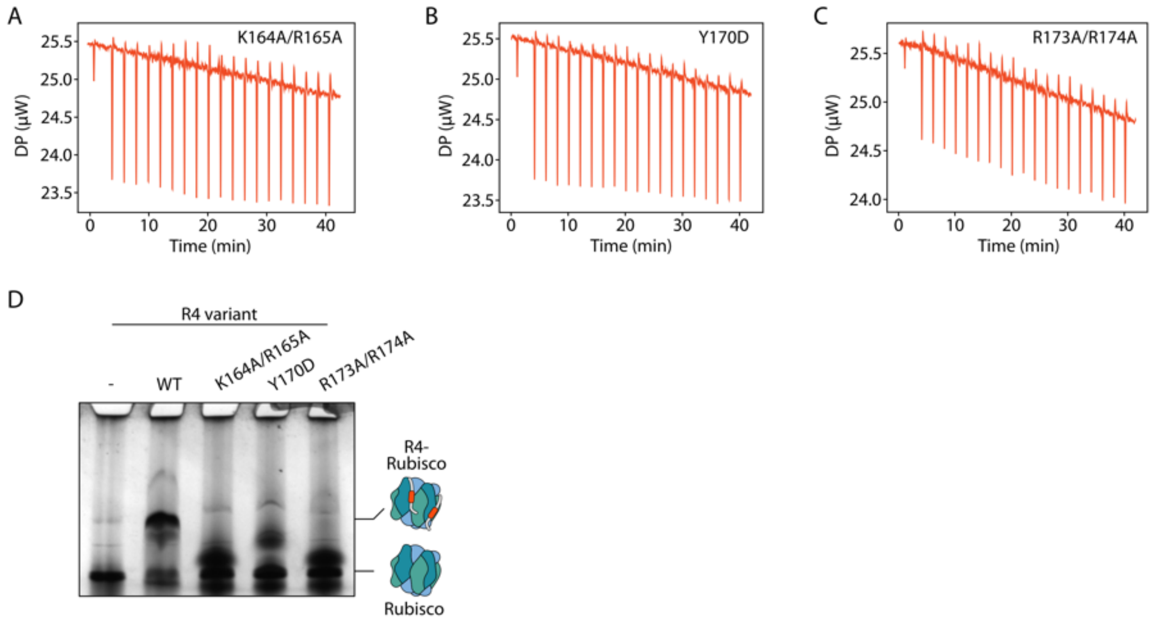


**Figure S13. ITC and native PAGE characterization of SUPA1-R4 binding-deficient mutants.**

(A-C). Raw isothermal titration calorimetry thermograms for R4 K164A/R165A, Y170D and R173A/R174A mutants titrated into Rubisco. No detectable heat exchange above dilution heat is observed for the mutants, precluding meaningful fitting and indicating the absence of measurable interaction under these conditions.

D. Native PAGE characterization of R4 mutant variants.

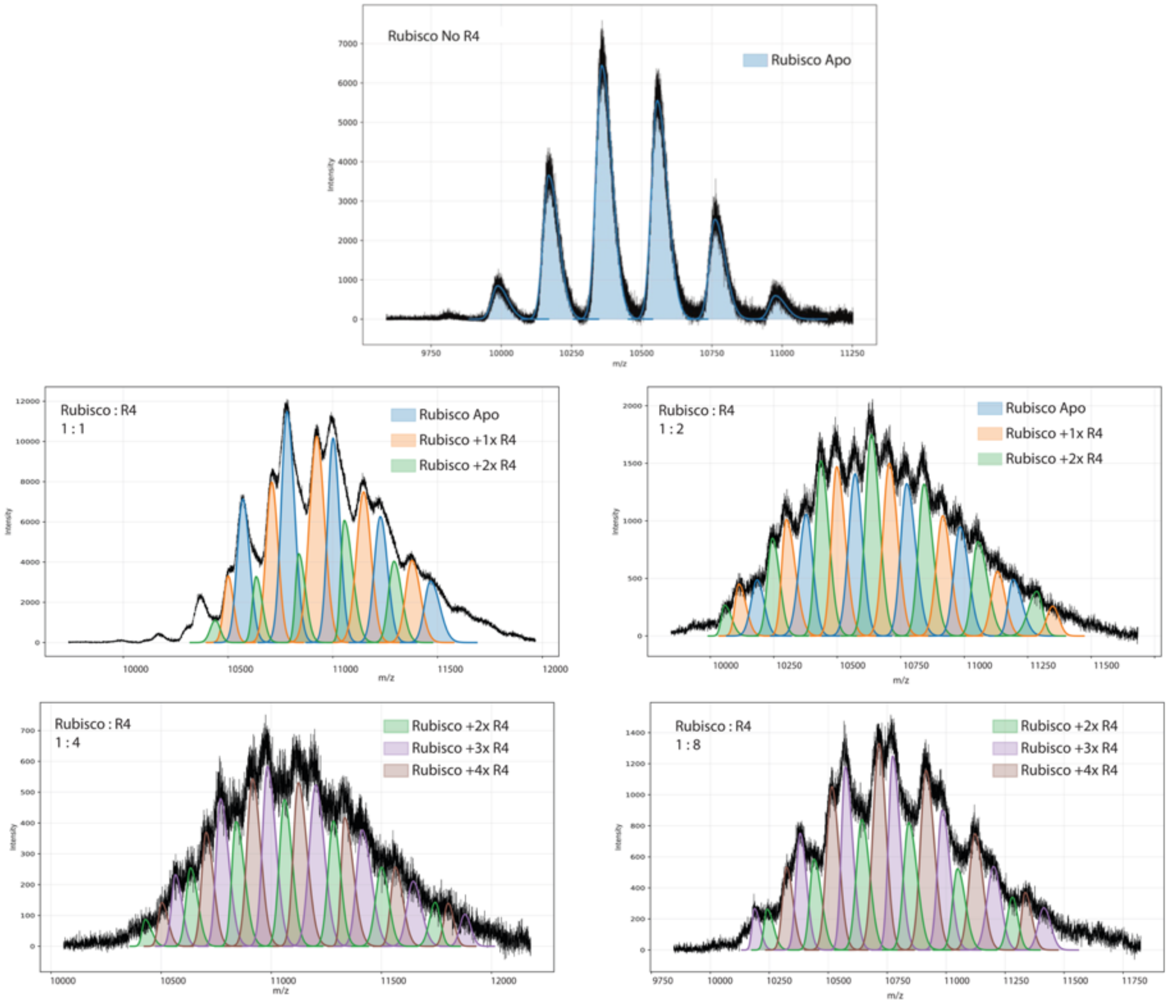


**Figure S14. Raw native mass spectrometry data corresponding to Fig. 6N.**

Native Mass spectra of apo-Rubisco (top). Representative native mass spectra of Rubisco incubated with SUPA1-R4 at Rubisco:R4 molar ratios ranging from 1:1 to 1:8. Peaks corresponding to distinct Rubisco–R4 stoichiometries are indicated, and shaded areas denote the integration ranges used for quantification. These integrated peak intensities were used to determine the species distributions shown in Fig. 6N.

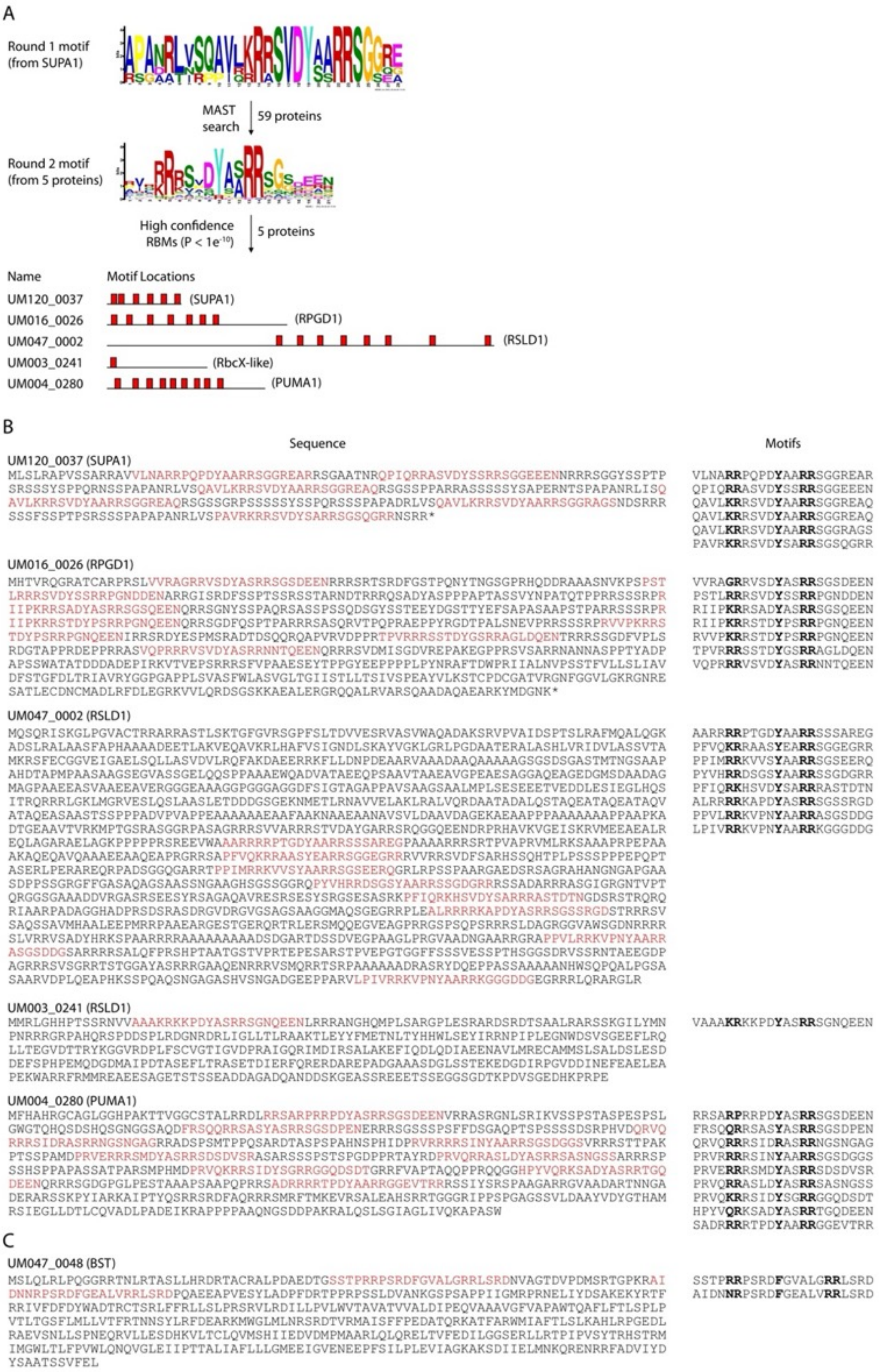


**Figure S15. Identification of RBM-containing proteins.**

A. A motif based on the 6 stickers of SUPA1 was used for a MAST search against the proteome, which enriched the motif. The subsequent motif was used to identify high-confidence sites ($p < 1e^{-10}$).
B. Sequences of Rubisco-binding motif (RBM) containing proteins. Motifs are shown in red in the sequences and aligned adjacent to each sequence. Bold residues are Rubisco-interacting.
C. Sequences of the two partially matching Rubisco-binding motifs of UM047_0048.

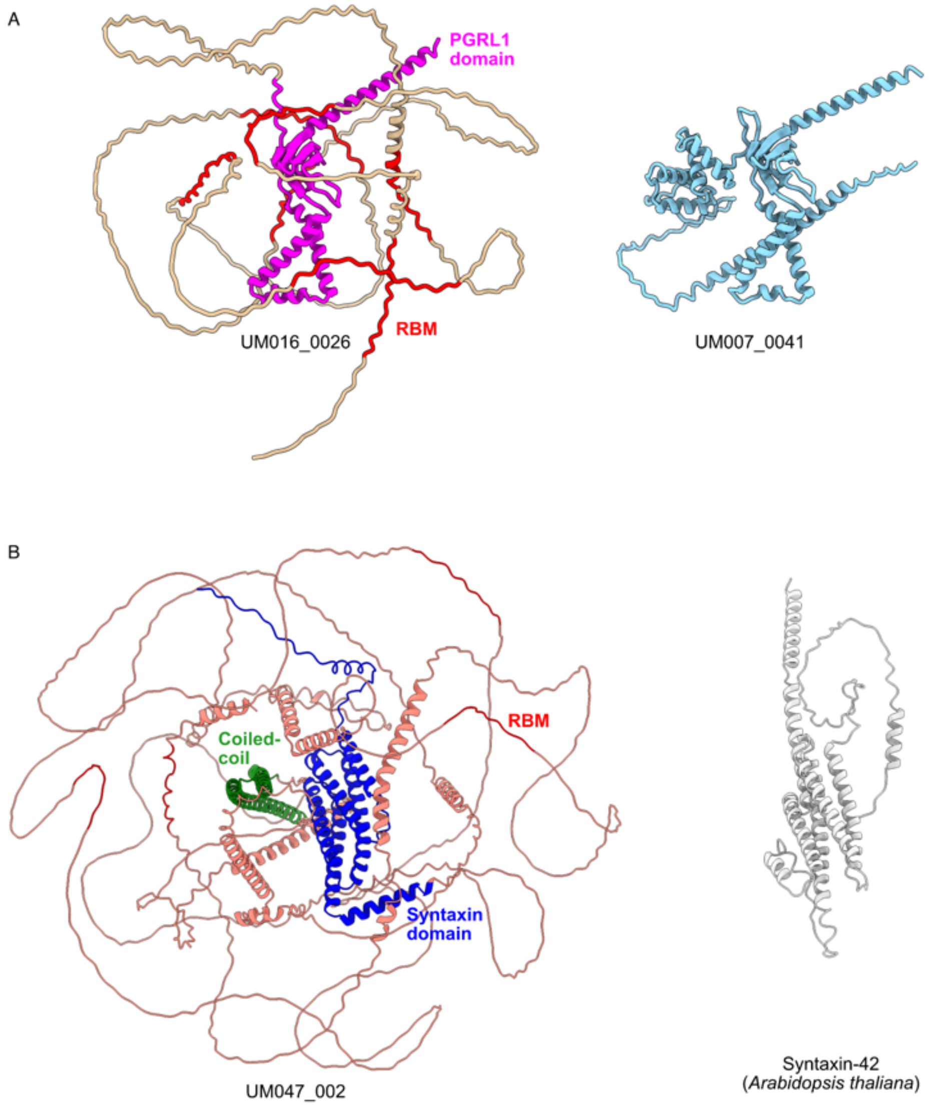


**Figure S16. Putative membrane-associated pyrenoid proteins.**

A. PGRL1 domain (magenta) and RBMs (red) of RPGD1 (UM016_0026) compared to the PGRL1 of *Ulva mutabilis* (UM007_0041).
B. Syntaxin-like (blue), coiled-coil (green) and RBMs of RSLD1 (UM047_002) compared to the structurally homologous syntaxin-42 of *Arabidopsis thaliana*.
